# Optimized reporters for multiplexed detection of transcription factor activity

**DOI:** 10.1101/2024.07.26.605239

**Authors:** Max Trauernicht, Teodora Filipovska, Chaitanya Rastogi, Bas van Steensel

**Affiliations:** Oncode Institute, Division of Gene regulation and Division of Molecular Genetics, Netherlands Cancer Institute, 1066 CX Amsterdam, the Netherlands; Department of Biological Sciences, Columbia University, New York, NY, USA

**Keywords:** MPRA, massively parallel reporter assay, TF, transcription factor, signaling pathways, reporter, specificity, multiplexed TF reporter assay, TF reporter assay

## Abstract

In any given cell type, dozens of transcription factors (TFs) act in concert to control the activity of the genome by binding to specific DNA sequences in regulatory elements. Despite their considerable importance in determining cell identity and their pivotal role in numerous disorders, we currently lack simple tools to directly measure the activity of many TFs in parallel. Massively parallel reporter assays (MPRAs) allow the detection of TF activities in a multiplexed fashion; however, we lack basic understanding to rationally design sensitive reporters for many TFs. Here, we use an MPRA to systematically optimize transcriptional reporters for 86 TFs and evaluate the specificity of all reporters across a wide array of TF perturbation conditions. We thus identified critical TF reporter design features and obtained highly sensitive and specific reporters for 60 TFs, many of which outperform available reporters. The resulting collection of “prime” TF reporters can be used to uncover TF regulatory networks and to illuminate signaling pathways.

**HIGHLIGHTS:**

- Systematic design and optimization of transcriptional reporters for 86 TFs
- Characterization of TF-specific reporter design optimization rules
- Evaluation of reporter TF-specificity across a wide array of TF perturbations
- Identification of a collection of 60 “prime” TF reporters with optimized performance

## INTRODUCTION

Intra- and extracellular signals intricately control the activity of dozens of interwoven signaling pathways, often converging on transcription factors (TFs). TFs respond to these upstream signaling cascades and translate them to orchestrate the regulation of the genome. If we knew the activity of all TFs in any given cell type, we might be able to understand how TFs interpret incoming signals, how they drive the downstream changes in gene expression, and how cascades of TF activities progress over time. However, we currently have no reliable method to directly detect many TF activities in parallel.

A variety of computational approaches have been developed to infer TF activities from genome-wide data such as TF binding data (chromatin immunoprecipitation (ChIP)-sequencing)^1^, chromatin accessibility maps (assay for transposase-accessible chromatin (ATAC)-sequencing)^2,3,4^ TF or target gene transcript abundance data (RNA-sequencing)^5,6–8^ or a combination of these methods.^9–11^ While these methods provide convenient tools to compute TF activities from well-established genomics assays, they do not *directly* measure the transcriptional activity of TFs and might therefore lack precision. For example, it is known that maps of TF binding poorly reflect TF activity;^12,13^ ATAC-seq detects open chromatin regions which are not necessarily predictive of transcription activity;^14^ and inferring TF activity from mRNA-seq data requires assumptions regarding the distance over which each TF may be able to control gene activity.

Traditional reporter assays, employing fluorescent or luminescent proteins expressed by TF response sequences, offer direct means to measure TF activities.^15–25^ These assays have been used for decades and detect TF activity with great sensitivity. However, conventional reporter assays do not allow to detect multiple TFs at once. A previous study circumvented this limitation and measured 58 TF activities in parallel from previously published TF reporters by utilizing RNA barcodes as reporters.^26^ This study also showed that TF reporter measurements can be more accurate than RNA-seq-inferred TF activities for a subset of TFs. Thus, directly measuring TF activities in a high-throughput fashion using barcoded reporters offers a direct and precise alternative to computational inference approaches.

Despite the advantages of multiplexed TF reporter assays, there are still several challenges in achieving accurate high-throughput TF activity detection. First, reporters are only available for a limited number of TFs. Expanding the collection of TF reporters will be crucial to make multiplexed reporter assays more scalable. Second, most of the published TF reporters rely on either (i) TF response elements found in the genome,^17,18,22,23^ which might lack specificity to the intended TF due to the presence of other TF binding sites (TFBSs), or (ii) poorly optimized synthetic TF response sequences,^15,16,27^ which could be suboptimal in terms of sensitivity and specificity. Hence, it is necessary to optimize TF reporters to obtain more reliable activity measurements. Finally, it is known that TFs within the same TF family, especially TF paralogs, can have highly similar DNA-binding domains and thus also TFBSs, which complicates the design of reporters that are specific for a single TF.

Here, we report the generation of highly optimized reporters for a large collection of TFs. Towards this goal we made use of massively parallel reporter assays (MPRAs) with a systematically designed library of 5,530 different reporter designs for 86 TFs, including TFs that respond to diverse signaling pathways and a variety of cell type-specific TFs. For each TF, we optimized the design of the reporter by varying the spacer sequences and spacer length between TFBSs, the distance to the core promoter and the core promoter itself. We evaluated the specificity and sensitivity of the generated TF reporters by probing the library across nine cell lines and almost 100 TF perturbation conditions. Detailed analysis of this rich dataset provided insights into the rules that determine the sensitivity and specificity of reporters for each TF, and yielded a collection of “prime” reporters for 60 TFs, for many of which no reporters were available yet. Our synthetic prime reporters outperform published reporters in >80% of all comparisons. We demonstrate the utility of the identified prime reporter set by detecting signaling pathway interdependencies upon pluripotency-challenging perturbations in mouse embryonic stem cells (mESCs).

## RESULTS

### Systematic probing of a TF reporter library

#### Selection of TFs

A main challenge in the design of specific TF reporters is the similarity between binding motifs of TFs. Therefore, to select TFs for which the generation of TF-specific reporters would be feasible, we manually examined all human TFs and reviewed their (i) TF motif quality (i.e., motif length and information content), (ii) the number of TFs with a similar motif, (iii) expression pattern across cell types, and (iv) stimulation and perturbation opportunities. Based on these criteria we selected a list of 86 TFs (**Table S1**). For each TF, we selected the best motif according to a previous motif curation.^28^ We also included several heterodimeric motifs (e.g., POU5F1::SOX2), for which we carefully reviewed available motifs. Most of the selected TFs have unique motifs (i.e., no other TF has a similar motif, **Figure S1A**), and cover a large diversity of the human TF motif landscape (**Figure 1A**). The selected 86 TFs include most well-known TFs downstream of generic signaling pathways such as MAPK, PI3K/AKT, TGFβ, WNT, and JAK-STAT, as well as a diversity of nuclear receptors and tissue-specific and pluripotency-specific TFs (**Table 1, Table S1**).

**Figure 1:**
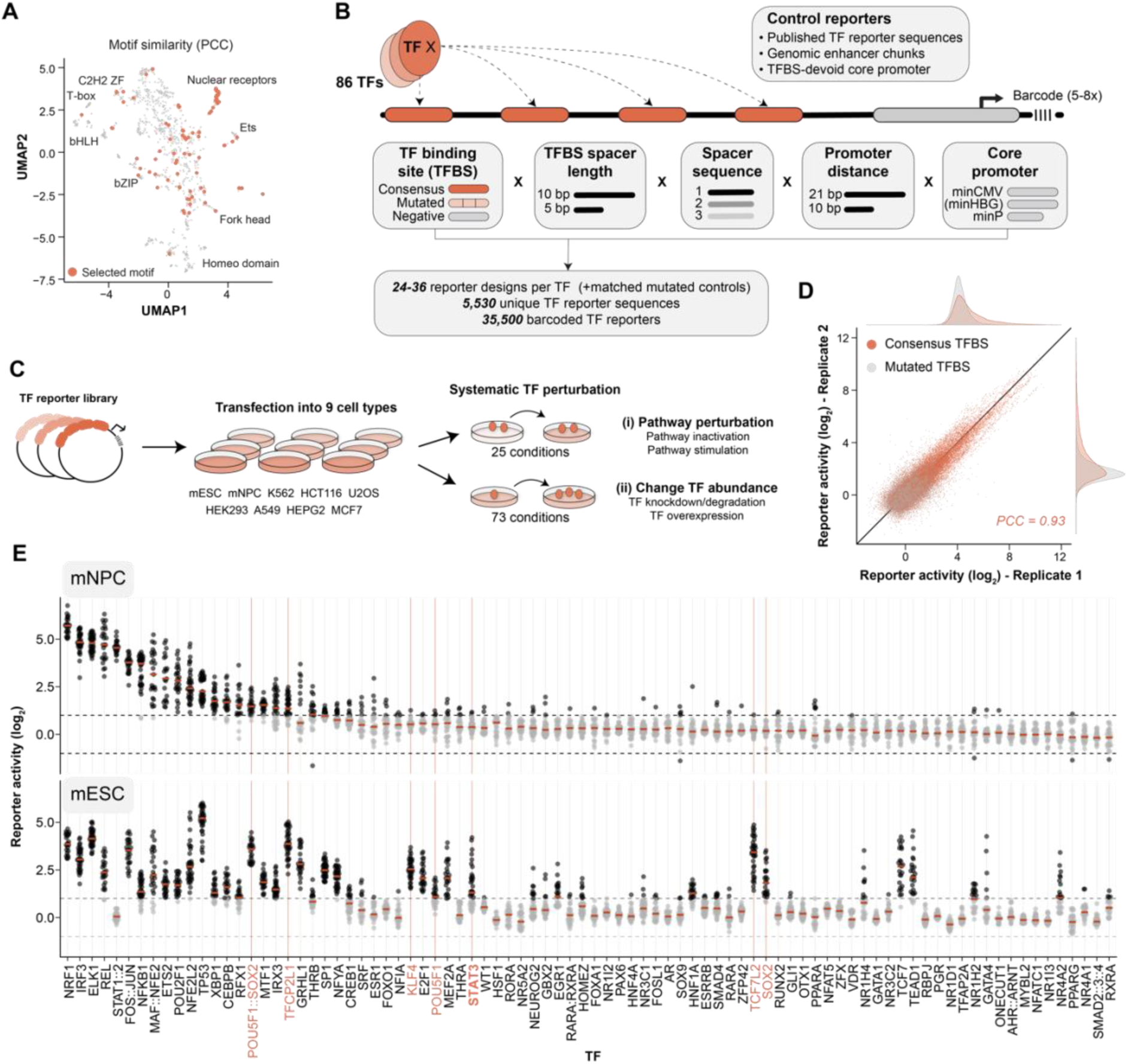
Systematic design and probing of TF reporters. (**A**) UMAP visualization of motif similarities (Pearson correlation coefficient (PCC)) of all human TFs (n = 1,244). The selected 86 motifs are highlighted in red. Local clusters of TF families are annotated. (**B**) Design of the TF reporter library. Four copies of the TFBS are placed around variable spacer sequences and spacer lengths upstream of a core promoter and a unique barcode. The arrow in the core promoter indicates the transcription start site. (**C**) Experimental layout. The TF reporter plasmid library was probed in nine distinct cell lines and in 98 TF perturbation conditions. (**D**) Correlation of all reporter activities (i.e., activation compared to TF-neg reporters) measured in biological replicate 1 compared to replicate 2 of reporters with mutated TFBSs (grey) and consensus TFBSs (red). Reporter activities in all nine cell lines are displayed together. (**E**) Activities of individual reporter designs per TF in mNPCs and mESCs. Highlighted in orange are mESC-specific TFs. Red line indicates median activity per TF.

**Table 1.** Overview of the selected TFs and their primary associated cellular functions. Note that some TFs might have multiple functions. TFs for which only published reporters were included are displayed in parentheses.

| TF | Main cellular function |
| --- | --- |
| AHR::ARNT, NR1I2, NR1I3 | Xenobiotic stress response |
| AR | Testosterone response |
| CEBPB, NFKB1, NR4A1, NR4A2, FOS::JUN, (ATF2) | Inflammation |
| CREB1 | Cyclic AMP |
| E2F1, MYBL2, TP53, (CEBPA) | Cell cycle |
| EGR1, ELK1, ETS2, SRF | MAPK |
| ESR1 | Estrogen response |
| ESRRB, KLF4, POU5F1, SOX2, ZFP42, ZFX | Pluripotency |
| FOXA1, FOSL1 | Cell identity, development |
| FOXO1 | PI3K/AKT |
| GATA1 | Erythroid development |
| GATA4 | Cardiac development |
| GBX2, HOMEZ, IRX3, NEUROG2, NFIA, OTX1, RFX1 | Neural development |
| GLI1 | Hedgehog |
| GRHL1 | Epithelial development |
| HNF1A, HNF4A, ONECUT1 | Hepatic gene activation |
| HSF1 | Heat shock response |
| IRF3, STAT1::2, (IRF1), (STAT1) | Interferon, immune response |
| MAF::NFE2, NFE2L2, NRF1 | Oxidative stress response |
| MEF2A | Myocyte development |
| MTF1 | Metal response |
| NFAT5, NFATc1 | Osmotic stress response |
| NFYA, SP1 | Constitutive activator |
| NR1D1, (CLOCK) | Circadian rhythm |
| NR1H2, PPARA, PPARG, (TFEB), (SREBF1) | Lipid metabolism |
| NR1H4, NR5A2 | Bile acid response |
| NR3C1 | Glucocorticoid response |
| NR3C2 | Mineralocorticoid response |
| PAX6 | Neural & pancreatic development |
| PGR | Progesterone response |
| POU2F1, RORA, TFAP2A, WT1, (MYC), (GATA3) | Various |
| RARA, RXRA | Retinoic acid response |
| RBPJ | Notch signaling |
| RUNX2, SOX9 | Osteoblast development |
| SMAD2::3::4, SMAD4 | TGFβ signaling |
| STAT3, (STAT4), (STAT6) | JAK-STAT signaling |
| TCF7, TCF7L2 | WNT signaling |
| TEAD1 | Hippo signaling |
| THRA, THRB | Thyroid hormone response |
| VDR | Vitamin D3 response |
| XBP1, (ATF4), (ATF6) | Unfolded protein response |
| (HIF1A) | Hypoxia response |

#### Library design

We then generated a library consisting of synthetic TF reporters for the selected 86 TFs. For each TF, we generated a consensus TFBS by choosing the most conserved base at each position of its motif. We also included two sets of negative control TFBSs. First, for each TF we generated a matched mutated TFBS in which two to four conserved bases of the TFBSs were modified (**Table S1**). Second, we designed three distinct 11 bp random sequences that are devoid of any known activator TFBS (TF-neg) that served as generic negative controls and were used for normalization. To generate synthetic TF reporter sequences, we placed four copies of the TFBSs in front of a core promoter that drives the transcription of a unique 13-bp barcode sequence and a GFP open reading frame (**Figure 1B**). We chose to use four copies of TFBSs, because this number was shown to yield optimal activation for many TFs.^29–32^ We then systematically varied several design parameters for each TFBS (**Figure 1B**). First, we designed three spacer sequences around the four TFBSs of either 5 or 10 bp (i.e., TFBS1-spacer1-TFBS2-spacer2-TFBS3-spacer3-TFBS4). These spacer sequences were computationally designed to minimize occurrences of other TFBSs, even in the junctions between the spacer sequences and the TFBSs (**Figure S1B**). For each spacer length (5 and 10 bp), we then selected three distinct sets of spacer sequences. Second, we coupled the TFBSs to three different core promoters (minP (derived from pGL4 (Promega, Madison, WI, USA)), minCMV,^33^ and for some TFs also minHBG^34^). Third, we placed the core promoter at either 10 or 21 bp from the nearest TFBS. Together, the combination of these design parameters yielded 36 reporter designs for TFs with minHBG, and 24 for TFs without. Additionally, for comparison we also included previously established and published reporter sequences for 62 TFs from three different public sources (see Methods; **Table S1**).^26,27^ A set of 120 enhancer fragments from the mouse *Klf2* locus (previously shown to be active in MPRAs in mESCs),^35^ and 86 reporters with a TFBS-devoid core promoter (one for each TF) were included as positive and negative controls, respectively (see Methods). Together, this yielded a collection of in total 5,530 unique reporter sequences. Finally, each of these sequences was coupled to 5-8 distinct barcodes to minimize biases caused by individual barcodes, yielding a library of 35,500 barcoded reporters (**Table S2**).

#### Systematic testing of TF activities

Among the 86 included TFs are many tissue-specific TFs. We therefore probed the reporter library in nine different cell lines from distinct tissues (**Figure 1C**). Since TF binding specificities are highly conserved between human and mouse,^36,37^ we tested the library in cell lines derived from both human (n = 7) and mouse (n = 2). Furthermore, we extensively perturbed TF activities by (i) activating or inhibiting upstream signaling pathways (n = 25), or (ii) changing the TF abundance by overexpressing, knocking down or degrading individual TFs (n = 73). We thus queried all 5,530 reporters across 98 TF-perturbation conditions (**Figure 1C**).

#### Data overview

For each tested condition or cell line, we first normalized the barcode counts in the mRNA to the barcode counts in the input plasmid DNA. Activities were then computed from the plasmid DNA-normalized counts by calculating the induction over the median counts of the collection of the TF-neg reporters. This was done separately per core promoter. Reporter activities between individual barcodes correlated highly (Pearson’s correlation coefficient (PCC) range 0.84 – 0.87, **Figure S1C**) and were averaged. We probed the reporter library per cell line in at least three (HEK293, K562) and up to 11 (mESCs) biological replicates, yielding per cell line an average PCC between replicates of 0.77 - 0.94 (**Figure S1D**). For downstream analyses we averaged the reporter activities of the replicates. As expected, reporters with consensus TFBSs were more active than reporters with minimal mutations in the TFBS in all nine tested cell lines (**Figure 1D, S1F**). Furthermore, the synthetic TF reporters reached activities as high as the genomic enhancer element reporters, showing that four copies of the same TFBS are as potent as highly active native enhancer elements of approximately the same length (**Figure S1F**).

#### Reporter activities depend on cell type

We first characterized activities for all TFs and their 24-36 reporter designs across the nine probed cell lines. We found that known generic TFs displayed activities in all cell lines (e.g., ELK1, FOS::JUN), while known cell type-specific TFs were predominantly detected in a subset of cell lines (e.g., HNF1A or HNF4A in HEPG2), and some were not active in any cell type (e.g., VDR, see below; **Figure S2**). Next, to explore these cell type-specific activities in more detail, we focused on two different cell lines: mESCs and mESC-derived neural precursor cells (mNPCs). As expected, the reporter activities differed for many TFs between the two different cell lines (**Figure 1E**). TFs that displayed substantially higher activity in mESCs compared to mNPCs included POU5F1::SOX2, TFCP2L1, STAT3, KLF4, SOX2, and TCF7 (**Figure 1E**, highlighted in orange), which are known activating TFs of the mESC pluripotency network.^38,39^

#### Reporter activities strongly vary between designs

Importantly, for several TFs the reporter designs showed substantial differences in activity, despite having identical TFBS. For instance, in mESCs some STAT3 reporter designs were as inactive as the TF-neg control reporters, while others were up to 25-fold more active than those controls (**Figure 1E**, highlighted in bold and orange). This indicates that the design of the reporter can have substantial effects on its activity.

### Understanding TF-specific reporter design rules

#### Reporter design explains variance in reporter activities

To investigate the relation between reporter design and reporter activity, we fitted for each TF a log-linear model using the reporter design features (core promoter identity, promoter distance, spacer sequence and length) as categorical input variables (**Figure 2A**). This analysis enabled us to extract which features contribute to the variation in reporter activity. For example, for STAT3 the model accurately reflected the measured reporter activities (adjusted R^2^ = 0.95) (**Figure 2B**), and indicated that spacer length and spacer sequence were crucial to achieve high transcriptional activity, while promoter identity contributed moderately, and promoter distance was largely irrelevant (**Figure 2C**). This suggested that STAT3 is more active with TFBSs spaced by 10 bp, and that the spacer sequence can strongly impact reporter activity, even though the spacers were designed not to contain any known TF motif.

**Figure 2:**
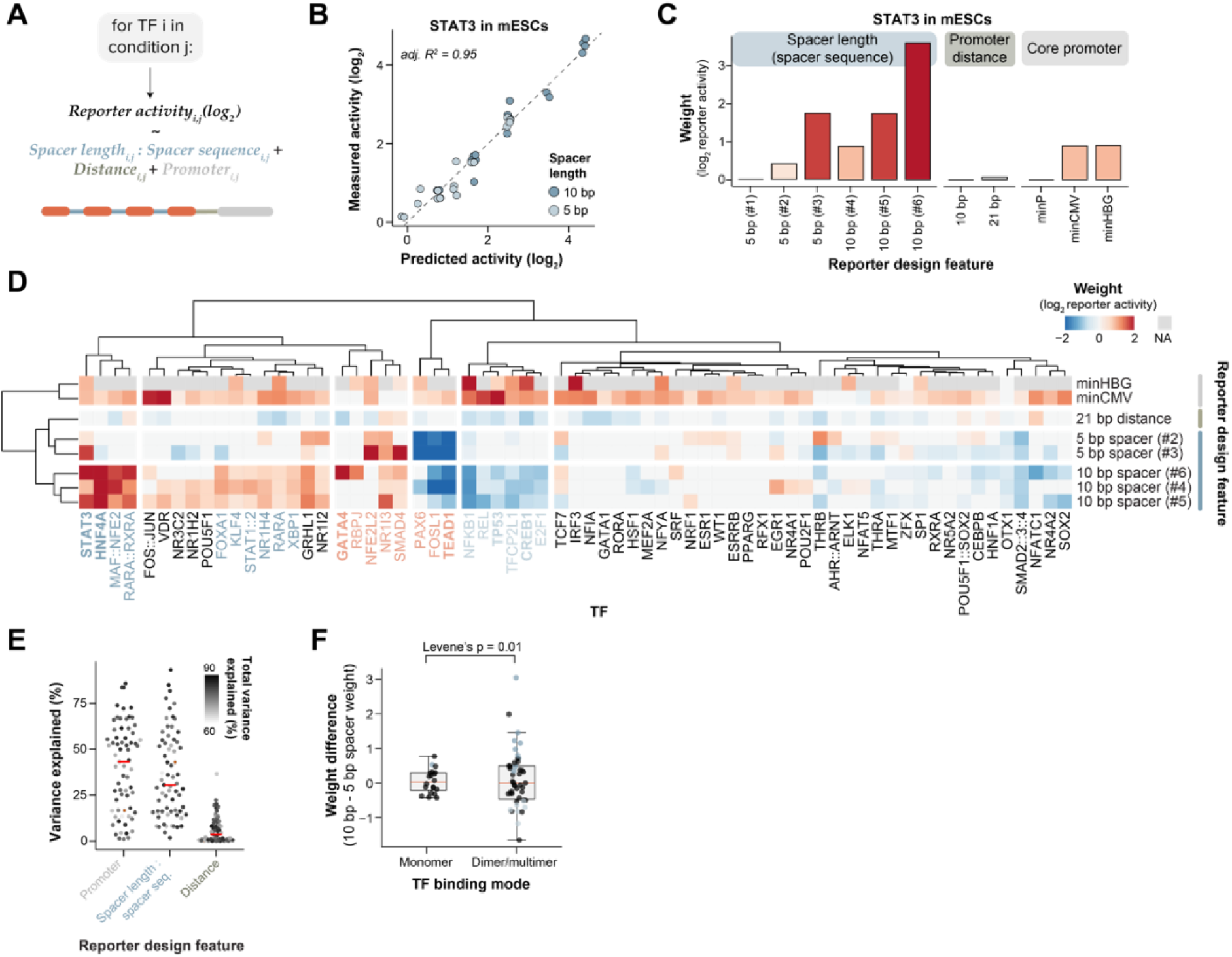
Identification of design features causative for TF reporter activity. (**A**) Equation of the log-linear model. Reporter design features are indicated by color and reflect the reporter design cartoon underneath the equation. (**B**) Correlation between measured STAT3 reporter activities and predicted reporter activities by the log-linear model in mESCs. Color-coded are the two spacer lengths. (**C**) Weights of the individual coefficients of the STAT3 log-linear model. The color indicates the strength of the weight. (**D**) Coefficient weight heatmap for all TFs with a significant log-linear model fit and a total explained variance of >50%. As in **C**, all weights are computed in contrast to the reference variables minP (core promoter), 10 bp (promoter distance), and #1 (spacer sequence). Weights of features that did not significantly contribute to the model (p ≥ 0.1) are set to 0 in this visualization. TFs highlighted in red display spacer sequence preferences, TFs in blue spacer length preferences. TFs indicated in bold are mentioned in the text. (**E**) Total variance explained by the individual design features for all TFs displayed in **D**. Red line indicates the median. The color of the dots indicates the total explained variance of the log-linear model. (**F**) Average difference between the weights of 10 bp spacer sequences in the log-linear model and the 5 bp spacer sequences, separately per monomeric and dimeric/multimeric TFBSs. Statistical significance of the difference in variance is estimated by Levene’s test.

#### Common rules to design active TF reporters

Next, we asked whether active TF reporters can be designed according to universal reporter design rules, or whether each TF requires its own specific rules. We applied the log-linear model analysis to each of the 86 probed TFs, focusing on the cell line and culture condition in which the TF is most active (see Methods). For 67 out of 86 TFs (78%) the models reached statistical significance (adjusted p-value < 0.05; **Figure S3A**) and explained >50% of the variance in reporter activity. For these models we then extracted the underlying weights of the individual reporter design features (**Figure 2D**). This analysis revealed several important insights. First, for almost all tested TFs, reporters were more active when having a minCMV or minHBG promoter compared to a minP promoter. Note that the fitted activities are normalized to the background activity of the core promoter (as described in the “*Data overview”* section), i.e., reporter activities are defined here as the TF-induced activity change compared to the promoter-only activity. Thus, minCMV and minHBG promoters allow for stronger induction, regardless of the TF. Second, although the promoter distance explained the least variance compared to all other investigated features (**Figure 2D, E**), the majority of TFs had a slightly decreased activity when the core promoter was placed 21 bp away from the first TFBS instead of 10 bp. This suggests that for many TFs placing the TFBS closer to the TSS can subtly increase transcription activity.

#### TFBS spacer length can affect activity

Besides the generic role of the core promoter and the core promoter distance, we found a striking TF-specific role for the spacer length between the TFBSs. For ten TFs, all three 10 bp spacer sequences consistently increased activity compared to the 5 bp spacer sequences (**Figure 2D**, TFs highlighted in dark blue). A readily interpretable example is HNF4A, for which > 90% of all variance in the reporter activity was caused by changing the spacer length from 5 to 10 bp (**Figure 2D, E**); this increased reporter activity by roughly 8-fold on average (**Figure S3B**). Conversely, six TFs had significant negative weights for all three 10 bp spacer sequences, and hence favored the shorter 5 bp spacer length (**Figure 2D**, highlighted in light blue). We then examined in greater detail which TFs exhibited these spacer length-preferences. Interestingly, we observed that TFs that bind DNA as monomers tended to be unaffected by changes in spacer length, while dimeric or multimeric TFs had significantly stronger spacer length-preferences (**Figure 2F**). In fact, 15 out of 16 TFs with consistent spacer length-preferences were TFs that bind to its TFBS as dimer or multimer. Possibly, dimeric or multimeric TF assemblies have more complex DNA interactions and might therefore need precise relative positioning to be able to activate efficiently from adjacent TFBSs. For some TFs (e.g., CREB1^29^, TP53^32^) it was previously described that optimal helical positioning of adjacent TFBSs (i.e., on the same face of the DNA helix) facilitates robust activation. We found similar TFBS spacer length preferences for these described TFs, and identified many more candidate TFs that might have similar helical positioning dependencies.

#### Several TFs benefit from specific spacer sequences

Besides TFs that clearly require certain spacer lengths to effectively activate, several TFs showed strong preferences for individual spacer sequences (**Figure 2D**, highlighted in red). For GATA4, for instance, only spacer sequence #6 (spacer length of 10 bp) significantly contributed to reporter activity, while for TEAD1 spacer sequence #1 (spacer length of 5 bp) was the only spacer sequence with strong activation. These specific preferences might be caused by an increased or decreased affinity for TF binding due to the sequences surrounding the TFBS, as has been reported before. ^40,41^ Although we ensured that all spacer sequences are devoid of any known TFBS, we cannot rule out that an unknown TF binds the spacer and synergizes with the TF for which the reporter was designed. Together, our log-linear model analysis revealed that TF reporter design can be optimized regardless of the TF through the choice and positioning of the core promoter. Nevertheless, many TFs require TF-specific spacer lengths or spacer sequences for efficient activation, underscoring the importance of systematic reporter design optimization.

### Cell type dependence of TF reporter activities

#### Correlating reporter activities with TF abundance across cell lines

After identifying the features facilitating high reporter activity, we aimed to characterize the TF specificity of each reporter. One line of evidence for such specificity would be if the activity of a reporter correlates positively with the abundance of the corresponding TF across the nine tested cell lines. Therefore, we collected publicly available mRNA-seq data for eight cell lines and generated data for mNPCs (see Methods). We then conducted a transcript abundance correlation analysis for 37 TFs that showed sufficient variation in expression level across the cell lines (**Figure 3A**). For some TFs (e.g., POU5F1::SOX2, HNF1A) the activities of all reporters significantly correlated with TF transcript abundance. For other TFs (e.g., IRX3), none of the reporters had a significant correlation. While this could indicate that the reporters for these TFs lack specificity, it is also possible that the activity of those TFs is controlled primarily by intracellular signaling or by certain co-factors; alternatively, their protein abundance is not reliably predicted by their mRNA level.

**Figure 3:**
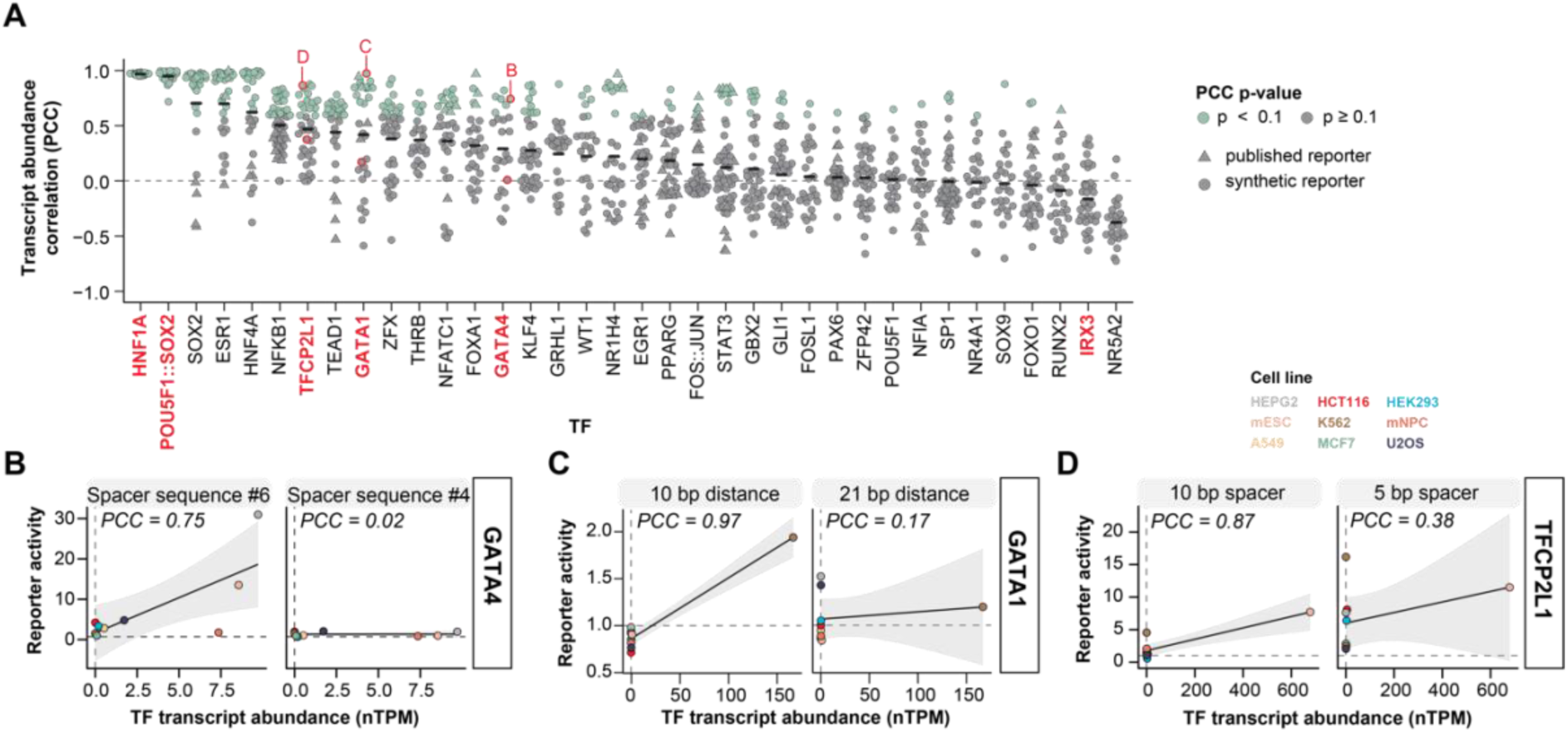
Investigating cell type specificity of reporters. (**A**) Correlations between TF reporter activities and TF transcript abundances across the nine probed cell lines per TF. Only TFs with variable expression across the nine cell lines are included (see Methods). The black solid line indicates the mean PCC per TF. TFs highlighted in red are mentioned in the text. Dots highlighted with a red stroke are depicted in **B-D**. (**B**) Correlation between GATA4 transcript abundance and reporter activity for a highly (spacer sequence #6) and a poorly correlating reporter (spacer sequence #4). The two displayed reporters are identical except for the spacer sequence mentioned above the panels. Cell lines are color-coded. Solid line indicates linear regression, grey shade indicates standard deviation. nTPM = normalized TPM (see Methods). (**C**) Same as **B**, but for GATA1 and two promoter distances. (**D**) Same as **B**, but for TFCP2L1 and two spacer lengths.

#### Using expression correlation to identify optimal reporters

For most TFs only a subset of reporters significantly correlated with TF transcript abundance (e.g., GATA4, TFCP2L1, GATA1; **Figure 3A**, highlighted in red). For example, one GATA4 reporter design with spacer sequence #4 was not active in any cell type, but the same design (i.e., the same core promoter, promoter distance and spacer length) with spacer sequence #6 was more active in cell types where GATA4 is expressed (HEPG2, mESC; **Figure 3B**). Indeed, GATA4 reporters with spacer sequence #6 almost exclusively displayed activities that significantly correlated with *GATA4* transcript abundance (**Figure S4A**), suggesting that this spacer sequence renders GATA4 reporters GATA4-specific. In line with these findings, spacer sequence #6 was also identified as the most important feature in the log-linear model for GATA4 (note that this model was fit in HEPG2, **Figure 2D**).

#### Additional examples of design-dependent TF specificity

GATA1 reporters were more GATA1-specific (i.e., activity only in K562) with a 10 bp rather than a 21 bp promoter distance (**Figure 3C**, **S4B**, **2D**). The latter displayed activity in GATA1-lacking cell types, possibly because these reporters respond to other GATAs (e.g., GATA3 in MCF7 or GATA4 in HEPG2). TFCP2L1 reporters give another example of design-dependent TF specificity. We found that a TFCP2L1 reporter with a 10 bp spacer length (spacer sequence #4) was predominantly active in the cell line where TFCP2L1 is highly expressed (mESC), while the same reporter with a 5 bp spacer length (spacer sequence #1) was also highly active in other cell types (**Figure 3D**). Indeed, all TFCP2L1 reporters with spacer sequence #4 and #5 (both 10 bp) displayed activities that significantly correlated with TFCP2L1 transcript abundance (**Figure S4C**). Activities of TFCP2L1 reporters in TFCP2L1-lacking cell types might be explained by response to GRHL1, which is a TF with a highly similar binding motif (**Figure S1A**), but a distinct expression pattern (GRHL1 is lowly expressed in all nine cell lines). Together, these findings highlight that fine-tuning the reporter design can substantially improve the specificity, even for TFs with highly similar TFBSs.

### Response of TF reporters to pathway stimulation and inactivation

#### Experimental design of pathway perturbations

Many TFs are known to depend on specific stimuli or upstream signaling events for their activity. To further test the responsiveness of the reporters, we therefore applied a total of 23 different pathway inhibitors, ligands, drugs and culture conditions that are known to influence the activity of at least one of the TFs (**Figure 4A, Table S3**). For each perturbation we chose one cell type that was most likely responsive to this stimulus. Altogether, we expected these perturbations to activate 27 TFs and suppress 9 TFs within our set of 86 TFs.

**Figure 4:**
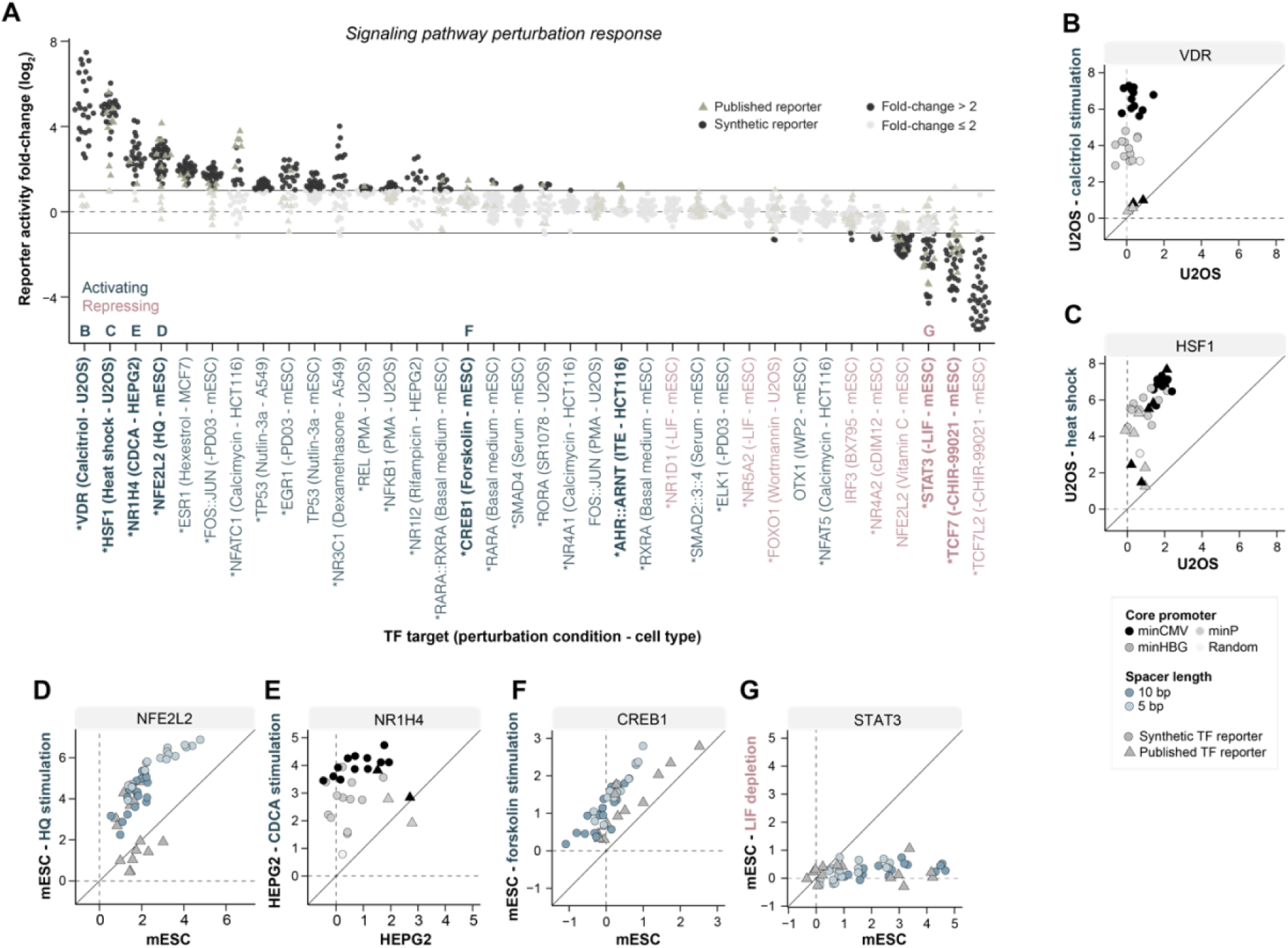
Response to signaling pathway perturbations. (**A**) Change in TF reporter activities upon signaling pathway perturbations. Shown are only the responses of the direct targets of the perturbations. Activating perturbations are shown in blue, repressing conditions in purple. Conditions that were selected as best perturbation condition for the TF (i.e., strongest average response of the tested perturbations for that TF, see Methods) are denoted by an asterisk. TFs highlighted in the text are indicated in bold. TFs depicted in **B-G** are indicated by letter. LIF = leukemia inhibitory factor. PMA = phorbol 12-myristate 13-acetate, HQ = hydroquinone, CDCA = chenodeoxycholic acid. (**B-G**) Response of reporters of six different TFs to TF-targeted pathway perturbation conditions. TFs for which reporters are displayed is denoted on top of each figure. Reporter activities (log_2_) in the basal condition are displayed on the x-axis and in the perturbation condition on the y-axis. Reporter design features are indicated by color, published reporters by shape.

#### Examples of strong responses

For some of those TFs (e.g., HSF1 upon heat shock, TCF7 upon removal of WNT activator CHIR-99021), we saw robust responses across almost all reporter designs (**Figure 4A**). The most potent TF-stimulating condition was activation of vitamin D receptor (VDR) reporters by its ligand calcitriol. In U2OS cells this yielded activation levels up to 180-fold (**Figure 4A, B**). Other strong reporter responses were also achieved by stimulating the heat shock-responsive HSF1 at 43°C (**Figure 4C**); the oxidative stress response factor NFE2L2 by treatment with hydroquinone (**Figure 4D**); the bile acid receptor NR1H4 by the bile acid CDCA (**Figure 4E**); the c-AMP responsive TF CREB1 by c-AMP activator forskolin (**Figure 4F**); and STAT3 by removal of JAK-STAT activator LIF (**Figure 4G**).

#### Variation in responses between reporter designs

Overall, there was a marked variation in the strength of the response between reporters of the same TF. The strength of the responses in the examples above strongly depended on the core promoter (VDR, HSF1, NR1H4), or the spacer sequences (NFE2L2, STAT3), which is in line with the findings of the log-linear model (**Figure 2D**). For other TFs (e.g., AHR::ARNT, NR4A2), only a few the reporters showed a clear response (fold-change > 2). The published reporters for VDR and NR1H4 showed a relatively poor response, as did a subset of the published NFE2L2, CREB1, HSF1 and STAT3 reporters. In total, of the 36 TFs targeted by the 23 perturbations, for 25 TFs we identified at least one reporter that responded in the expected direction by at least 2-fold (**Figure 4A**).

### Testing reporters by TF depletion or overexpression

#### Altered TF expression: experimental design and interpretation

Finally, as a more direct method of perturbing TF activity, we tested the response of all reporters to transient knockdown (KD), protein degradation, or overexpression of individual TFs. Among our set of 86 TFs, we knocked down 16 TFs in mESCs and 28 TFs in HEPG2 cells by RNA interference. For SOX2 and POU5F1 we additionally used degron-mediated depletion in mESCs.^42^ Moreover, to evaluate specificity and off-target responses of the TF reporters, we also included nine KDs in mESCs and 13 KDs in HEPG2 cells of related TFs that have similar TFBSs as our candidate TFs. Finally, we overexpressed four TFs that are not naturally expressed in mESCs. The scale of these experiments prohibited the verification of the KD or overexpression efficiency for each individual TF by Western blotting or mass-spectrometry. For this reason, a lack of a response of reporters to the perturbation of their cognate TF does not necessarily imply that the reporters lack specificity; it is possible that we simply failed to alter the level of the TF sufficiently. Conversely, however, a strong response of reporters to the perturbation of the cognate TF can be regarded as evidence of specificity.

#### Overall response of reporters

The results of these experiments are summarized in **Figure 5A**. Approximately one-third of all KD-targeted TFs showed a strong decrease in reporter activity (fold-change > 2) across the majority of reporters, although for most of these TFs the strength of the response varied substantially between reporters. Protein degradation of SOX2 strongly reduced activities of all POU5F1::SOX2 reporters, and a subset of SOX2 reporters. Similarly, POU5F1 degradation decreased activity of a subset of POU5F1 reporters and all POU5F1::SOX2 reporters. Overexpression of FOXA1 significantly increased the majority of the FOXA1 reporters, while FOSL1 overexpression only led to an increase in FOS::JUN, but not FOSL1 reporter activities. GATA1 and NR4A2 overexpression did not increase activities of their target reporters.

**Figure 5:**
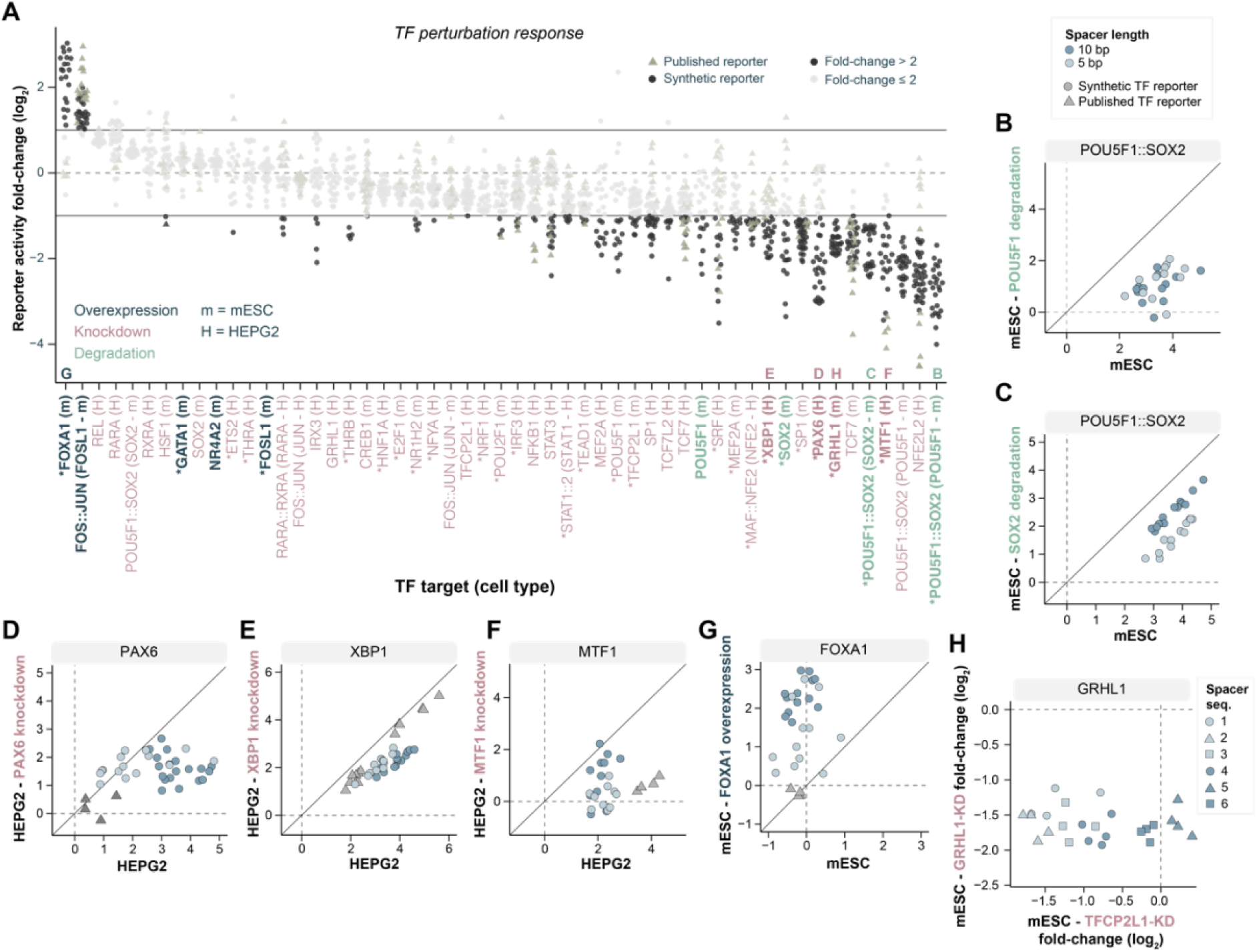
Response of reporters to direct TF perturbation. (**A**) Change in TF reporter activities upon direct TF perturbation. In some cases the target TF consists of two TFs (e.g., POU5F1::SOX2); the perturbed TF is then indicated in the x-axis labels. TF overexpression is shown in black, TF knockdown in purple, and TF degradation in green. Conditions that were selected as best perturbation condition for the TF are denoted by asterisk. TFs highlighted in the text are indicated in bold. TFs depicted in **B-H** are indicated by letter. (**B-G**) Response of TF reporters to six different direct TF perturbation conditions. TFs for which reporters are displayed is denoted on top of each figure. Reporter activities (log_2_) in the basal condition are displayed on the x-axis and in the TF perturbation condition on the y-axis. (**H**) Response of GRHL1 reporters to GRHL1 (y-axis) and TFCP2L1 knockdown (x-axis).

#### Perturbation response depends on reporter design

Again, we found that the reporter responses were often dependent on the precise design. While all POU5F1::SOX2 reporters strongly reduced their activity upon POU5F1 degradation (**Figure 5B**), there was a marked difference in response to SOX2 degradation, with POU5F1::SOX2 reporters with a 10 bp spacer length showing stronger responses (**Figure 5C**). Similarly, we found that PAX6 reporters with reduced activity upon PAX6 KD mostly had 10 bp spacers, while the published PAX6 reporters did not show any response (**Figure 5D**). Other examples of design-dependent responses are highlighted in **Figures 5E-G**. Overall, of the 44 TFs that were targeted by KD, 34 had at least one reporter with a more than two-fold reduction in activity (**Figure 5A**).

#### Probing reporter cross-reactivity

Many TFs belong to families that share highly similar binding motifs. Therefore, to test for off-target responses we also evaluated responses upon perturbations of other members within the same TF family. In total, we investigated 50 pathway perturbations and 87 TF perturbations that could potentially result in cross-reactivity due to TFBS similarity of the target TF and another TF. Of these, reporters for around 20 TFs showed substantial off-target responses (**Figure S5A, B**). A striking example of high selectivity is NR1H4 reporters, which have a TFBS that is highly similar to other nuclear receptor TFBSs (**Figure S1A**); nevertheless, they strongly responded only to bile acid stimulation (CDCA) and not to any other nuclear receptor stimulation (**Figure S5C**). Off-target responses often varied in magnitude depending on the reporter design. For example, all GRHL1 reporters had a reduced activity upon KD of GRHL1, while only GRHL1 reporters with spacer sequences #1-4 additionally responded to TFCP2L1 KD (**Figure 5H**). We found that CLOCK reporters, for which we only probed published reporter designs, reduced their activity by approximately twofold upon removal of LIF (**Figure S5D**); these reporters carry a repeat sequence that significantly matches the STAT3 motif (**Figure S5E**), possibly explaining the erroneous response to LIF.

### A collection of “prime” TF reporters

#### Assigning confidence levels to TF reporters

Using the abundance of the cell type-specific activities and the perturbation data described above, we aimed to integrate all data to identify the most optimal reporters for each TF. To do so, we assigned confidence levels to each individual reporter, ranging from 0 (low confidence) to 4 (very high confidence), based on the criteria summarized in **Figure 6A**. For level 4, we required reporters to be responsive to a relevant stimulus, display activities that correlate with the abundance of the TF across the tested cell lines, and show a substantial response to depletion or overexpression of the TF, without responding to off-target perturbations. **Figure 6B** illustrates how each of the confidence level criteria contributes to the confidence scores of all STAT3 reporters. Out of 51 reporters, 21 had a confidence level of 0 because they did not display any significant activity, and also did not respond to LIF removal. Only six reporters were assigned level 4 because they displayed high activity in basal conditions, correlated with STAT3 abundance, strongly responded to LIF removal, and did not show an off-target response to STAT1 KD (**Figure S5A**). As established previously (**Figure 2B-D, 4G**), these high-confidence reporters are characterized by a 10 bp spacer sequence #6, but also include published reporters. We generated similar reporter confidence heatmaps for all 86 TFs (**Figure S6**).

**Figure 6:**
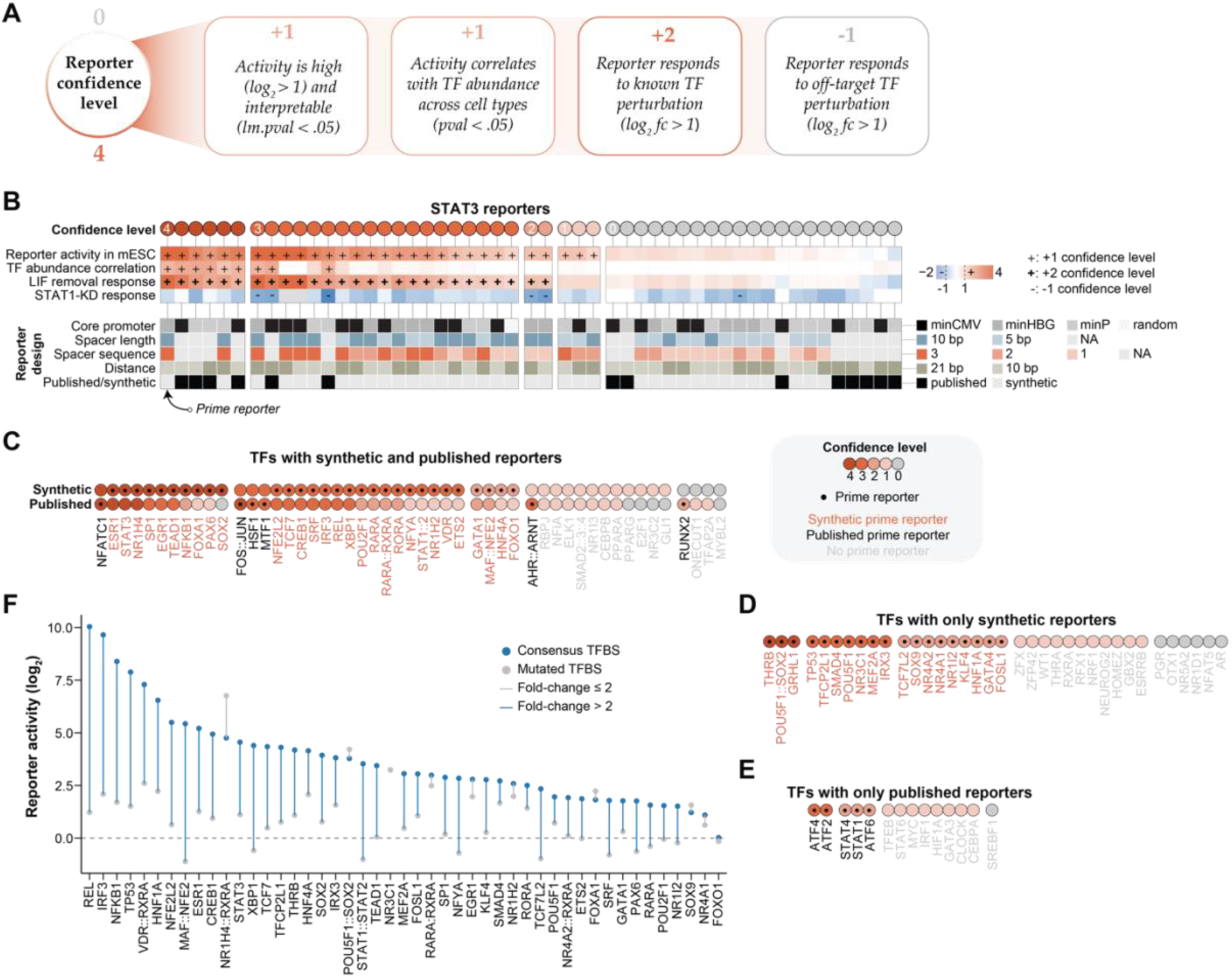
Identification of TF-specific and sensitive reporters. (**A**) Reporter confidence levels are defined based on the four threshold criteria mentioned in the boxes. Response to known TF perturbation is given a higher weight due to its importance. (**B**) Reporter confidence scores of STAT3 reporters. Reporter activity, TF abundance correlation, or TF perturbation response meeting the threshold criteria outlined in **A** contribute to the reporter confidence level and are denoted by a plus or minus sign. (**C**) Overview of the confidence level of the best reporter per TF for TFs with both synthetic and published reporters probed. (**D**) Same as **C** but for TFs with only synthetic reporters probed. TP53 and NR3C1 are included in this list because their published reporters were not probed in TP53/NR3C1 perturbation conditions, prohibiting comparisons between synthetic and published reporters. (**E**) Same as **C** and **D** but for TFs for which only published reporters were included in the reporter library design. (**F**) Reporter activity of the 60 prime reporters with consensus TFBS (blue dot) and mutated TFBS (grey dot). Activities displayed are from the same conditions as used for the log-linear models.

#### Selecting the set of prime reporters

Finally, for TFs with reporters with a confidence level of 2 or higher, we selected a single “prime” reporter, based on the confidence scores and – in case of ties – additional performance criteria (**Table S4**; see Methods). For a total of 60 TFs, this yielded a prime reporter with confidence level 4 (15 TFs), 3 (28 TFs) or 2 (17 TFs). We emphasize that level 2 means that the reporter is significantly active and that there is evidence for TF specificity, and thus such a reporter is likely to provide meaningful information. While most prime reporters feature a minCMV or minHBG core promoter (46/60), the spacer sequences are distributed relatively evenly across prime reporters (#1 (5 bp): 13, #2 (5 bp): 4, #3 (5 bp): 7, #4 (10 bp): 10, #5 (10 bp): 6, #6 (10 bp): 9), highlighting their TF-specific nature. This underscores the necessity for TF-specific spacer sequence optimization. Furthermore, the set of 60 prime reporters consists of 49 synthetic reporters and 11 published reporters. Notably, of the 36 TFs in the prime reporter set for which we probed both synthetic and published reporters, synthetic reporters outperformed the published reporters for 30 TFs (83%), while published reporters outperformed the synthetic reporters for only 6 TFs (**Figure 6C**). For 18 TFs, the synthetic prime reporters even scored at least one confidence level higher than the published reporters. This demonstrates the value of systematic optimization. Additionally, the prime set includes 19 TFs for which we did not test published reporters, primarily because they were not available, (**Figure 6D**), and five published reporters for which we did not test synthetic designs (**Figure 6E, Table S4**).

#### Prime reporters typically require high-affinity BSs

As a final characterization of the synthetic prime reporters, we checked whether their activities are dependent on full integrity of the respective TFBSs (**Figure 1A, Table S1**). Indeed, of the 47 synthetic prime reporters for which we had matched mutated TFBS controls, 37 decreased their activity upon mutation of a two to three nucleotides in the TFBS (see **Table S1**) by at least 2-fold, and up to 500-fold (**Figure 6F**). Prime reporters also had a significantly increased sensitivity to these mutations compared to reporters of the same TF with a lower confidence level (**Figure S7**). These strong responses to minimal alterations in the TFBS reaffirm the TF specificity of the identified prime reporters. We note that the remaining 10 reporters (of which four are confidence level 4, and three are confidence level 3) should not be rejected based on this result, because some TFs might be able to activate a promoter stronger through low- or medium-affinity TFBSs than through high-affinity TFBSs.^32^

### Utilizing prime reporters for accurate multiplexed TF activity detection

#### Specific TF activity detection across nine cell lines

Having identified the prime reporters for 60 TFs, we reassessed the activities of those TFs across all tested conditions. We first focused on the steady-state activities across the nine probed cell lines (**Figure S8A**). To be able to compare reporters of different strengths with each other, we rescaled the reporter activities separately per TF. This allowed us to identify cell type-specificities of TFs and to identify clusters of TFs with similar activity patterns (**Figure 7A, S8B**). We found a large number of TFs displaying distinct cell type-specific activities, which match their known biological functions (e.g., HNF4A in HEPG2, ESR1 in MCF7, or SOX2 in mESC; **Figure 7A, S8C**). The prime reporters also discriminate TFs with highly similar TFBSs, like GATA1/GATA4, TFCP2L1/GRHL1, EGR1/KLF4, or a variety of nuclear receptor TFs. Thus, our set of 60 prime reporters can identify TF activity differences between cell types, and highlight functional similarities between TFs.

**Figure 7:**
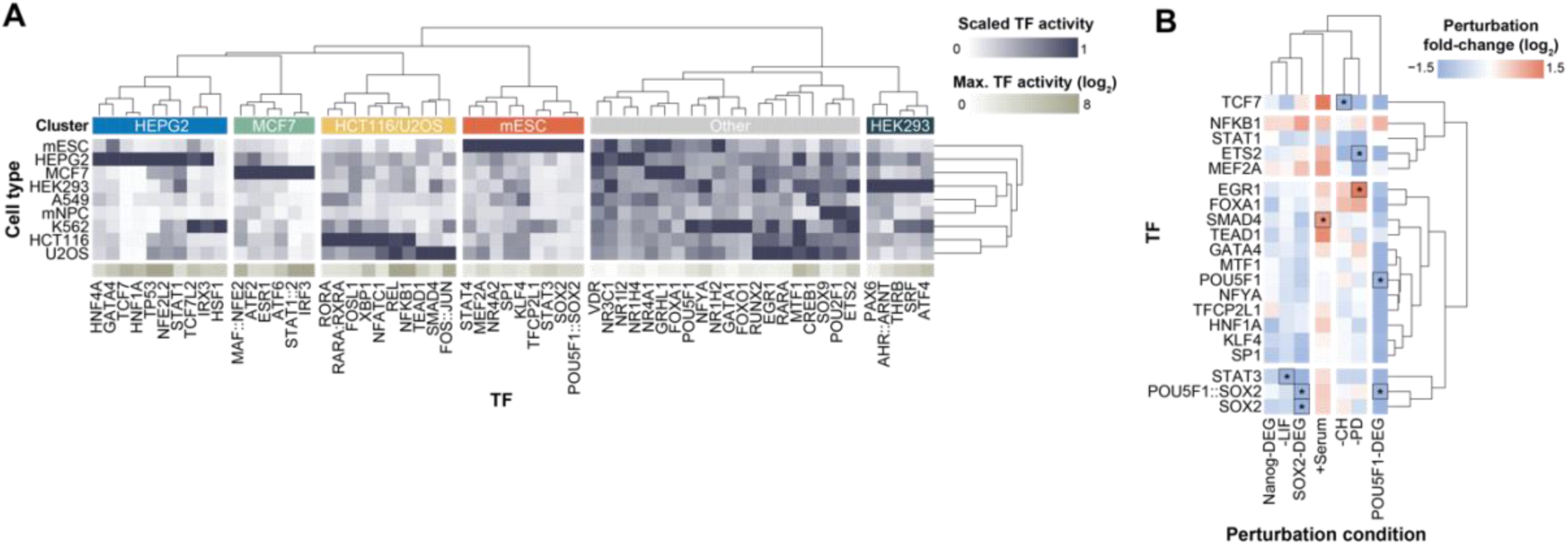
Multiplexed detection of TF activities with prime reporters. (**A**) TF activities as measured by the 60 prime reporters across all nine probed cell lines. Activities were scaled by dividing the reporter activities by the maximum activity per TF. (**B**) Changes in prime reporter TF activities upon various TF perturbations in mESCs. TF targets of perturbations are indicated by black rectangles and asterisks. Only TFs expressed in mESCs (nTPM > 4) and with a substantial perturbation response (fold-change > 2) in at least one condition are displayed. DEG = degradation.

#### Exploring TF-TF communications

Besides steady-state activities, the prime reporters reveal the dynamics of 60 TF activities across all tested 98 TF perturbation conditions. As an example, we quantified prime reporter responses upon all KDs in HEPG2 cells with a strong effect on their direct target (n = 21). We found a large number of TFs that change their activity upon downregulation of another TF (e.g., PAX6 activation upon HNF1A KD, **Figure S8D**). These data offer a large resource to explore cascades of TF activities.

#### Signaling interdependencies in the pluripotency network

We then focused our analysis on perturbations in mESCs that affect the pluripotency network (**Figure 7B**). Interestingly, besides altering the activity of its cognate TF, most perturbations led to strong secondary TF activity changes. For instance, we found that degradation of key pluripotency factors POU5F1 and SOX2 substantially reduced the activity of other pluripotency TFs like STAT3, TFCP2L1 or KLF4, highlighting their core function in the pluripotency network.^38^ Furthermore, removal of JAK-STAT activator LIF not only led to strong inactivation of its target STAT3, but also decreased the activity of WNT target TCF7 as well as many other pluripotency TFs like SOX2 or KLF4 (**Figure 7B**). This suggests that LIF is needed to maintain pluripotency, potentially through crosstalk with the WNT signaling pathway. Similarly, we found that MEK-ERK inhibitor PD (PD0325901) crosstalks with WNT signaling, and WNT activator CH (CHIR-99021) with MEK-ERK signaling, suggesting that these signaling pathways reinforce each other and have redundant targets, as has been discussed before.^38,39^ Besides this, we found that addition of serum increased the activity of pluripotency TFs such as POU5F1::SOX2, reinforcing the pluripotency network. Together, this analysis shows that multiplexed TF activity detection using prime reporters has the potential to link targeted signaling pathway perturbations to functional changes in TF activity to discover signaling pathway interdependencies.

## DISCUSSION

### Applicability of the identified prime reporters

We here present the systematic design and identification of a large collection of optimized “prime” TF reporters. This collection encompasses reporters that significantly outperform currently available reporters (e.g., VDR, SOX2, PAX6), and reporters for TFs for which no reliable reporters were available yet (e.g., GATA4, TFCP2L1, KLF4). The sequences of the prime reporter for each TF are documented in **Table S4**, which can be used for various purposes. For instance, the prime reporters can be used individually in a conventional fluorescence/luminescence reporter assay to better characterize the role of single TFs in certain biological processes. Alternatively, the identified 60 prime reporters can be employed in a multiplexed fashion, where each TF drives a unique barcode. Signaling pathways could be challenged by an array of inhibitors or activators, similar to what has been done in this study, to unveil novel roles of TFs in signaling pathways. Likewise, TF responses can be tracked upon TF depletion to dissect TF-TF communications. Potentially, this can also be done in single cells and in time-course experiments to detect cascades of TF activities. Although the prime reporters are top-rated based on our performance criteria, there may be instances where other reporters with specific attributes are preferred for certain TFs (e.g., high cell type-specificity or responsiveness to perturbation of related TFs). **Figure S6** can aid in identifying such cases (e.g., for identifying generic STAT reporters instead of STAT3-specific reporters).

### Increased TF specificity of prime reporters

We have shown that our synthetic reporters outperform published reporters for >80% of all comparisons. This underscores that a subset of currently available reporters is suboptimal in terms of sensitivity (e.g., VDR, PAX6, NR1H4) or specificity (e.g., CLOCK, TP53, POU2F1). In comparison to published TF reporters, which rely on genomic response elements or unoptimized synthetic designs, the designed prime reporters exclusively contain TFBSs for the candidate TF, and are highly optimized to enable effective transcription. Through careful optimization of the spacer sequences between the TFBSs and the choice of the core promoter, we were able to achieve reporters with increased TF sensitivity and specificity. In some cases, this even enabled us to identify specific reporters for TFs with highly similar TFBSs (e.g., GATA1/GATA4, TFCP2L1/GRHL1). While we established prime reporters for 60 TFs, good reporters for many other TFs are still lacking. For instance, our set of TFs did not include a large number important activator TFs that belong the basic domain or homeodomain TF superclass, many of which have non-unique binding motifs. These TFs can have crucial roles in development (e.g., HOX TFs), hence, generating reporters for these TFs would be important to dissect the roles of TFs during differentiation. Although it remains challenging to generate specific reporters for TFs with non-unique TFBSs, careful optimization of TFBS spacer sequences and thorough evaluation of reporter responses to a variety of target TF and off-target TF perturbations could offer solutions.

### An alternative to TF activity inference

We envision that multiplexed TF reporter measurements could complement indirect TF activity inference methods that rely on ATAC-seq, ChIP-seq, or RNA-seq data. While these methods are able to impute activities for any TF with a reliable motif from commonly available datasets, they are not necessarily predictive of transcriptional activity and remain inferential.^14^ Furthermore, TF inference methods often struggle to discern the activity of individual TFs, instead reporting on the activity of TF clusters sharing similar TFBSs.^9,43^ Multiplexed (prime) TF reporter assays offer an orthogonal approach that provides functional evidence of TF activity with high specificity for the candidate TF.

## MATERIALS & METHODS

### TF reporter library design

The 86 TFs were manually chosen by reviewing all human TFs. Selection criteria included motif quality, motif uniqueness, expression patterns, and perturbation opportunities. Motif quality and uniqueness was assessed using a previous review and curation of available motifs for all human TFs.^28^ Mainly TFs with a unique motif were selected, which ensured to capture a wide diversity of motifs within the human TF motif landscape. TFs with unique motifs, but no known activator function were not included. Some TFs with non-unique motifs but distinct expression pattern or ligands were also selected; we reasoned that reviewing specificity for these TFs would be feasible by testing the reporter in different cell types or upon perturbation. For each TF, consensus TFBSs were generated by taking the most conserved base at each position, and mutated TFBSs were created by mutating at least two and up to four conserved bases (**Table S1**). In addition to the mutated TFBSs, three random TFBS-devoid (TF-neg) 11 bp sequences were included as negative controls. The absence of TFBSs of known activator TFs was confirmed in the mutated and random sequences using FIMO (p-value threshold 1e-4).^44^ Synthetic TF reporters were then created by placing four adjacent copies of the consensus, mutated, or negative TFBS. The four TFBSs were separated by *in silico*-designed TFBS-devoid spacer sequences with lengths of 5 or 10 bp. In total, three different spacer sequences were generated per spacer length. To do so, random sequences with a GC content of 40-60% were generated (sim.DNAseq function in R from package SimRAD (version 0.96)). These sequences were combined with 3 bp of the left and right side of all TFBSs and then scanned using FIMO (**Figure S1C**). For the two spacer lengths (5 and 10 bp), nine sequences with the fewest predicted significant TFBSs were selected and placed in between the TFBSs (three different spacer sequences per reporter, times the three spacer sequences). A similar approach was taken to generate three 10 or 21 bp spacer sequences in front of the core promoter. One of three core promoter sequences, minCMV,^33^ minHBG,^34^ or minP (derived from pGL4 (Promega, Madison, WI, USA)), was placed downstream of the TFBSs and spacer sequences, followed by a S1 Illumina adapter sequence and a unique 12-13 bp random barcode sequence (each unique construct was linked to five to eight different barcodes). All generated random barcodes had a Levenshtein distance of at least three with respect to one another and barcodes with an unbalanced GC ratio were removed (create.dnabarcodes function from the R package DNABarcodes (version 1.2.2) ^45^). For 64 TFs we also included published reporter sequences. The response element sequences were retrieved from three different sources (**Table S1**). ^26,27^ Promega pGL4.XX sequences were retrieved from https://www.snapgene.com/plasmids/luciferase_vectors. For some TFs, multiple TF response elements were included (see **Table S1** for all included published TF response elements). Again, each published response element was placed 10 or 21 bp upstream of a minP or minCMV core promoter. The same spacer sequence as for the synthetic TF reporters was used upstream of the core promoter. Several other controls were included in the design. First, to estimate the effect of the TFBSs alone, TF reporters with a TFBS-devoid core promoter were designed. This promoter was previously shown to be inactive.^32^ For each TF, this TFBS-devoid core promoter was attached to one reporter design only (background #4, promoter distance 21 bp). Second, two different positive controls were included to benchmark the expression levels of the synthetic TF reporters: 1) a 183-bp region of the hPGK promoter, and 2) 120 (40 for each of the three core promoters minP, minCMV, and minHBG) 100-bp regions of *Klf2* gene enhancers with known activity in reporter assays.^35^ Each of these control reporters were also linked to five to eight different barcodes. All reporter sequences were completed with 18 bp primer adapter sequences (that were also scanned using FIMO) in both flanks for cloning purposes. The resulting sequence pool had a total length of on average 202 bp (at least 148 bp up to 297 bp) and was ordered as oligonucleotide library from Twist Biosciences.

### Cloning of the TF reporter library

The vector backbone was constructed as mentioned previously.^32^ The oligonucleotide library was resuspended in TE buffer (Invitrogen) to a final concentration of 20 ng/µl. 10 ng of the oligonucleotide library was then PCR amplified (1’ 95°C, 6x(15’’ 95°C, 15’’ 57°C, 15’’ 72°C), 1’ 72°C) by MyTaq Red mix (Bioline) using primers that add overhangs with EcoRI (MT024, **Table S5**) or NheI (MT025) restriction enzyme sites. The PCR product was then purified using CleanPCR beads (#CPCR, CleanNA) at 1.8:1 beads:sample ratio, digested with EcoRI-HF (#R3101, NEB) and NheI-HF (#3131, NEB) by incubating the PCR product at 37°C for 1 h, and then again bead purified as before. 1 µg of the entry vector was also digested with EcoRI-HF and NheI-HF and the linearized product was purified from a 2% agarose gel using PCR Isolate II PCR and Gel Kit (Bioline). The digested and purified reporter pool was then ligated into 80 ng of the linearized entry vector using Takara ligation kit v1.0 (#6021; Takara) at a 1:3 (vector:insert) ratio. The ligation mix was then bead purified as before and transformed into MegaX DH10B T1R Electrocomp™ Cells (Invitrogen) using 1 µl of the ligation mix. The library complexity was estimated from plated serial dilutions of the transformed cells to be ∼300,000 colony forming units. Transformed cells were transferred to 200 ml standard Luria Broth (LB) plus kanamycin (50μg/ml), grown overnight and purified using a Maxi plasmid purification kit (#12162; Qiagen).

### Cell culture

MCF7 (#HTB-22, ATCC), HEK293 (#CRL-1573, ATCC), and A549 (#CCL-185, ATCC) cells were cultured in DMEM medium (#41966029, Gibco), K562 (#CCL-243, ATCC) in RPMI 1640 medium (#11875093, Gibco), U2OS (#HTB-96, ATCC) and HCT116 (#CCL-247, ATCC) in McCoy’s 5a medium (#26600023, Gibco) and HEPG2 (#HB-8065, ATCC) in MEM (#11095080, Gibco). All media were supplemented with 10% fetal bovine serum (FBS, Sigma). mESC (E14TG2a, #CRL-1821, ATCC) were cultured in 2i+LIF culturing media according to the 4DN protocol (https://data.4dnucleome.org/protocols/cb03c0c6-4ba6-4bbe-9210-c430ee4fdb2c/). The reagents used were neurobasal medium (#21103-049, Gibco), DMEM-F12 medium (#11320-033, Gibco), BSA (#15260-037, Gibco), N27 (#17504-044, Gibco), B2 (#17502-048, Gibco), LIF (#ESG1107, Sigma-Aldrich), CHIR-99021 (#HY-10182; MedChemExpress) and PD0325901 (#HY-10254, MedChemExpress), monothioglycerol (#M6145-25ML, Sigma) and L-Glutamine (#25030-081, Gibco). The mNPCs used in this study were differentiated from E14TG2a mESCs and cultured in mNPC medium as mentioned previously ^46^. HEK293T (#CRL-3216, ATCC) cells used for lentivirus production were cultured in DMEM-F12 (#11320-033, Gibco) supplemented with FBS (Sigma) and L-glutamine (#25030-081, Gibco). All cells used in this study were routinely tested for mycoplasm.

### Reporter library transfection and pathway perturbations

All cell lines except for K562 were transfected using lipofection. Per lipofection condition, 1.5×10^5^ cells were seeded in a 12-well and transfected 8 hours later by adding 1 µg of TF reporter plasmid library with 3 µl of Lipofectamine 3000 (#L3000150, ThermoFisher) in 100 µl Opti-MEM (#31985070, Gibco). mESCs were plated directly before lipofection instead of 8 hours prior and transfected using Lipofectamine 2000 (#11668027, ThermoFisher). K562 cells were electroporated using an Amaxa 2D Nucleofector. Per transfection, 1×10^6^ K562 cells were resuspended in transfection buffer (100 mM KH2PO4, 15 mM NaHCO3, 12 mM MgCl2, 8 mM ATP, 2 mM glucose (pH 7.4)) supplied with 1 µg of plasmid library and electroporated using program T-003. After nucleofection, cells were resuspended in 2 mL complete medium and plated in 6-well plates. For the signaling pathway perturbation conditions, inhibitors or activators were added to the cells directly after transfections. All inhibitors and activators used in this study are mentioned in **Table S3**. 24 hours after transfection, cells were harvested and resuspended in 800 µl TRIsure (#BIO-38032; Bioline) and stored at -80 °C until further use. Transfections were done at least in biological duplicates on separate days.

### siRNA TF knockdown experiments

The TF knockdown experiments were performed in HEPG2 and mESCs. For HEPG2 cells, reverse siRNA transfections were done by mixing 20 nM siRNA with 1.5 µl Lipofectamine RNAiMAX transfection reagent (#13778075, ThermoFisher) in 100 µl Opti-MEM in 24-wells. Then, 7.5×10^4^ HEPG2 cells were added to the wells. The list of siGENOME SMARTpool siRNAs (Dharmacon) used in the screen can be found in **Table S3**. 24h after siRNA transfection, 0.5 µg of the TF reporter plasmid library was transfected by mixing the library with 1.5 µl Lipofectamine 3000 in 50 µl Opti-MEM and adding the mix directly to the cells. For mESCs, 1.5×10^5^ cells were reverse lipofected in 12-wells using 40 nM siRNA and 3 µl Lipofectamine RNAiMAX transfection reagent (#13778075, ThermoFisher) in 200 µl Opti-MEM. All used ON-TARGETplus siRNAs (Dharmacon) are listed **Table S3**. 24h after siRNA transfection, 1 µg of the TF reporter plasmid library was mixed with 3 µl Lipofectamine 2000 in 100 µl Opti-MEM and plated in new 12-wells. The siRNA-transfected mESCs were then collected and added to new 12-wells with the TF reporter plasmid library lipofection mix. Knockdown efficiency was evaluated by killing controls using siRNAs targeting PLK1 (# L-003290 (human), #L-040566 (mouse), Dharmacon). Non-targeting siRNAs were used as negative controls (#D-001210-01, Dharmacon). 24 hours after TF reporter library plasmid transfection and 48 hours after siRNA transfection the cells were harvested as mentioned in the “*Reporter library transfection and pathway perturbations*” section.

### TF overexpression experiments

Lentiviral plasmids carrying doxycycline-inducible open reading frames for GATA1, FOSL1, FOXA1, NR4A2 or RFX1 and a puromycin selection cassette were a kind gift from Bart Deplancke (EPFL, Lausanne, Switzerland).^47^ To generate lentivirus, 5×10^5^ HEK293T cells were plated in 6-well plates per condition. At ∼75% confluency, 1.5 µg TF ORF lentiviral plasmid was mixed with 1.125 µg psPAX2 (#12260, Addgene), 0.375 µg pMD2.G (#12259, Addgene) and 5 µl Lipofectamine 2000 in 250 µl Opti-MEM and added to the 6-wells. The medium was refreshed after 12 hours and lentivirus was collected after 48 hours from the supernatant. To transduce cells with the lentivirus, 1×10^5^ mESCs were plated in 12-wells in 500 µl 2i/LIF medium supplemented with 8.5 µg polybrene (#TR-1003, Sigma). Then, 500 µl of lentiviral supernatant was added to the cells. Medium was changed to fresh 2i/LIF medium 24 hours later and to puromycin-containing (2 µg/ml) 2i/LIF medium after 48 hours. Puromycin-resistant cells were grown and used for the subsequent TF reporter plasmid library transfection experiments. To transfect the TF reporter plasmid library, the TF ORF-carrying mESCs were pretreated for 24 hours with 2 µg/ml doxycycline (#D9891, Sigma) and then lipofected as mentioned in the “*Reporter library transfection and pathway perturbations*” section.

### TF degradation experiments

mESCs with FKBP-tagged POU5F1 (genetic background: V6.5)^48^, SOX2 (IB10), or NANOG (E14tg2a) were generated as described previously^42^ and were a kindly provided by Elzo de Wit (Netherlands Cancer Institute). TF degradation was induced directly after TF reporter library transfections using 500 nM dTAG-13 (#SML2601, Sigma). Cells were harvested for RNA extraction 24h after library transfection and degradation induction.

### RNA extraction, reverse transcription and barcode amplification

RNA extraction was done using the standard procedure according to the TRIsure protocol. After RNA extraction, 1 µg of RNA was treated with DNase I for 30 minutes (#04716728001; Roche) and subsequently treated with 1 μl 25 mM EDTA at 70 °C for 10 minutes to inactivate DNase I. cDNA synthesis was primed by addition of 1 μl gene-specific primer targeting the GFP ORF (10 µM, MT165) and 1 μl dNTPs (10 mM each) followed by incubation at 65 °C for 5 minutes. Then, the reverse transcription reaction was set up by adding 20 units RiboLock RNase inhibitor (#EO0381; ThermoFisher Scientific), 200 units of Maxima reverse transcriptase (#EP0743; ThermoFisher Scientific, 4 μl of 5x Maxima reverse transcriptase buffer and 2.5 μl of nuclease-free water. The reaction was then incubated for 30 minutes at 50 °C followed by heat-inactivation at 85 °C for 5 minutes. 20 μl of cDNA were then PCR amplified (1′ 96 °C, 20x(15″ 96 °C, 15″ 60 °C, 15″ 72 °C)) in a 100 μl reaction using MyTaq Red mix and primers containing the Illumina S1 and p5 adapter (MT397) and the Illumina S2 and p7 adapter (MT164). To generate input plasmid DNA (pDNA) barcode counts that serve as normalization control, the plasmid library that was used for the transfections was linearized using EcoRI-HF and subsequently 1 ng of linearized vector was PCR amplified as before using 8 cycles. PCR products were pooled and purified by double-sided CleanPCR bead purification using beads:sample ratios of 0.6:1 followed by 1.2:1 on the supernatant. The sequencing library was then sequenced using a 75 bp single-read NextSeq High Output kit (Illumina), yielding on average ∼8.8×10^6^ reads per sample, and thus on average ∼248 reads per barcode.

### RNA-seq data generation and analysis

RNA-seq data was generated for mNPCs as following. 1×10^6^ mNPCs were collected on two separate days and resuspended in 600 µl RLT buffer (#79216, Qiagen). RNA was isolated using RNeasy column purification (#74104, Qiagen). Sequencing libraries were prepared using TruSeq polyA stranded mRNA library prep kit (#20020595, Illumina) and sequenced on a NovaSeq 6000 with 51 bp paired-end reads yielding 20×10^6^ reads per sample. RNA-seq data for mESCs was retrieved from public resources.^49^ Data for all other cell lines was collected from the Human Protein Atlas (https://www.proteinatlas.org/about/download, #25 - RNA HPA cell line gene data, The Human Protein Atlas version 23.0, Ensembl version 109). For all cell lines and all genes, transcripts per million (TPM) were calculated and then normalized to nTPM using Trimmed mean of M values^50^ to allow for between-sample comparisons. To compute correlations between TF reporter activity and TF expression, only TFs with differences in expression across cell lines were included (nTPM > 8 in at least one cell line, nTPM < 1 in at least one cell line). Additionally, TFs that were not active in any cell line (reporter activity (log_2_) < 0.75) were excluded. Several TFs were included in the analysis even though they did not pass these filters (STAT3, SP1, TEAD1, NFKB1, ZFX, NR4A1). In case of heterodimeric TFs (e.g., POU5F1::SOX2), we considered in each cell line the nTPM value of the TF with the lowest abundance, since this TF is the limiting factor of the heterodimer.

### Reporter activity computation and normalizations

Raw barcode counts were clustered using *starcode*^51^ using a maximum Levenshtein distance of 1. Next, clustered barcode counts were normalized by library size. To be more precise, the clustered barcode counts were divided by the total sum of all barcode counts per sample per million. From these normalized barcode counts activities were computed by dividing the cDNA barcode counts by the plasmid DNA barcode counts. The activities were normalized by dividing the activities by the median of the activities of the TF-neg reporters per core promoter and sample. Normalized activities were then averaged over the different barcodes and finally over the independent replicates per condition.

### Log-linear model of reporter activities

To explore the impact of the reporter design on the reporter activity, for each TF a log-linear model was fit using the following equation.

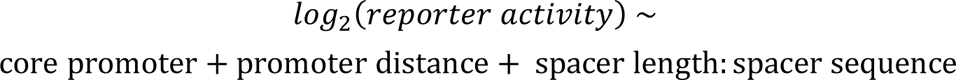

The reporter activities were fit for each TF in three different conditions where the TF is a) expressed highest, or b) stimulated or overexpressed (if data available). We reasoned that these conditions would represent the most TF-specific conditions. The condition with the best model performance was chosen as representative model for the TF and is displayed in **Figure 2D**. See **Table S2** for chosen reference conditions. All input features in the model were used as categorical variables. Models were fit using the lm function in R from the stats package (version 3.6.2).

### Reporter confidence level and reporter score computation

To evaluate the performance of each individual TF reporter, reporter confidence levels were computed as mentioned in the Results section. In case more than one perturbation condition was tested for a TF, the perturbation with the strongest average reporter activity fold-change was selected (conditions denoted by asterisk in **Figure 4A** & **Figure 5A**). The same selection was done in case of multiple off-target TF perturbation conditions. TF abundance correlation was only taken into consideration for TFs that were included in the TF abundance correlation analysis (see **Figure 3A**, “*RNA-seq data generation and analysis*” section). Moreover, to rank reporters within a confidence level, a reporter quality score was computed as follows.

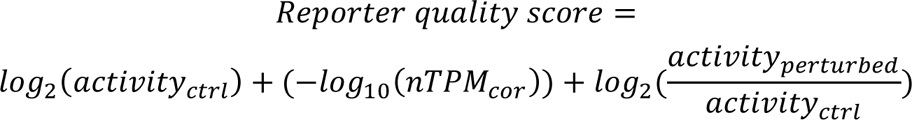

where *nTPM_cor_* refers to the correlation of the reporter activities with the TF transcript abundance across the nine tested cell lines, and *activity_ctrl_* refers to the selected reference condition mentioned in the “*Log-linear model of reporter activities*” section.

## SUPPLEMENTARY FIGURES

**Figure S1:**
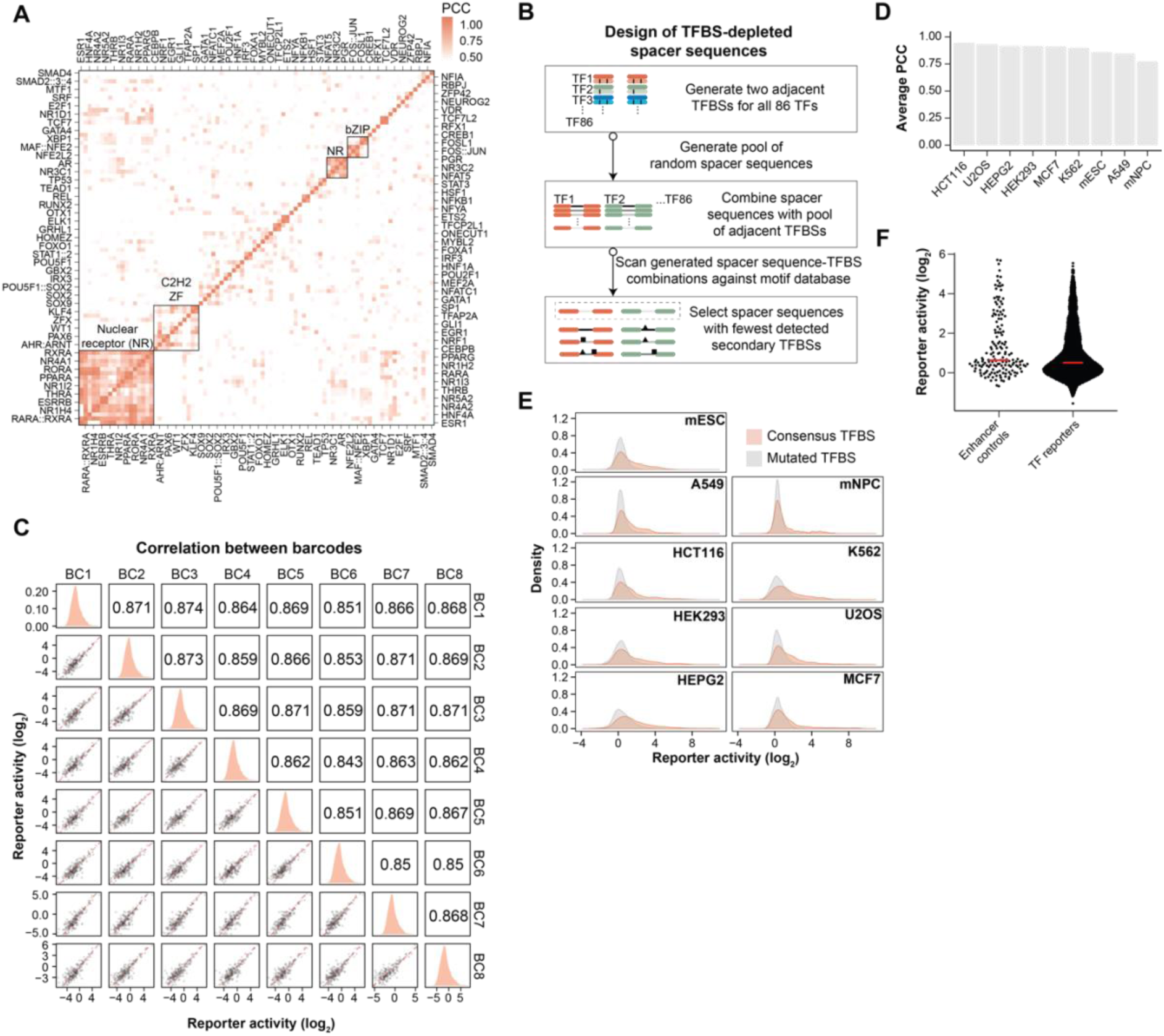
TF reporter library design and activity characterization. (**A**) Pairwise Pearson correlation coefficient (PCC) heatmap for all 86 motifs chosen for the TF reporter library design. Motifs clustering together are highlighted. (**B**) Schematic overview of the design of the motif-depleted spacer sequences. (**C**) Correlations between the reporter activities of individual barcodes. Displayed are 300 randomly sampled reporters. Figure panels on the diagonal show the density distribution of the reporter activities per barcode. Figure panels below the diagonal show the pairwise correlation plots, and panels above the diagonal indicate the PCCs. (**D**) Average PCC of all pairwise correlations between biological replicates per cell line. (**E**) Reporter activity distributions per cell line of reporters with mutated TFBSs (grey) and consensus TFBSs (red). (**F**) Comparison of reporter activities of TF reporters and the genomic *Klf2* gene enhancer controls in mESCs. Red line indicates median reporter activity.

**Figure S2:**
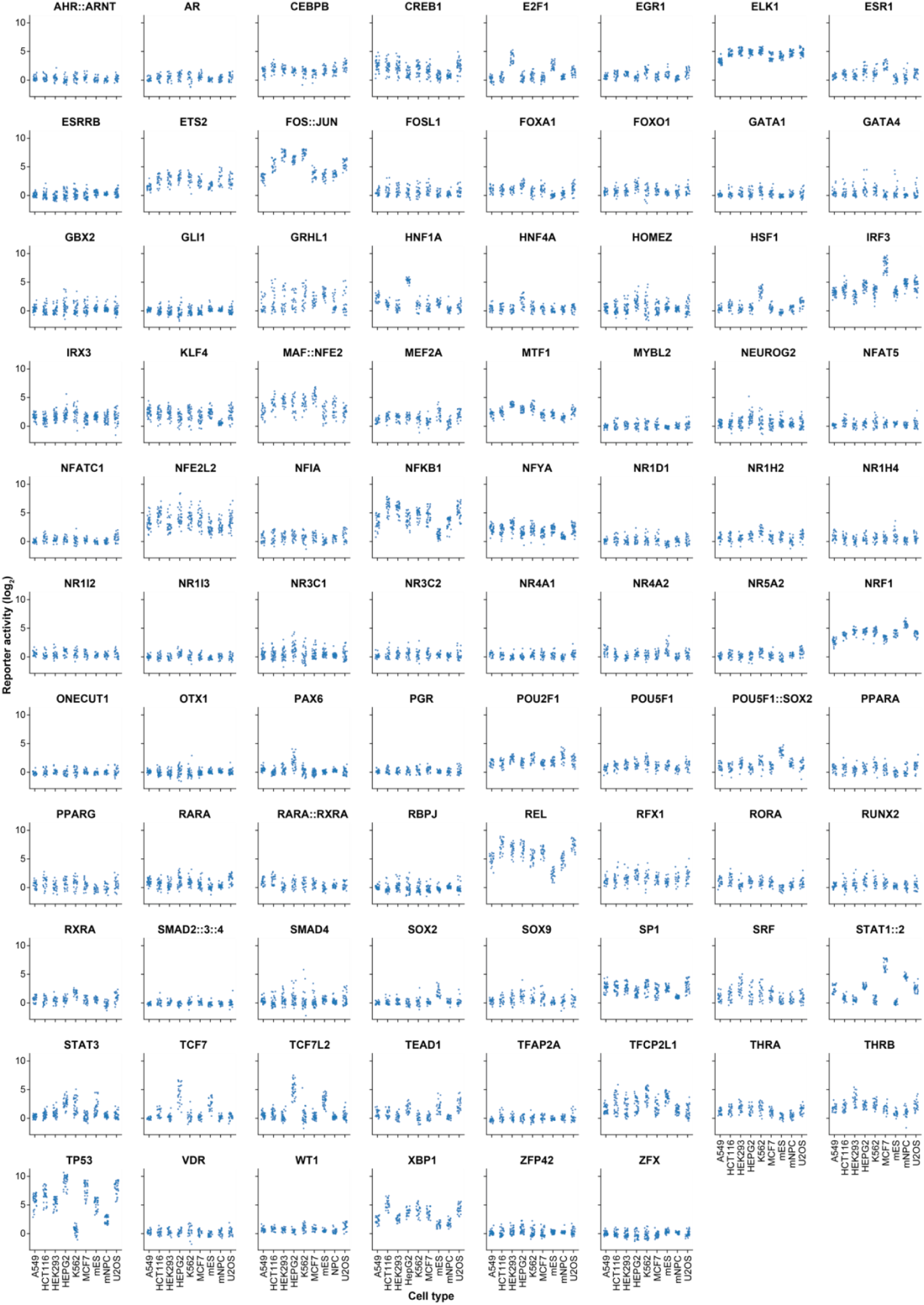
TF reporter activities across all probed cell lines. Reporter activities per TF in all nine probed cell lines. Each dot represents a unique reporter design.

**Figure S3:**
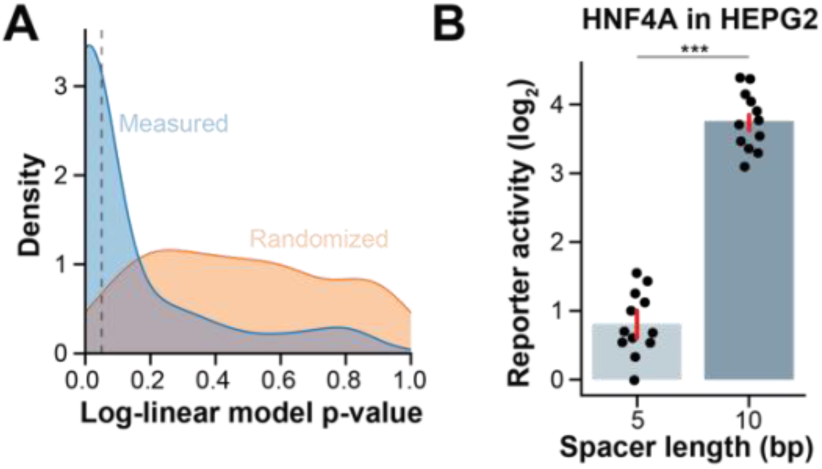
Log-linear models highlight importance of reporter design. (**A**) Distribution of the p-values of all log-linear models (one per TF) with measured data (blue) and randomized data (orange; measured activities were randomly assigned to reporter designs per TF). (**B**) HNF4A reporter activities per spacer length. The bar indicates the mean reporter activity per spacer length, the dots indicate activities of individual reporters. The red line denotes the standard deviation. Difference in activity between the groups is tested by student’s t-test; *** < 0.001.

**Figure S4:**
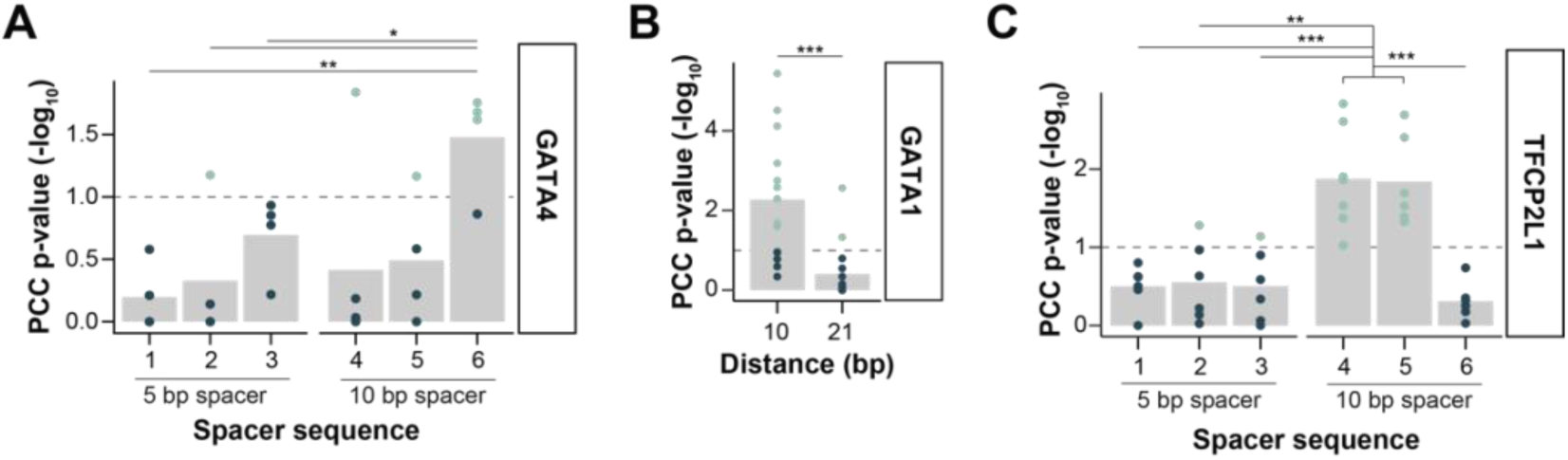
Reviewing TF specificity of reporters. (**A**) PCC p-value of GATA4 reporters per spacer sequence and spacer length. Mean is indicated by the bar, and individual dots represent individual reporters. Significant p-values (p < 0.1) are indicated by green color. (**B**) Same as (**A**) but for GATA1 reporters per promoter distance. (**C**) Same as (**A**) but for TFCP2L1 reporters. Difference in PCC p-value between the groups is tested by student’s t-test; * < 0.05, ** < 0.01, *** < 0.001.

**Figure S5:**
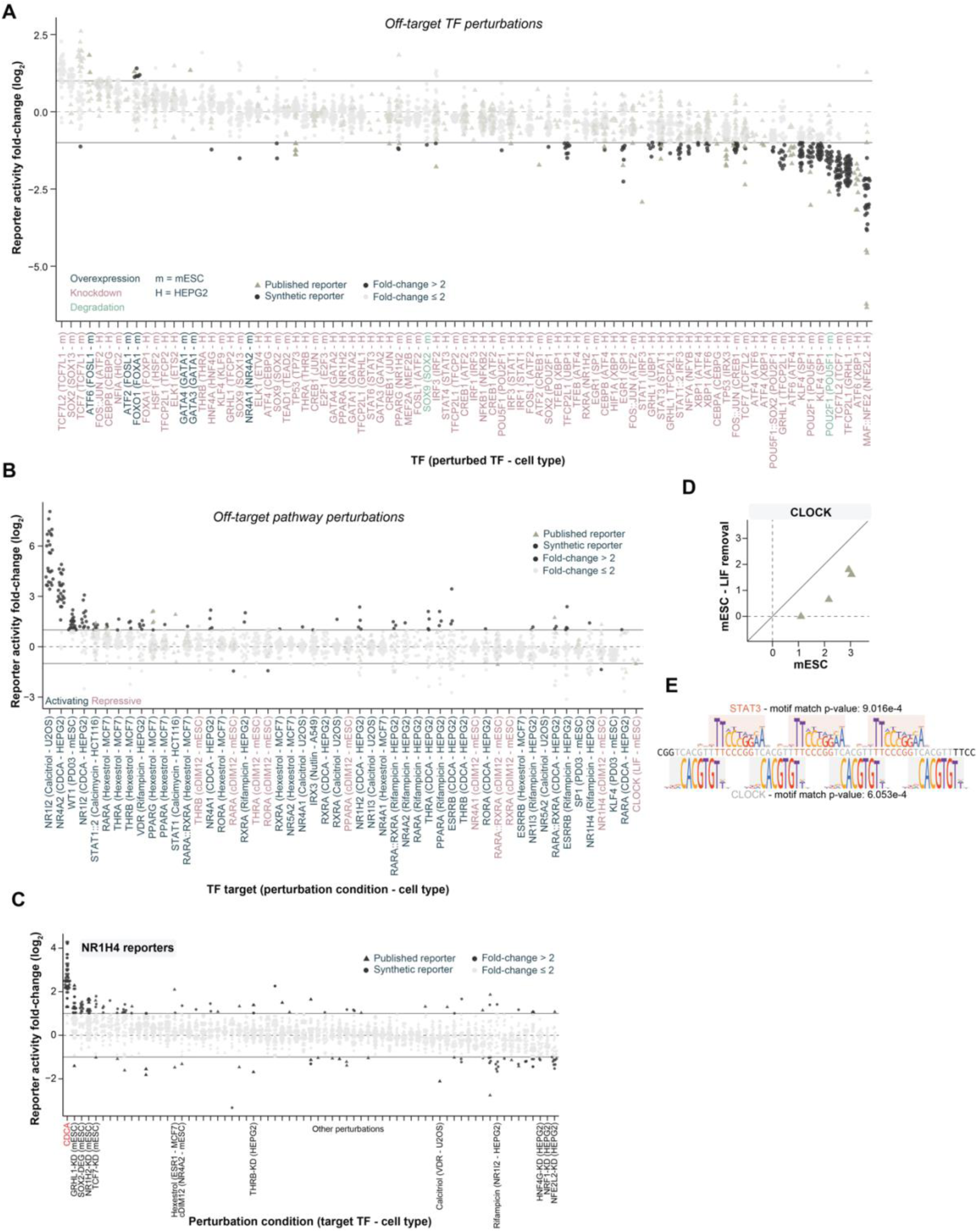
Reviewing TF specificity of reporters. (**A**) Changes in reporter activities upon perturbation of TFs with similar TFBSs by either TF overexpression (black), knockdown (purple), or degradation (green). (**B**) Same as **A** but for pathway perturbations of TFs with similar TFBSs. (**C**) Changes in reporter activity of NR1H4 reporters upon all perturbations. Only top changing conditions and conditions that perturb NR1H4 or TFs with similar TFBSs are indicated. (**D**) Activity of published CLOCK reporters in mESCs and mESCs cultured in the absence of LIF. (**E**) Sequence of the published CLOCK reporter and motif energy logos of STAT3 (red; motif ID: STAT3.H12CORE.0.P.B) and CLOCK (grey; motif ID: CLOCK.H12CORE.1.PS.A). Motif match p-values were computed using MoLoTool.^52^

**Figure S6:** Reporter confidence level heatmaps for all TFs (external file). Upper heatmap: confidence levels per reporter. Middle heatmap: Activities (first row), TF abundance correlation (second row), perturbation fold-change (third row), and off-target perturbation fold-change (fourth row) per reporter. Lower heatmap: Color-coding of the reporter design.

**Figure S7:**
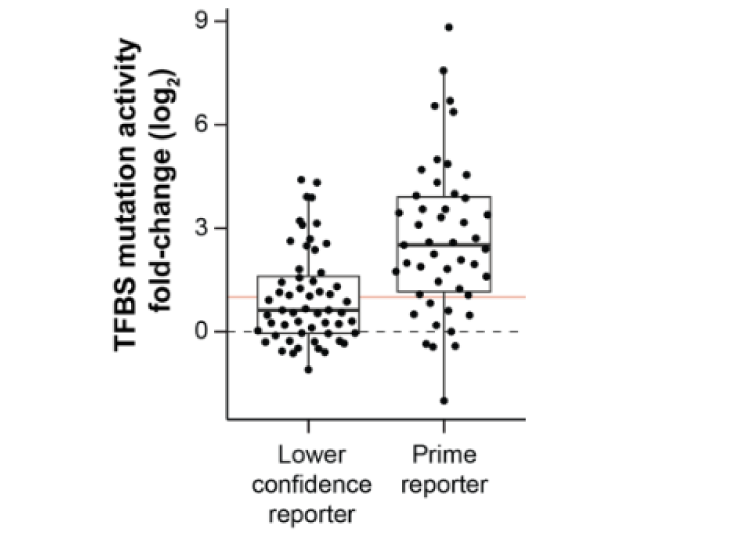
Identifying high-confidence TF reporters. Change in reporter activity upon TFBS mutation of the 47 prime reporters with matched mutated reporters compared to reporters for the same TFs with a lower confidence level (mean across all reporters with one confidence level lower than the prime reporter). Red line indicates a fold-change of 2.

**Figure S8:**
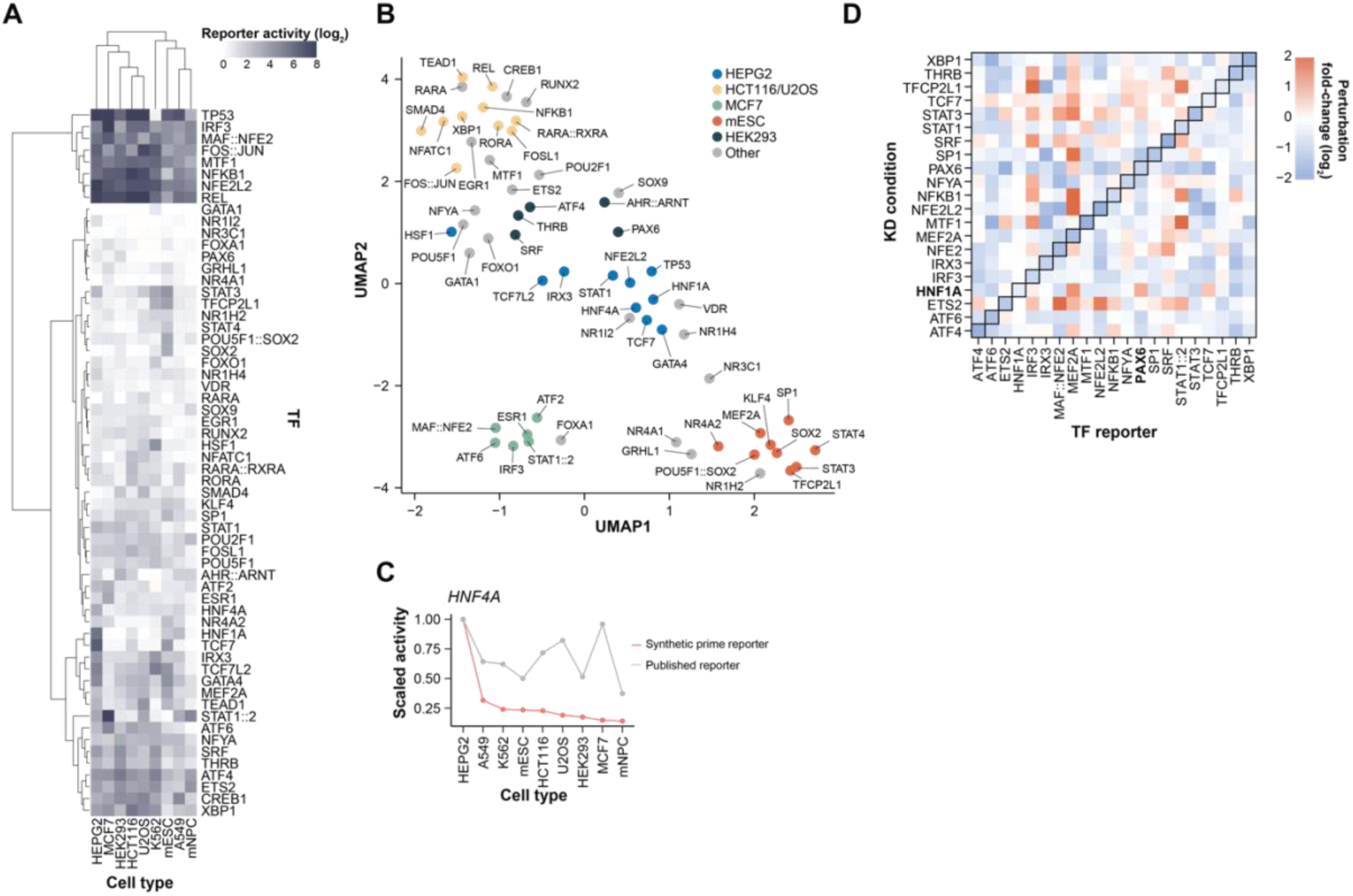
Perturbation responses of high-confidence TF reporters. (**A**) Activities of the 60 prime reporters in the nine probed cell lines. (**B**) Data shown in Figure 7A visualized as UMAP. Color codes are based on clustering in Figure 7A. (**C**) Reporter activities (max-normalized) of the synthetic HNF4A prime reporter (red) and the highest-ranking published HNF4A reporter per cell line. (**D**) Changes in TF activities upon KD of TFs in HEPG2 cells. KDs that did not reduce the activity of their target TF (log_2_-fc > -0.25) are not included in this visualization. Example mentioned in the text is highlighted in bold.

## DATA AVAILABILITY

Laboratory notes and supplementary raw data are available at Zenodo (https://doi.org/10.5281/zenodo.11199257). Code and analysis pipelines are available at GitHub (https://github.com/mtrauernicht/TF_MPRA). A released version of the GitHub repository is available at Zenodo (https://doi.org/10.5281/zenodo.11203837). RNA-seq of the mNPCs is available at GEO under accession number GSE267969. Raw sequencing data of the RNA-seq and all MPRAs is available at SRA under accession number PRJNA1112759.

## ACKNOWLEDGMENTS

We thank members of our laboratories for helpful comments; the NKI Genomics and Research High-Performance Computing core facilities for technical support. We also want to thank Wangjie Liu, Antoni Gralak and Bart Deplancke (EPFL, Lausanne, Switzerland) for sharing lentiviral plasmids used for the TF overexpression experiments. We also thank Harmen J. Bussemaker (Columbia University, New York, USA) for discussions and feedback during the design of the reporter library. This work was funded by the Oncode Institute and the European Union (European Research Council Advanced Grant RE_LOCATE, 101054449), and the National Institutes of Health (NIMH Grant R01MH106842). Views and opinions expressed are however those of the author(s) only and do not necessarily reflect those of the European Union or the European Research Council. Neither the European Union nor the granting authority can be held responsible for them. Research at the Netherlands Cancer Institute is supported by an institutional grant of the Dutch Cancer Society and of the Dutch Ministry of Health, Welfare and Sport. The Oncode Institute is partially funded by the Dutch Cancer Society.

## AUTHOR CONTRIBUTIONS

Reporter library design: M.T. with input from C.R.; experiments and data analysis: M.T. with help from T.F.; manuscript writing: M.T. and B.vS.; Project supervision: B.vS.

