## Supplementary material for "Optimized reporters for multiplexed detection of transcription factor activity": Figure S6

### EXAMPLE FIGURE

TF = A

Perturbation condition = B

Off-target perturbation condition = C

Cell type = D

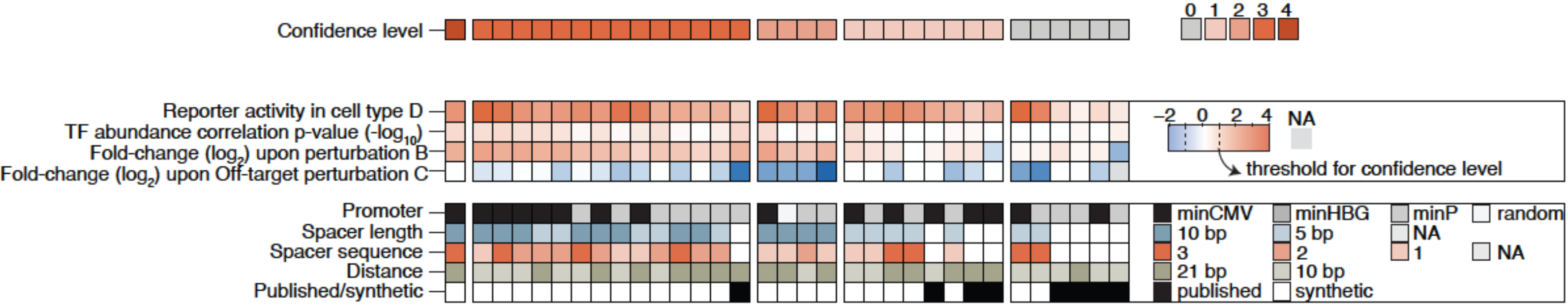

**TF** = AHR::ARNT

**Perturbation condition** = ITE (HCT116)

**Off-target perturbation condition** = NA

**Cell type** = mESC

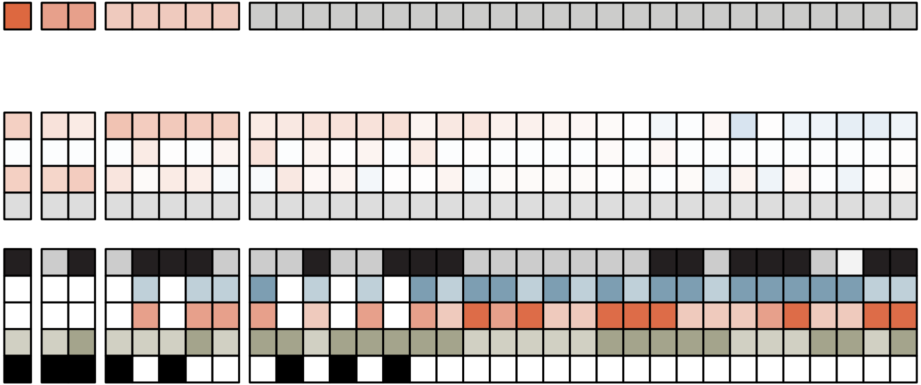

**TF** = AR

**Perturbation condition** = NA

**Off-target perturbation condition** = NA

**Cell type** = U2OS

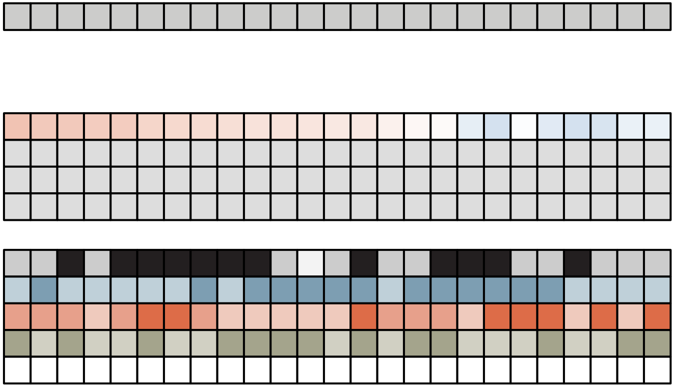

**TF** = CEBPB

**Perturbation condition** = NA

**Off-target perturbation condition** = ATF4-KD (HEPG2)

**Cell type** = mESC

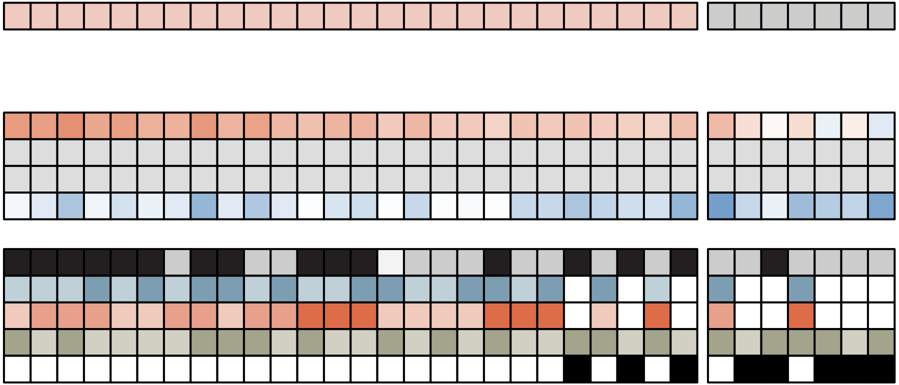

**TF** = CREB1

**Perturbation condition** = FK (mESC)

**Off-target perturbation condition** = ATF2-KD (HEPG2)

**Cell type** = U2OS

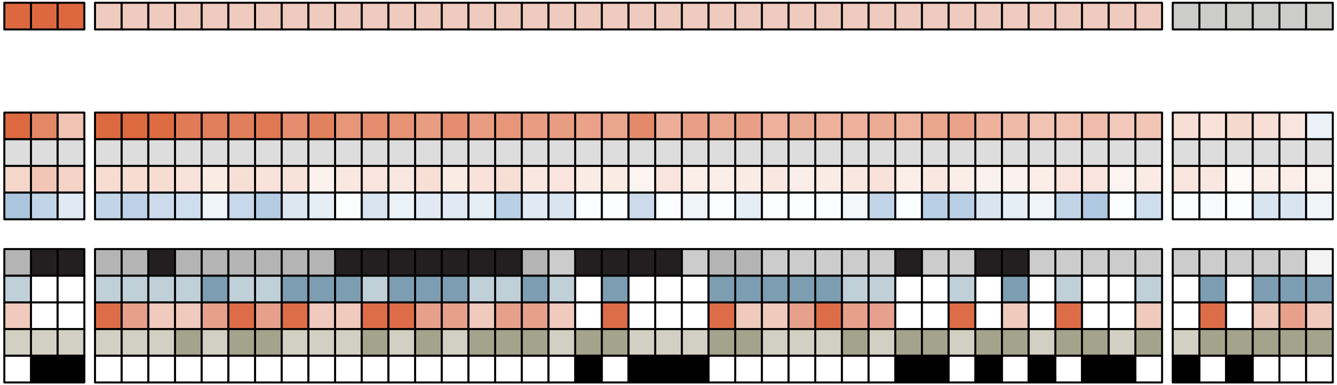

**TF** = E2F1

**Perturbation condition** = E2F1-KD (mESC)

**Off-target perturbation condition** = E2F3-KD (mESC)

**Cell type** = mESC

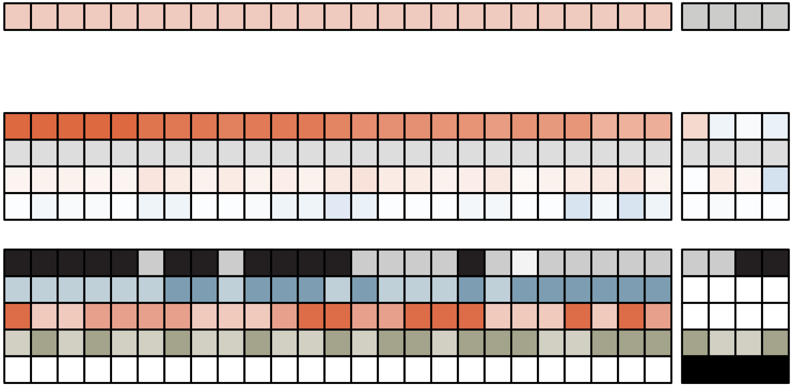

**TF** = EGR1

**Perturbation condition** = PD removal (mESC)

**Off-target perturbation condition** = SP1-KD (mESC)

**Cell type** = mESC

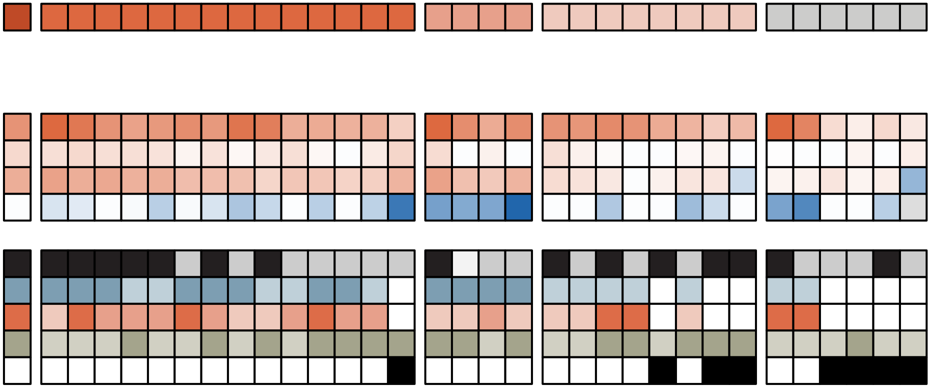

**TF** = ELK1

**Perturbation condition** = PD removal (mESC)

**Off-target perturbation condition** = ETV4-KD (HEPG2)

**Cell type** = U2OS

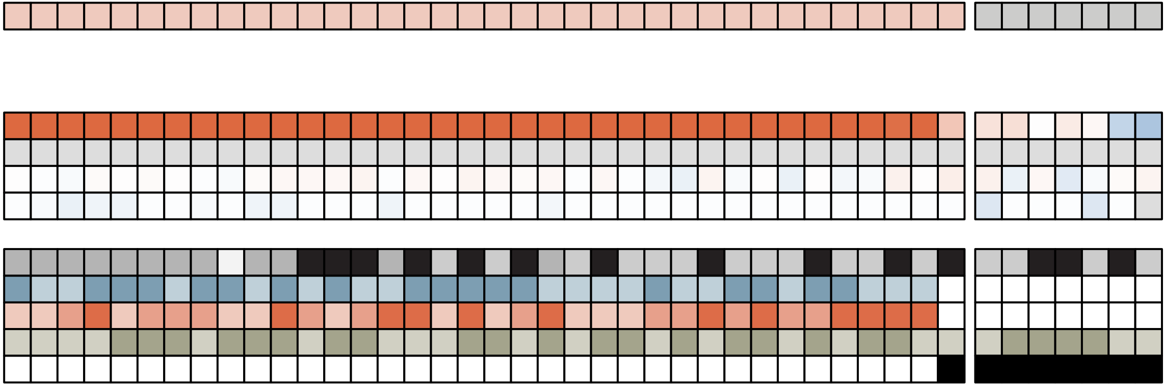

**TF** = ESR1

**Perturbation condition** = Hexestrol (MCF7)

**Off-target perturbation condition** = NA

**Cell type** = MCF7

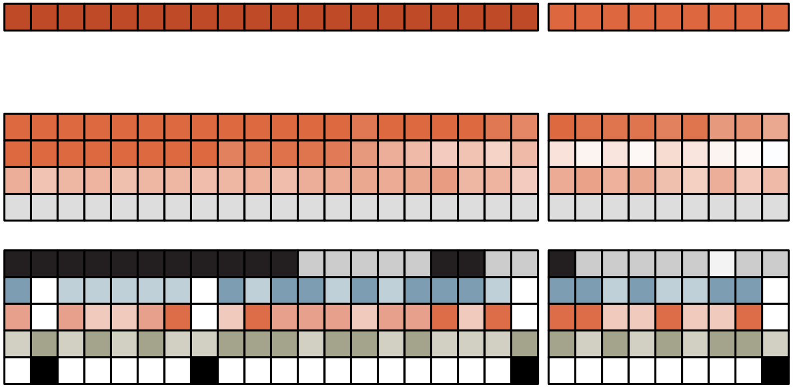

**TF** = ESRRB

**Perturbation condition** = NA

**Off-target perturbation condition** = Rifampicin (HEPG2)

**Cell type** = mESC

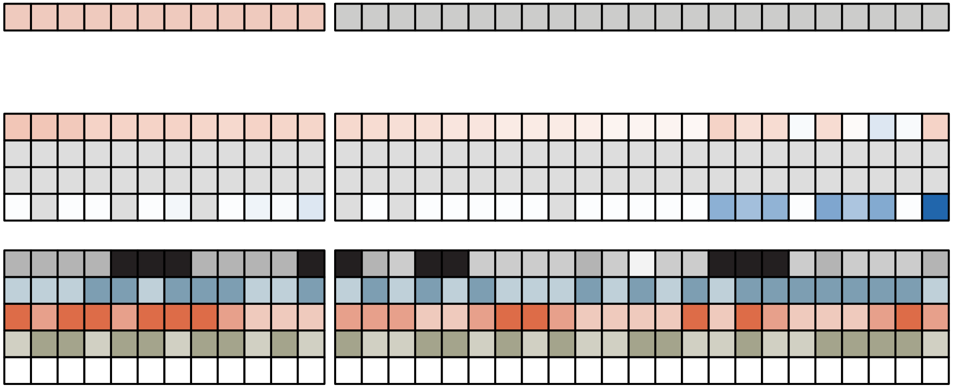

**TF** = ETS2

**Perturbation condition** = ETS2-KD (HEPG2)

**Off-target perturbation condition** = NA

**Cell type** = mESC

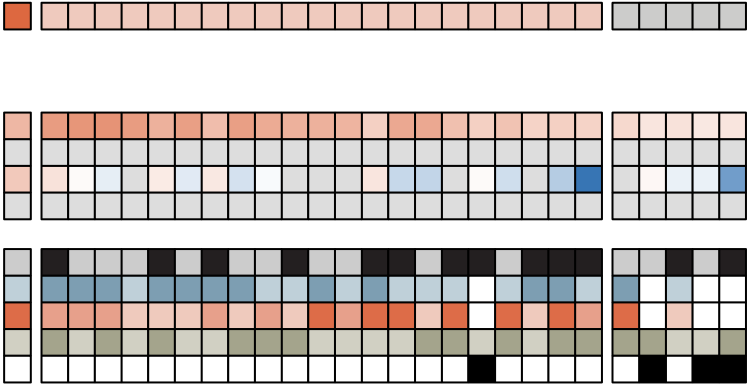

**TF** = FOS::JUN

**Perturbation condition** = PD removal (mESC)

**Off-target perturbation condition** = CREB1-KD (mESC)

**Cell type** = U2OS

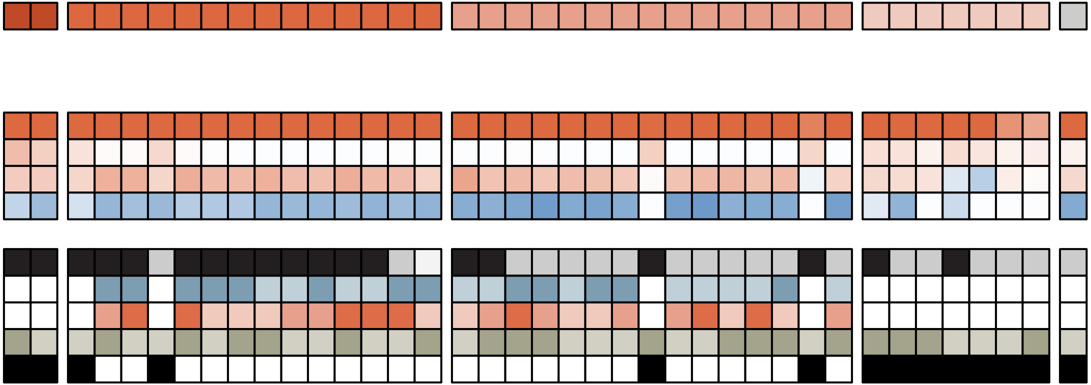

**TF** = FOSL1

**Perturbation condition** = FOSL1 overexpression (mESC)

**Off-target perturbation condition** = ATF2-KD (HEPG2)

**Cell type** = HCT116

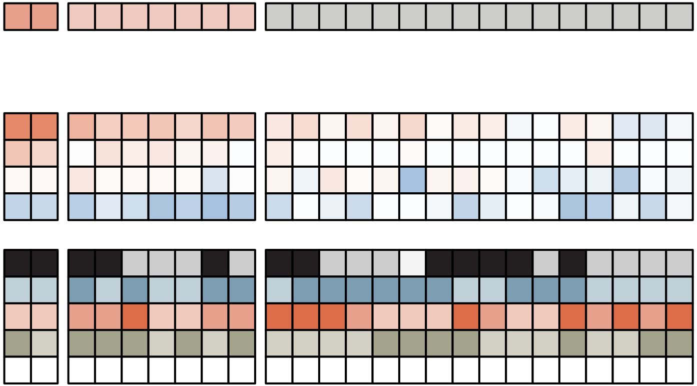

**TF** = FOXA1

**Perturbation condition** = FOXA1 overexpression (mESC)

**Off-target perturbation condition** = FOXP1-KD (HEPG2)

**Cell type** = mESC

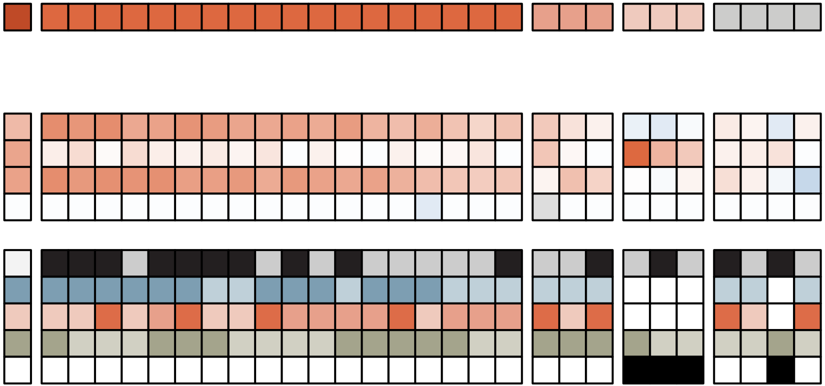

**TF** = FOXO1

**Perturbation condition** = Wortmannin (U2OS)

**Off-target perturbation condition** = FOXA1 overexpression (mESC)

**Cell type** = mESC

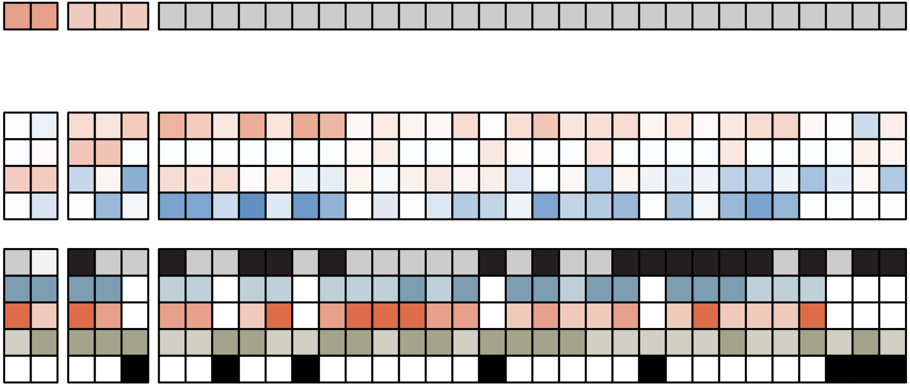

**TF** = GATA1

**Perturbation condition** = GATA1 overexpression (mESC)

**Off-target perturbation condition** = GATA2-KD (HEPG2)

**Cell type** = K562

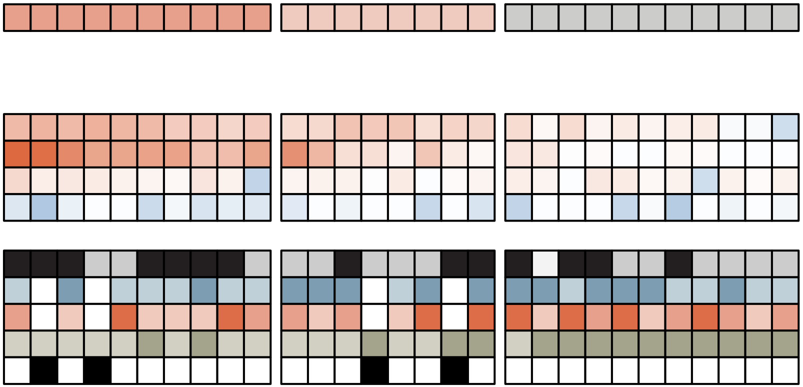

**TF** = GATA4

**Perturbation condition** = NA

**Off-target perturbation condition** = GATA1 overexpression (mESC)

**Cell type** = HEPG2

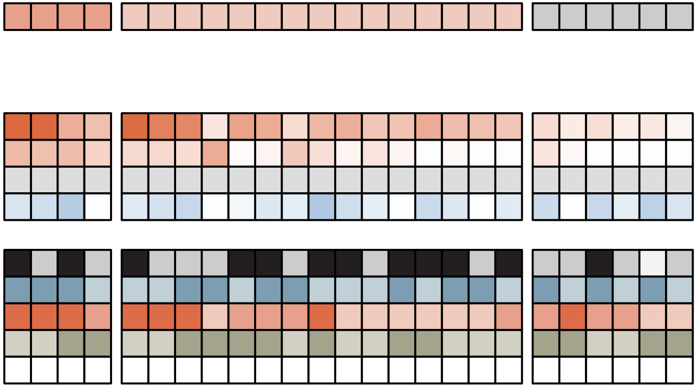

**TF** = GBX2

**Perturbation condition** = NA

**Off-target perturbation condition** = NA

Cell type = mESC

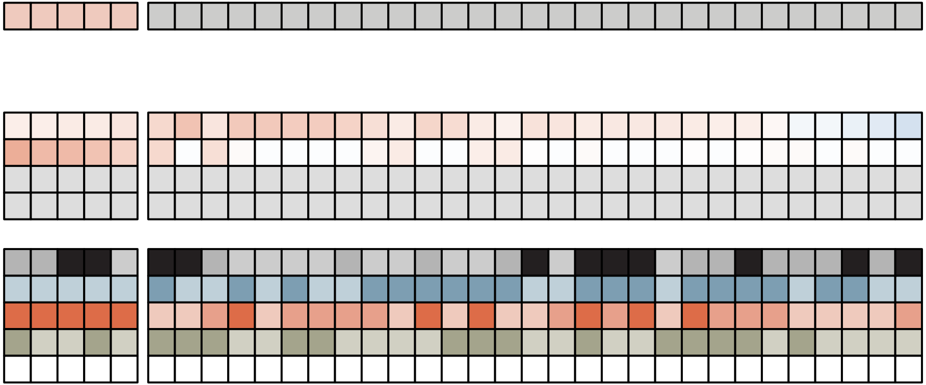

**TF** = GLI1

**Perturbation condition** = NA

**Off-target perturbation condition** = NA

**Cell type** = mNPC

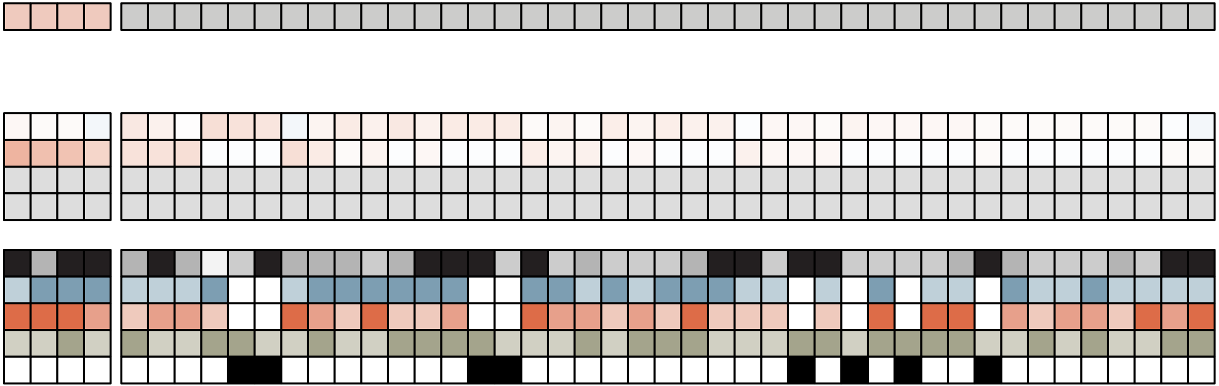

**TF** = GRHL1

**Perturbation condition** = GRHL1-KD (mESC)

**Off-target perturbation condition** = TFCP2L1-KD (mESC)

**Cell type** = mESC

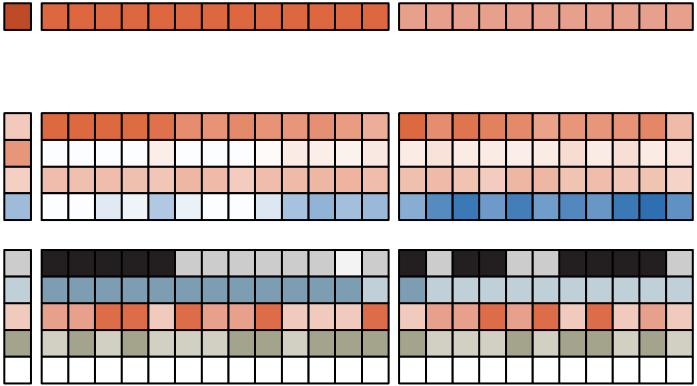

**TF** = HNF1A

**Perturbation condition** = HNF1A-KD (HEPG2)

**Off-target perturbation condition** = NA

**Cell type** = HEPG2

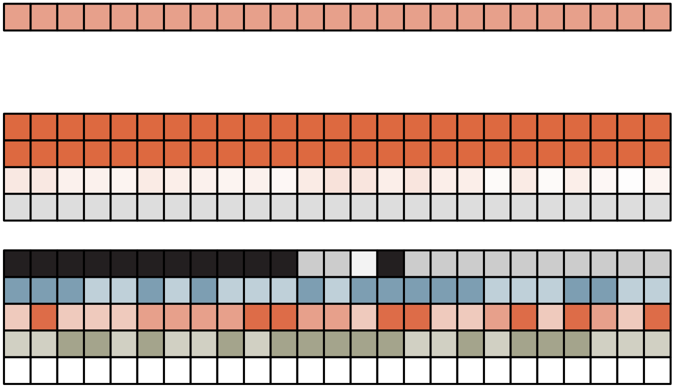

**TF** = HNF4A

**Perturbation condition** = NA

**Off-target perturbation condition** = HNF4G-KD (HEPG2)

**Cell type** = HEPG2

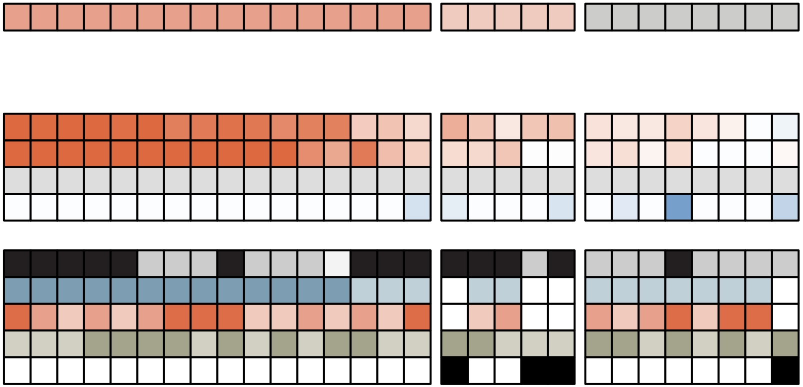

**TF** = HOMEZ

**Perturbation condition** = NA

**Off-target perturbation condition** = NA

**Cell type** = HEPG2

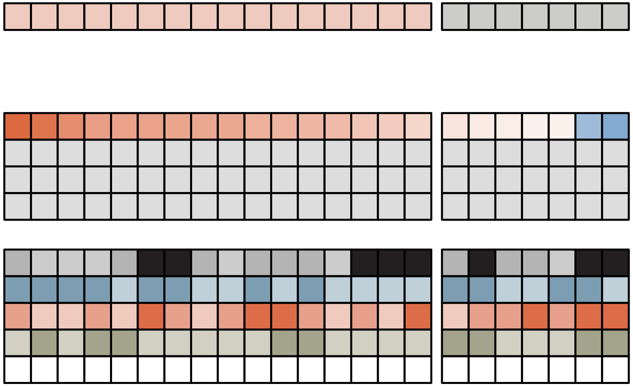

**TF** = HSF1

**Perturbation condition** = Heat shock (U2OS)

**Off-target perturbation condition** = HSF2-KD (mESC)

**Cell type** = U2OS

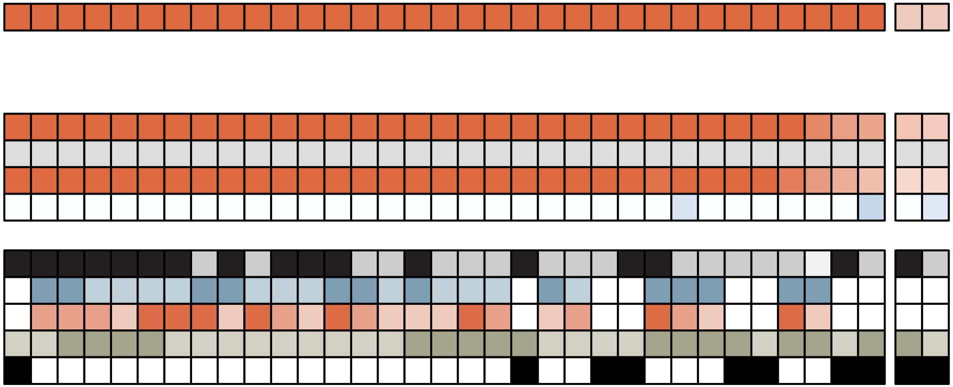

**TF =** IRF3

**Perturbation condition =** IRF3-KD (HEPG2)

**Off-target perturbation condition =** STAT1-KD (HEPG2)

**Cell type =** MCF7

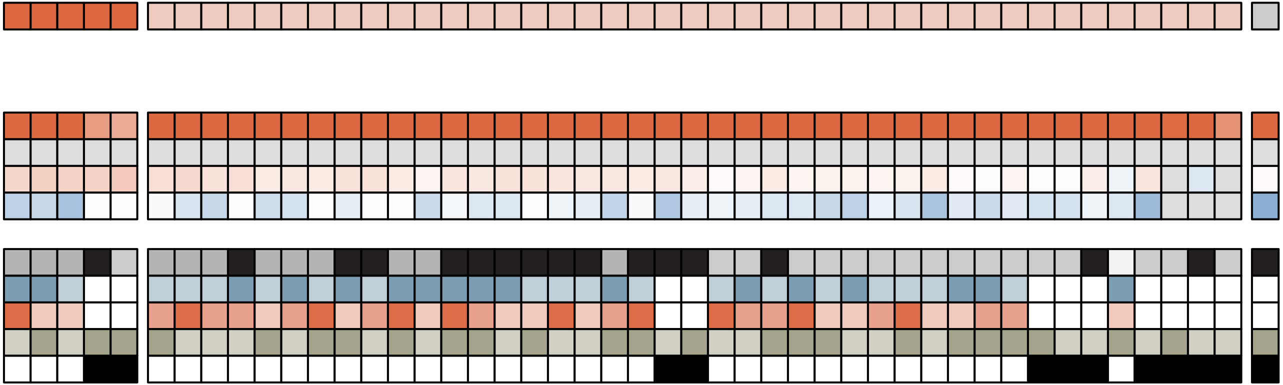

**TF** = IRX3

**Perturbation condition** = IRX3-KD (HEPG2)

**Off-target perturbation condition** = Nutlin-3a (A549)

**Cell type** = HEPG2

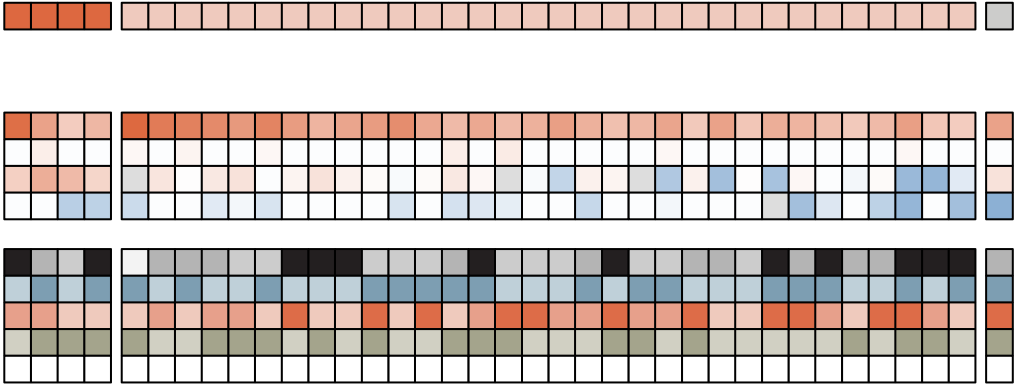

**TF** = KLF4

**Perturbation condition** = NA

**Off-target perturbation condition** = SP1-KD (mESC)

**Cell type** = mESC

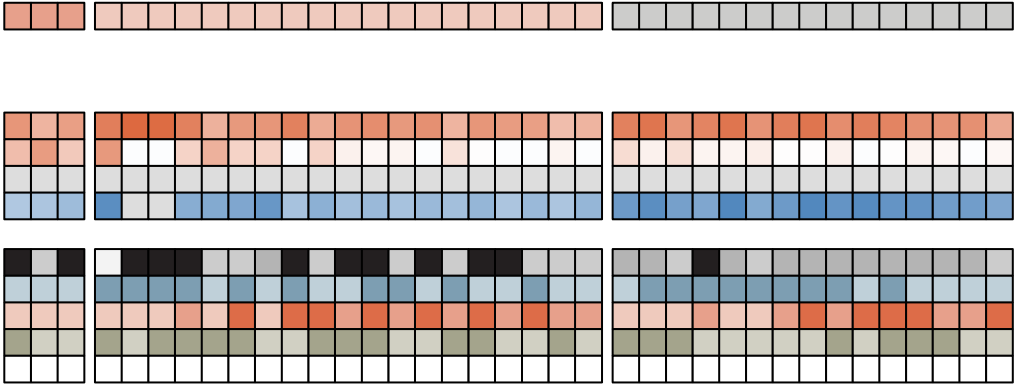

**TF** = MAF::NFE2

**Perturbation condition** = NFE2-KD (HEPG2)

**Off-target perturbation condition** = NFE2L2-KD (HEPG2)

**Cell type** = K562

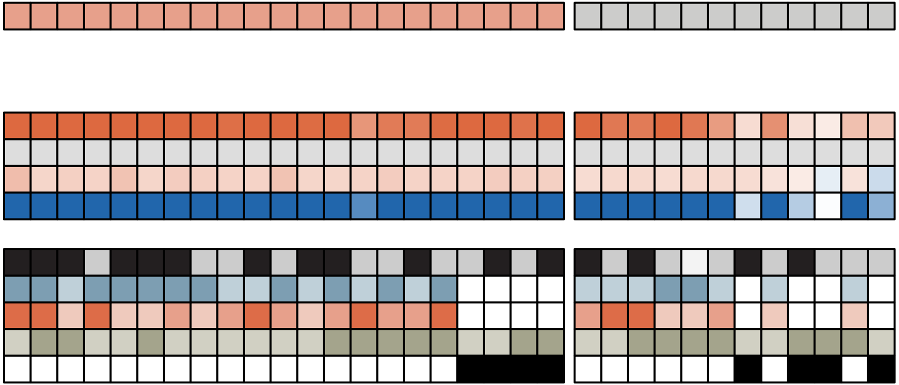

**TF** = MEF2A

**Perturbation condition** = MEF2A-KD (mESC)

**Off-target perturbation condition** = MEF2B-KD (mESC)

**Cell type** = U2OS

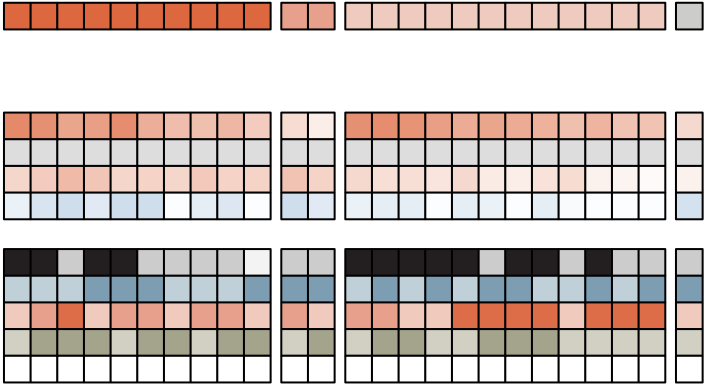

**TF** = MTF1

**Perturbation condition** = MTF1-KD (HEPG2)

**Off-target perturbation condition** = NA

**Cell type** = mESC

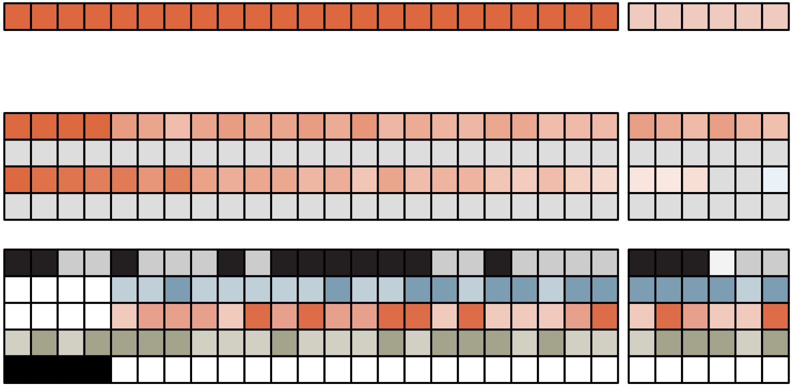

**TF** = MYBL2

**Perturbation condition** = NA

**Off-target perturbation condition** = NA

**Cell type** = mESC

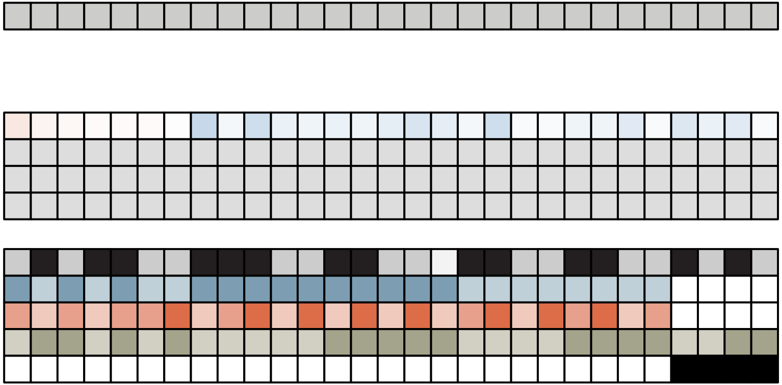

**TF** = NEUROG2

**Perturbation condition** = NA

**Off-target perturbation condition** = NA

**Cell type** = HEPG2

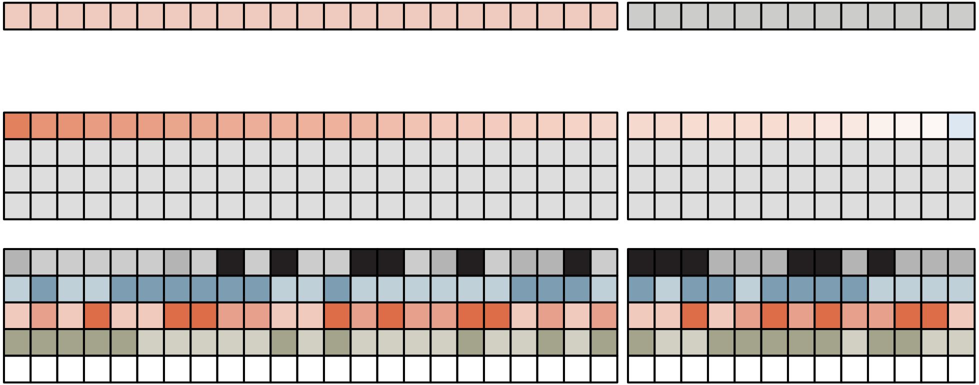

**TF** = NFAT5

**Perturbation condition** = Calcimycin (HCT116)

**Off-target perturbation condition** = NA

**Cell type** = mESC

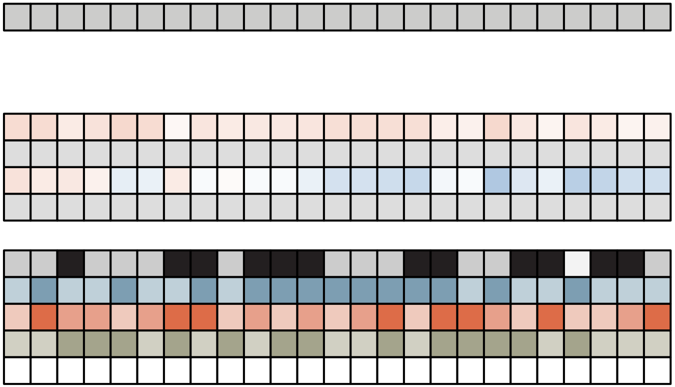

**TF** = NFATC1

**Perturbation condition** = Calcimycin (HCT116)

**Off-target perturbation condition** = NA

**Cell type** = HCT116

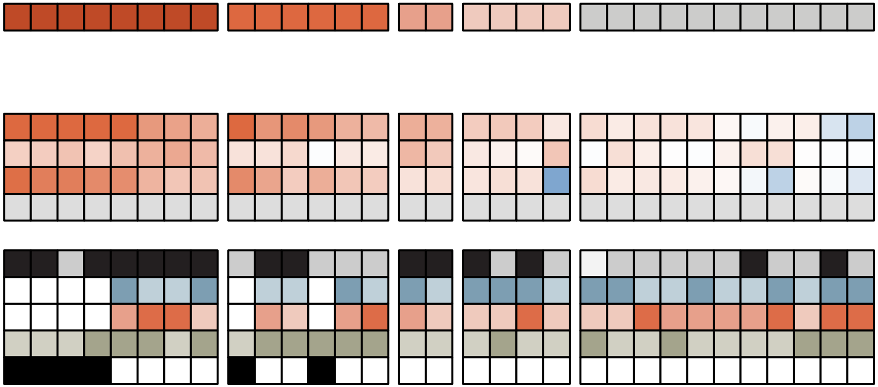

**TF** = NFE2L2

**Perturbation condition** = HQ (mESC)

**Off-target perturbation condition** = NA

**Cell type** = mESC

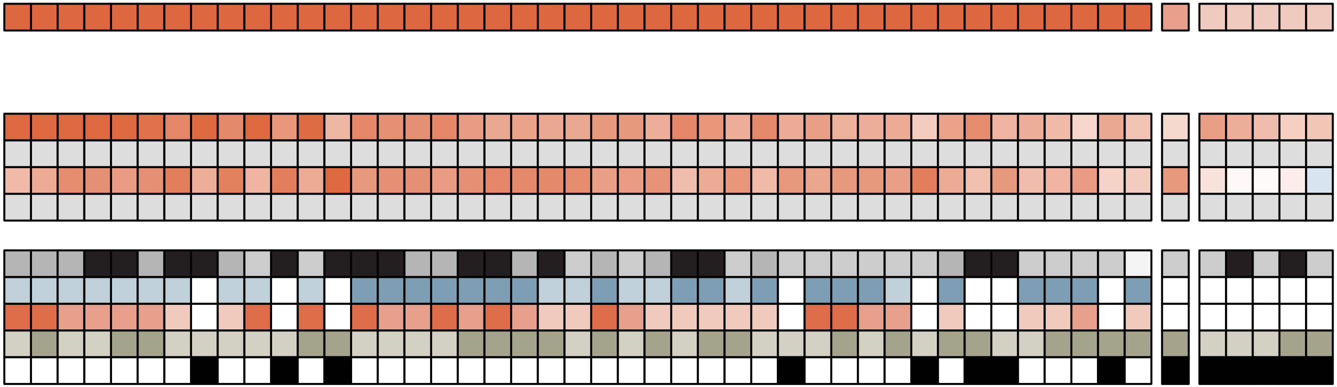

**TF** = NFIA

**Perturbation condition** = NA

**Off-target perturbation condition** = HIC2-KD (mESC)

**Cell type** = HCT116

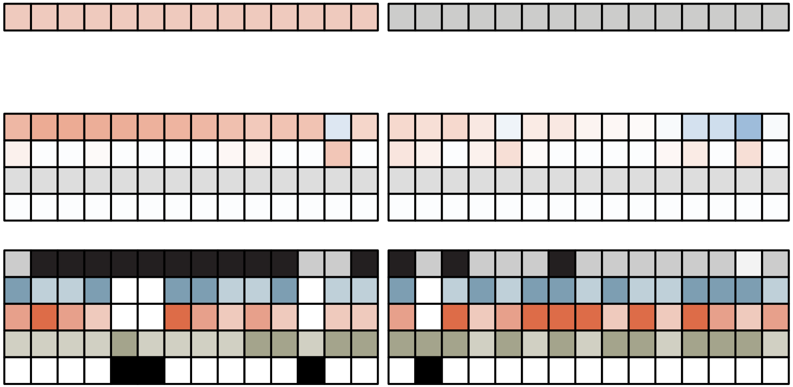

**TF** = NFKB1

**Perturbation condition** = PMA (U2OS)

**Off-target perturbation condition** = NFKB2-KD (HEPG2)

**Cell type** = U2OS

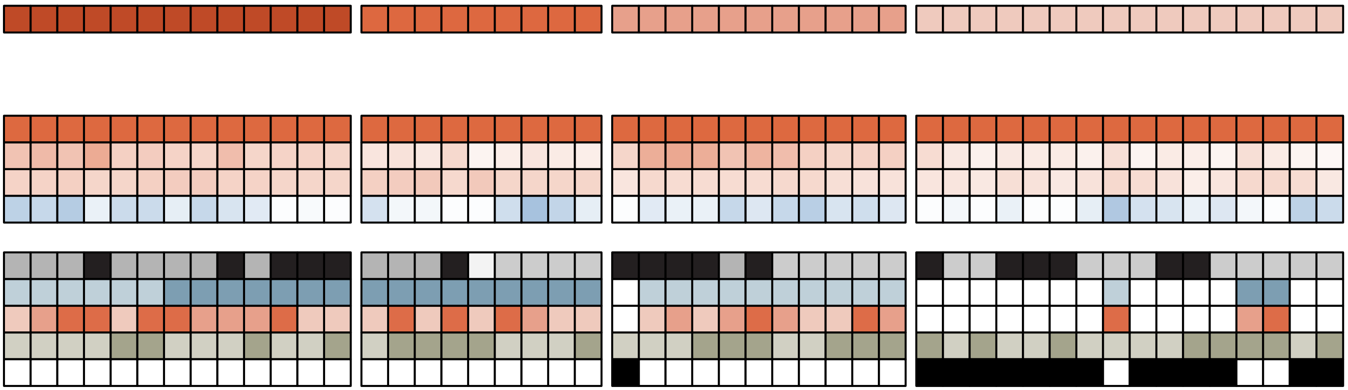

**TF** =  NFYA

**Perturbation condition** = NFYA-KD (HEPG2)

**Off-target perturbation condition** =  NFYB-KD (HEPG2)

**Cell type** =  U2OS

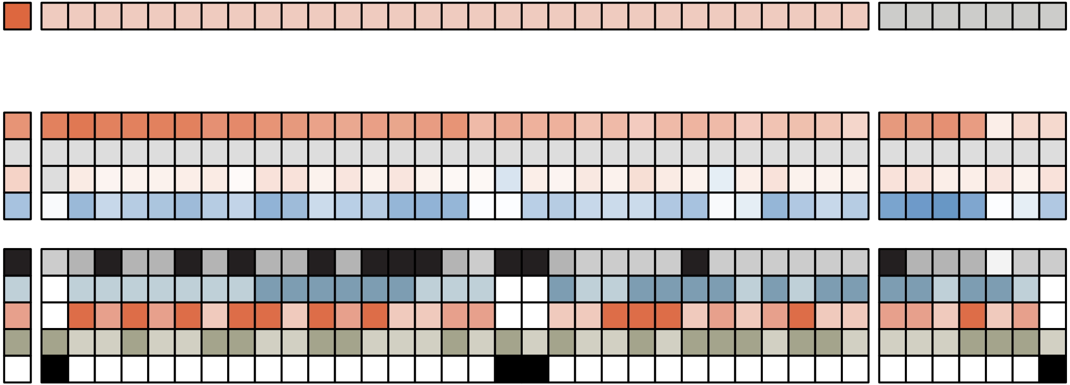

**TF** = NR1D1

**Perturbation condition** = LIF removal (mESC)

**Off-target perturbation condition** = NA

**Cell type** = HCT116

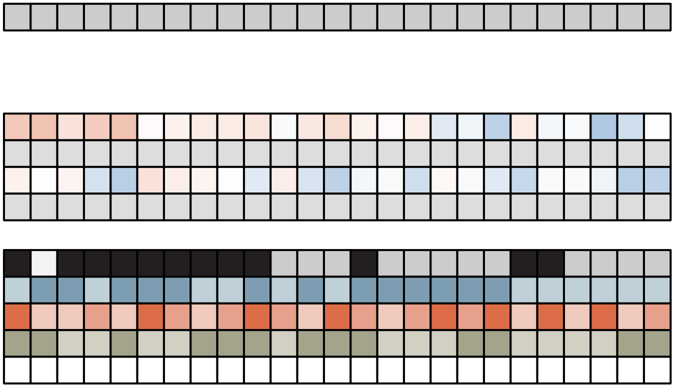

**TF** = NR1H2

**Perturbation condition** = NR1H2-KD (mESC)

**Off-target perturbation condition** = CDCA (HEPG2)

**Cell type** = mESC

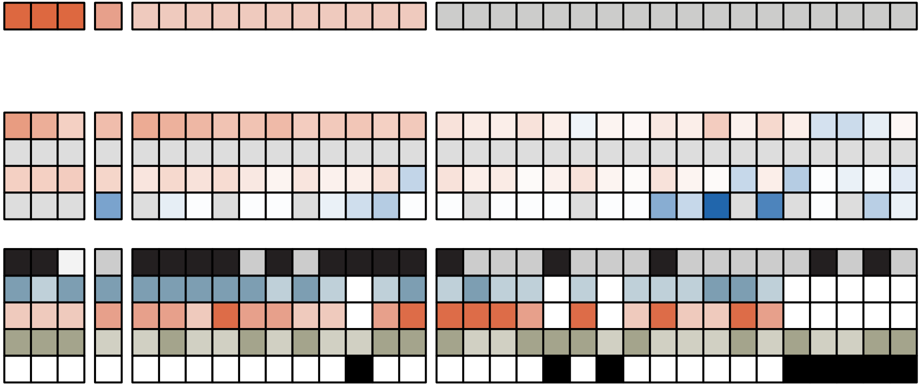

**TF** = NR1H4

**Perturbation condition** = CDCA (HEPG2)

**Off-target perturbation condition** = NR4A2 overexpression + cDIM12 (mESC)

**Cell type** = HEPG2

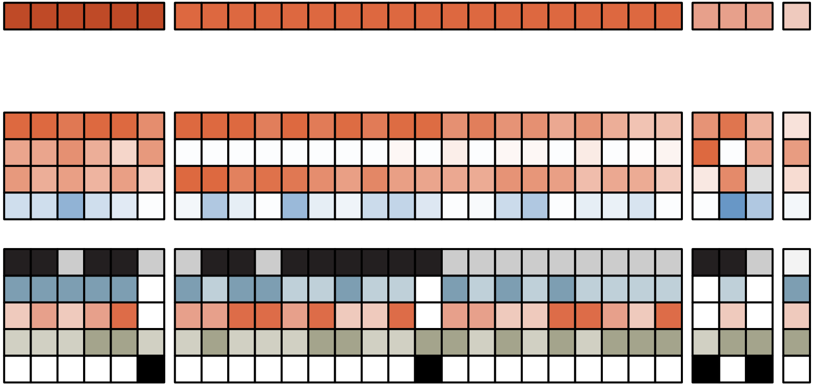

**TF** = NR1I2

**Perturbation condition** = Rifampicin (HEPG2)

**Off-target perturbation condition** = Calcitriol (U2OS)

**Cell type** = HEPG2

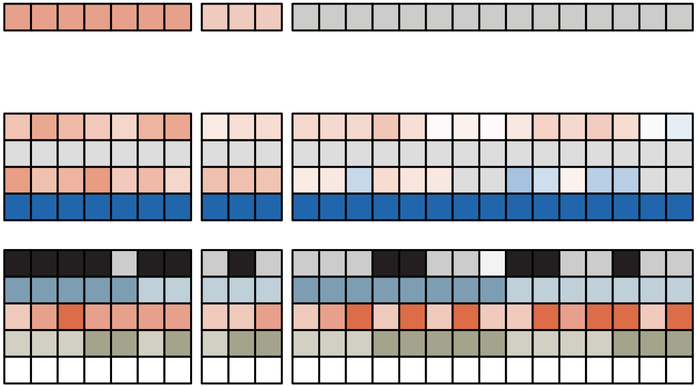

**TF** = NR1I3

**Perturbation condition** = NA

**Off-target perturbation condition** = Calcitriol (U2OS)

**Cell type** = HEPG2

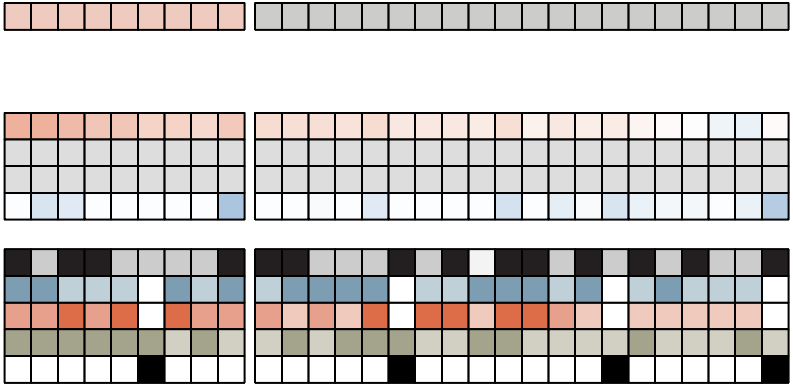

**TF** = NR3C1

**Perturbation condition** = Dexamethasone (A549)

**Off-target perturbation condition** = NA

**Cell type** = A549

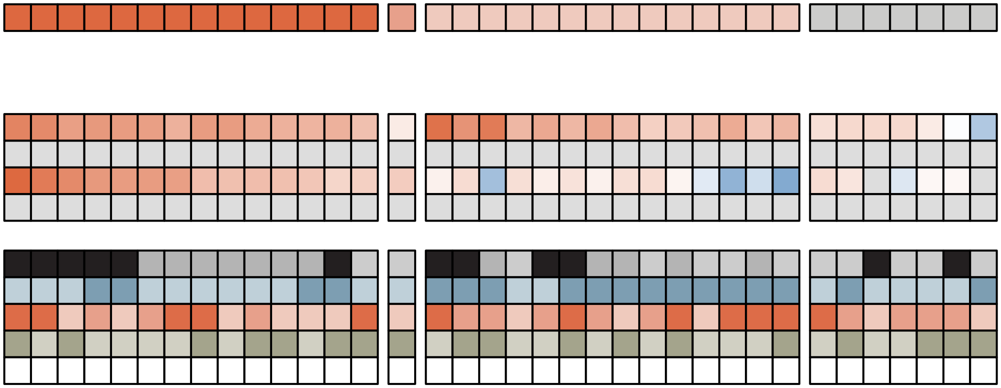

**TF** = NR3C2

**Perturbation condition** = NA

**Off-target perturbation condition** = NA

**Cell type** = mESC

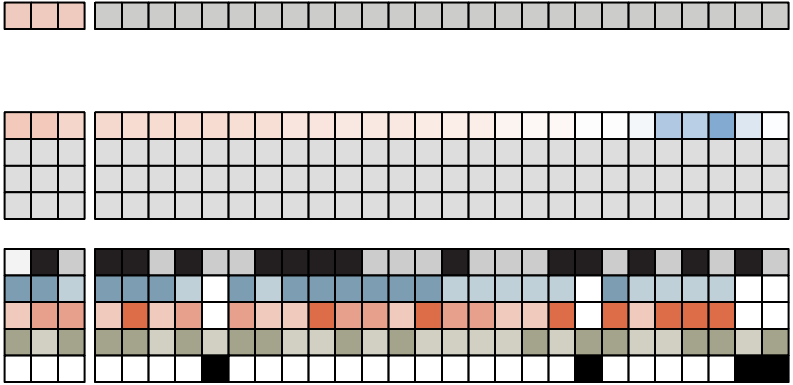

**TF** = NR4A1

**Perturbation condition** = Calcimycin (HCT116) **Off-target perturbation condition** = CDCA (HEPG2)

**Cell type** = mESC

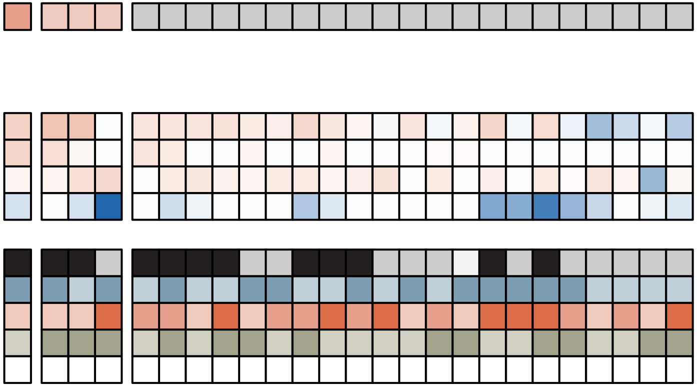

**TF** = NR4A2

**Perturbation condition** = NR4A2 overexpression + cDIM12 (mESC)

**Off-target perturbation condition** = CDCA (HEPG2)

**Cell type** = mESC

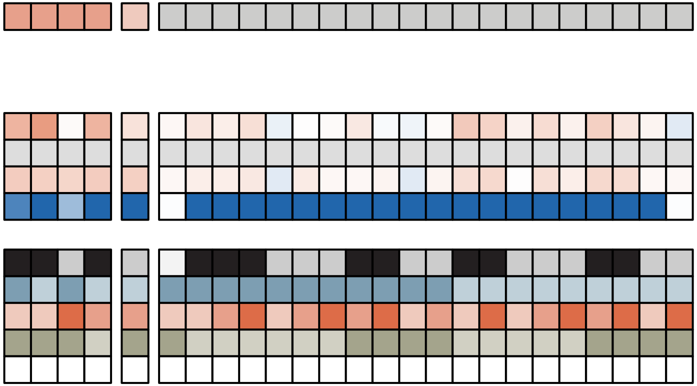

**TF** = NR5A2

**Perturbation condition** = LIF removal (mESC)

**Off-target perturbation condition** = Hexestrol (MCF7)

**Cell type** = mESC

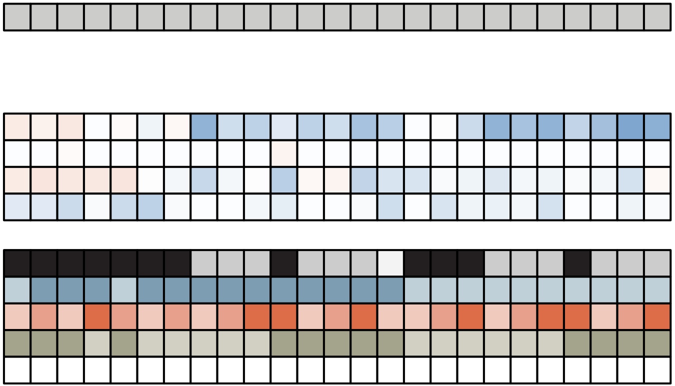

**TF** = NRF1

**Perturbation condition** = NRF1-KD (HEPG2)

**Off-target perturbation condition** = NA

**Cell type** = mESC

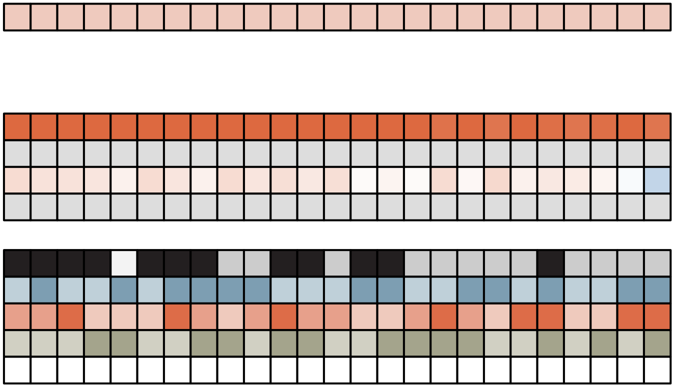

**TF** = ONECUT1

**Perturbation condition** = NA

**Off-target perturbation condition** = NA

**Cell type** = HEPG2

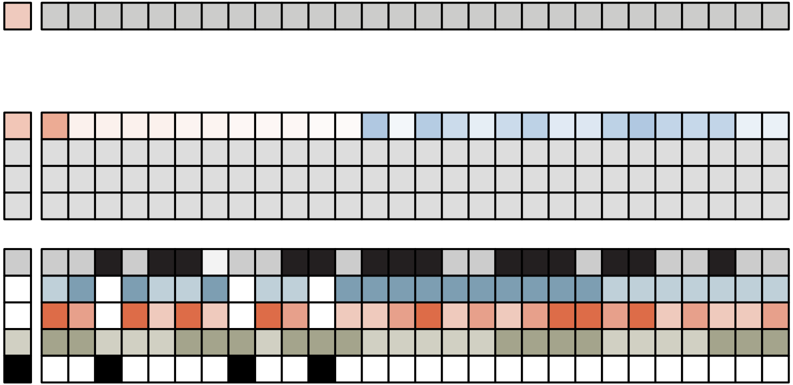

TF = OTX1

Perturbation Condition = mES_IWP2

Off target perturbation condition = NA

Cell type = HepG2

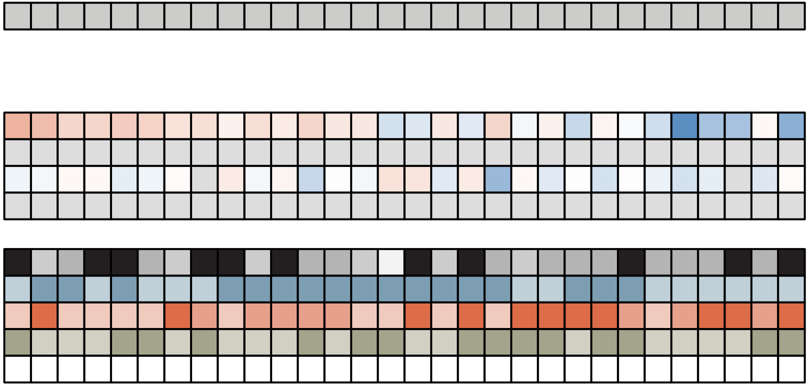

**TF =** PAX6

**Perturbation condition =** PAX6-KD (HEPG2)

**Off-target perturbation condition =** NA

**Cell type =** HEPG2

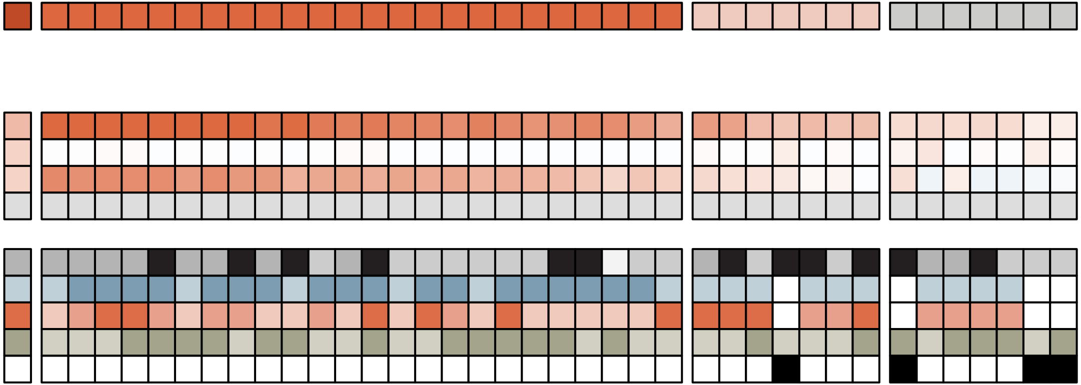

**TF** = PGR

**Perturbation condition** = NA

**Off-target perturbation condition** = NA

**Cell type** = MCF7

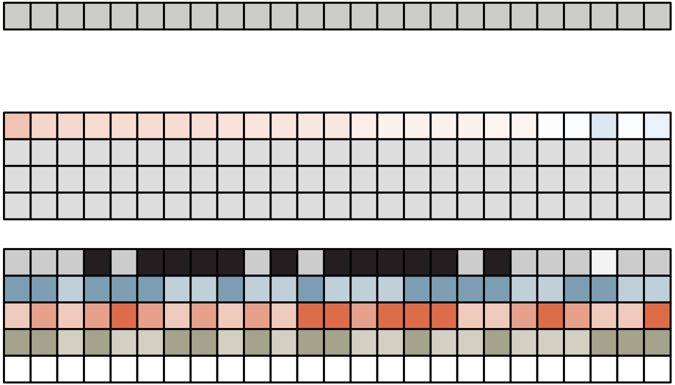

**TF** = POU2F1

**Perturbation condition** = POU2F1-KD (mESC)

**Off-target perturbation condition** = POU5F1-DEG (mESC)

**Cell type** = mESC

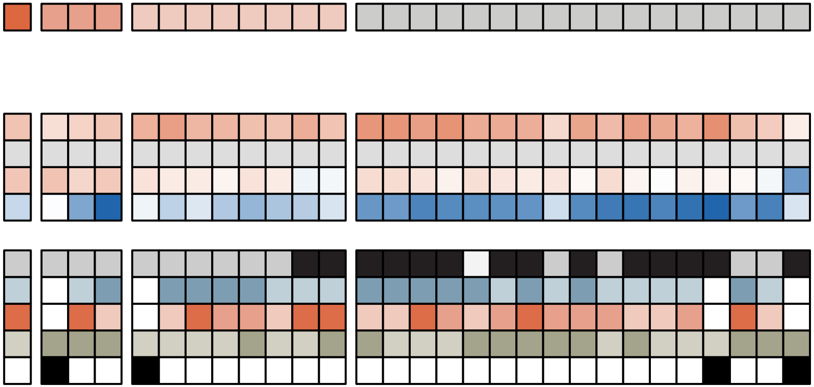

**TF** = POU5F1

**Perturbation condition** = POU5F1-DEG (mESC)

**Off-target perturbation condition** = POU2F1-KD (mESC)

**Cell type** = mESC

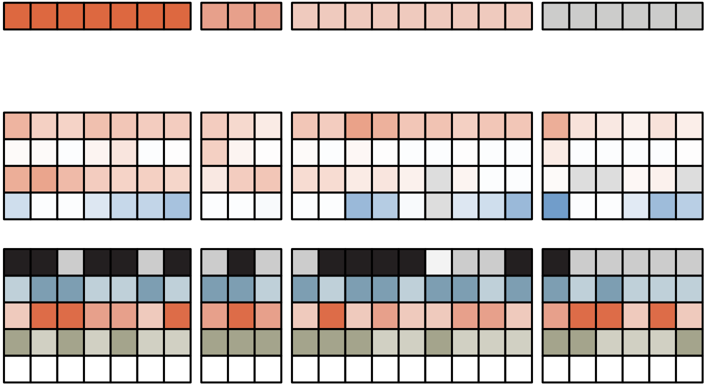

**TF** = POU5F1::SOX2

**Perturbation condition** = POU5F1-DEG (mESC)

**Off-target perturbation condition** = POU2F1-KD (mESC)

**Cell type** = mESC

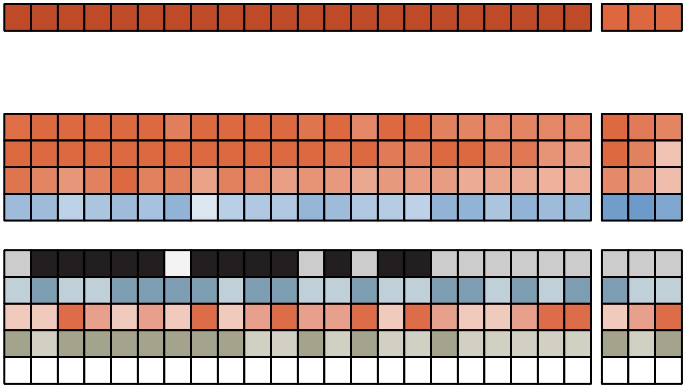

**TF** = PPARA

**Perturbation condition** = NA

**Off-target perturbation condition** = Rifampicin (HEPG2)

**Cell type** = U2OS

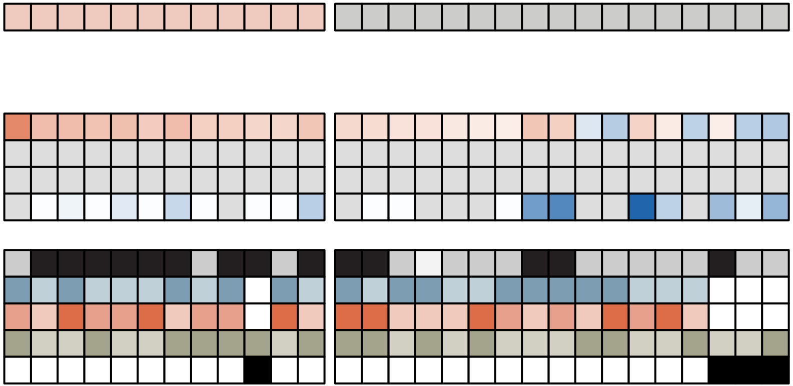

**TF** = PPARG

**Perturbation condition** = NA

**Off-target perturbation condition** = NR1H2-KD (mESC)

**Cell type** = A549

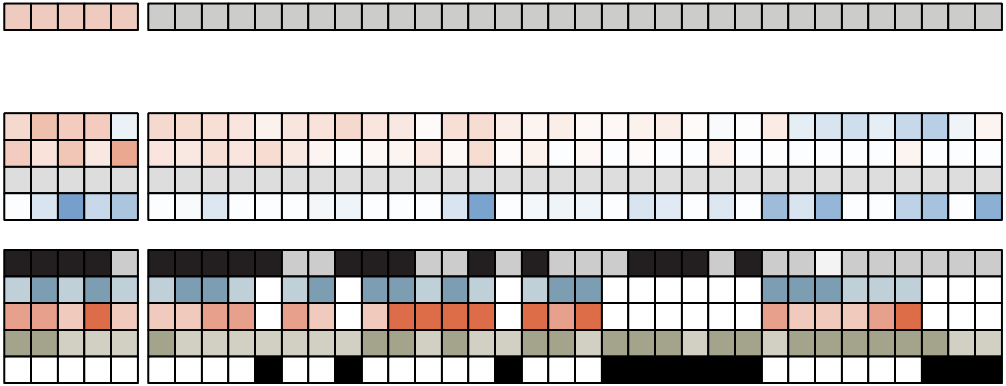

**TF** = RARA

**Perturbation condition** = Basal medium (mESC) **Off-target perturbation condition** = Hexestrol (MCF7)

**Cell type** = U2OS

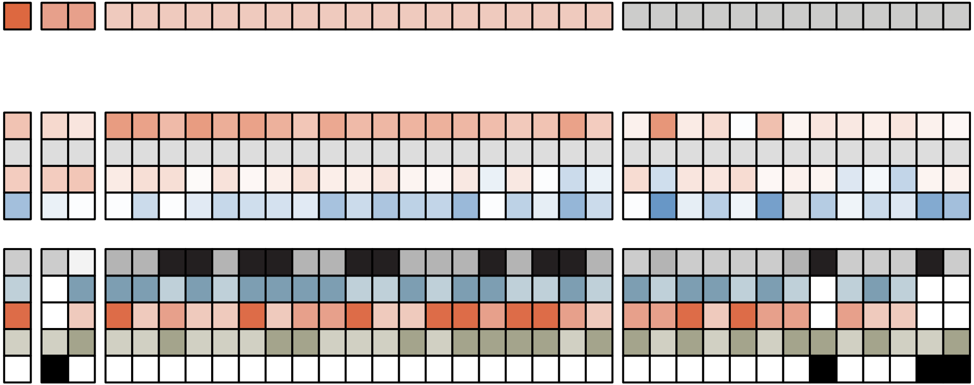

**TF** = RARA::RXRA

**Perturbation condition** = Basal medium (mESC)

**Off-target perturbation condition** = NR4A2 overexpression (mESC)

**Cell type** = mESC

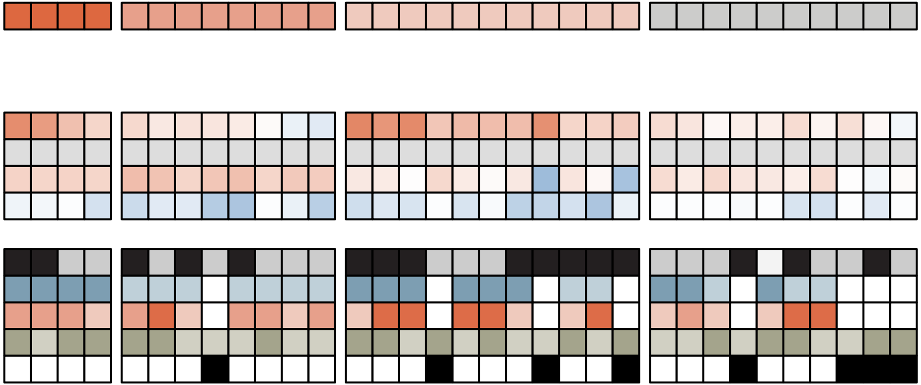

**TF** = RBPJ

**Perturbation condition** = NA

**Off-target perturbation condition** = NA

**Cell type** = HEK293

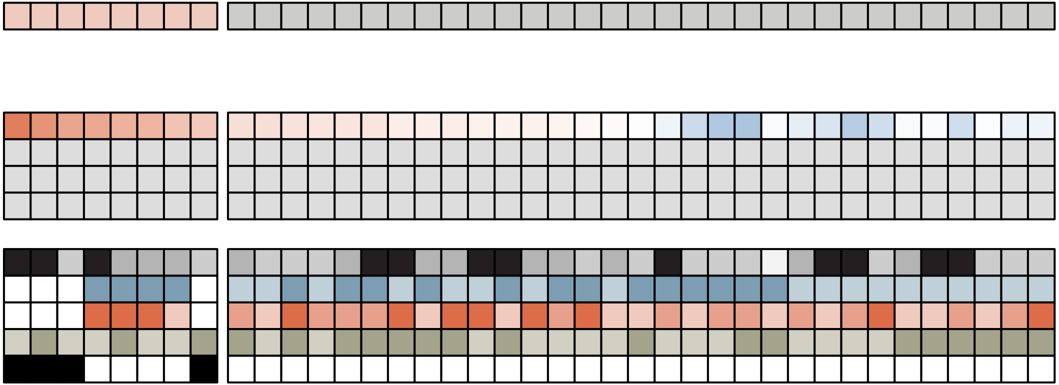

**TF** = REL

**Perturbation condition** = PMA (U2OS)

**Off-target perturbation condition** = NA

**Cell type** = U2OS

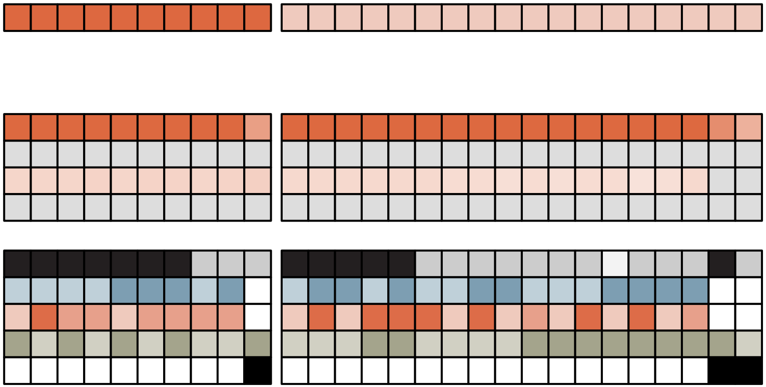

**TF** = RFX1

**Perturbation condition** = NA

**Off-target perturbation condition** = NA

**Cell type** = mESC

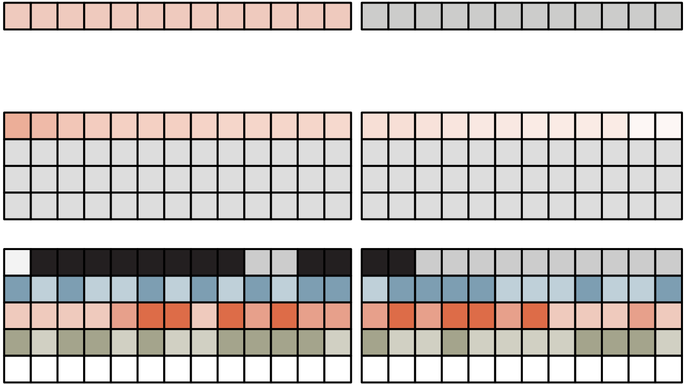

**TF** = RORA

**Perturbation condition** = SR1078 (U2OS)

**Off-target perturbation condition** = Hexestrol (MCF7)

**Cell type** = U2OS

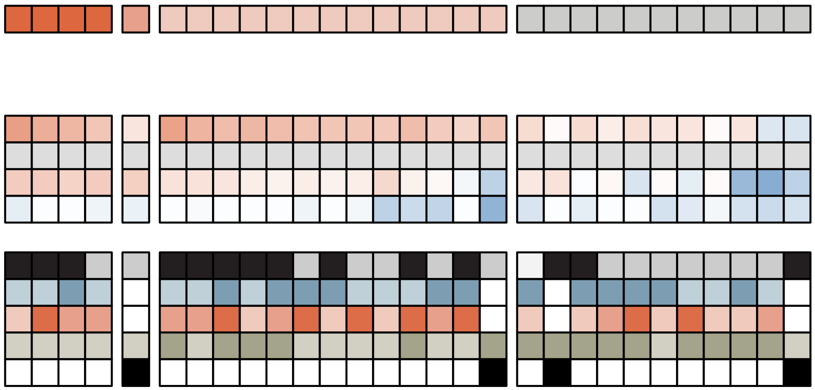

**TF** = RUNX2

**Perturbation condition** = NA

**Off-target perturbation condition** = NA

**Cell type** = U2OS

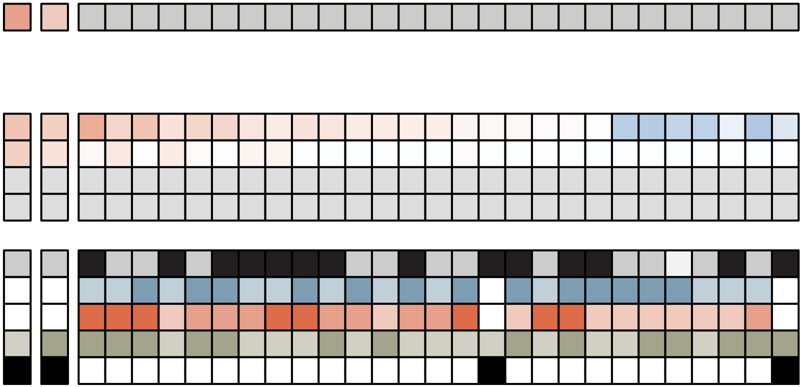

**TF** = RXRA

**Perturbation condition** = Basal medium (mESC)

**Off-target perturbation condition** = NR1H2-KD (mESC)

**Cell type** = mESC

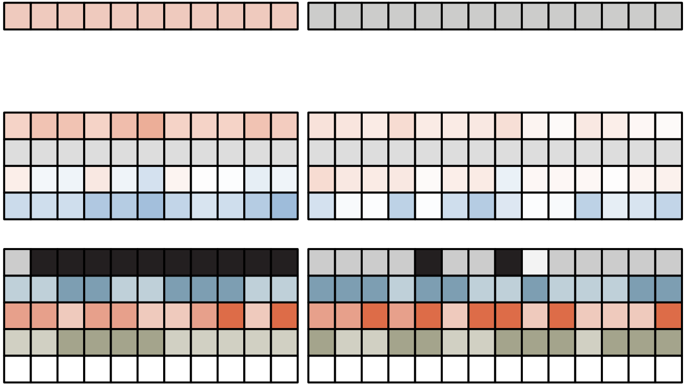

**TF** = SMAD2::3::4

**Perturbation condition** = Serum (mESC)

**Off-target perturbation condition** = NA

**Cell type** = mESC

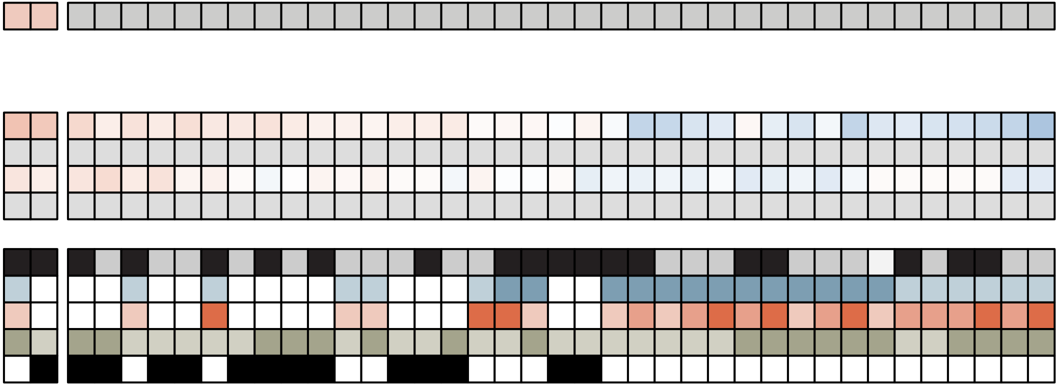

**TF** = SMAD4

**Perturbation condition** = Serum (mESC)

**Off-target perturbation condition** = NA

**Cell type** = mESC

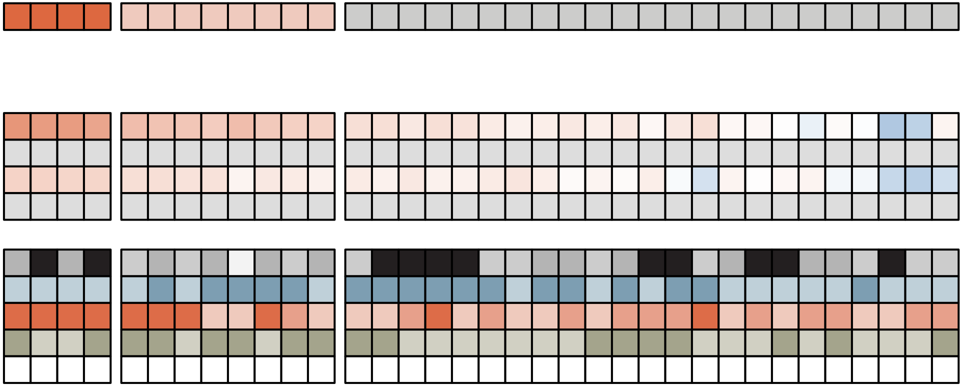

**TF** = SOX2

**Perturbation condition** = SOX2-DEG (mESC)

**Off-target perturbation condition** = SOX13-KD (mESC)

**Cell type** = mESC

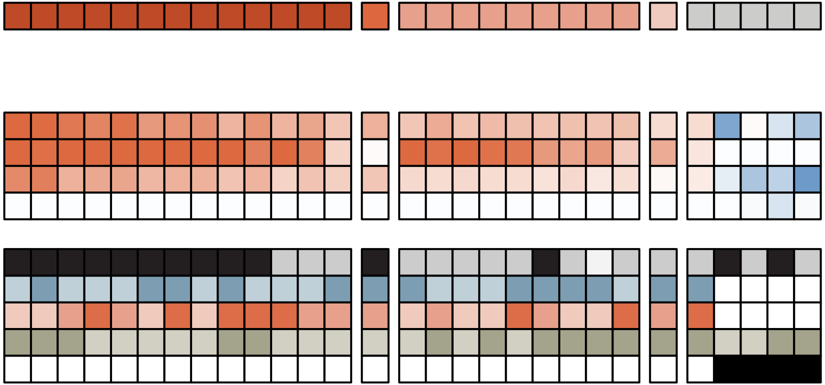

**TF** = SOX9

**Perturbation condition** = NA

**Off-target perturbation condition** = SOX13-KD (mESC)

**Cell type** = HEPG2

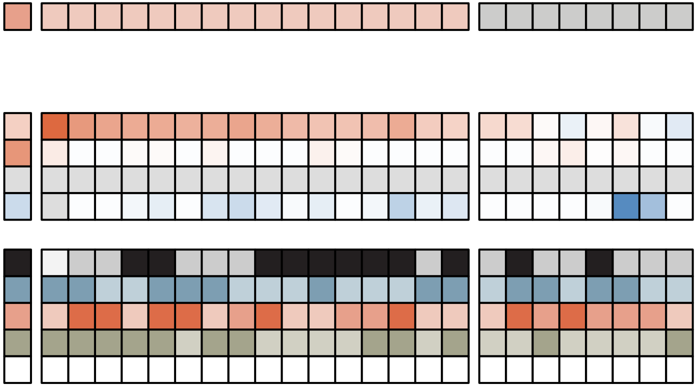

TF = SP1

Perturbation Condition = mES_SP1

Off target perturbation condition = mES_LIF_CH

Cell type = mES

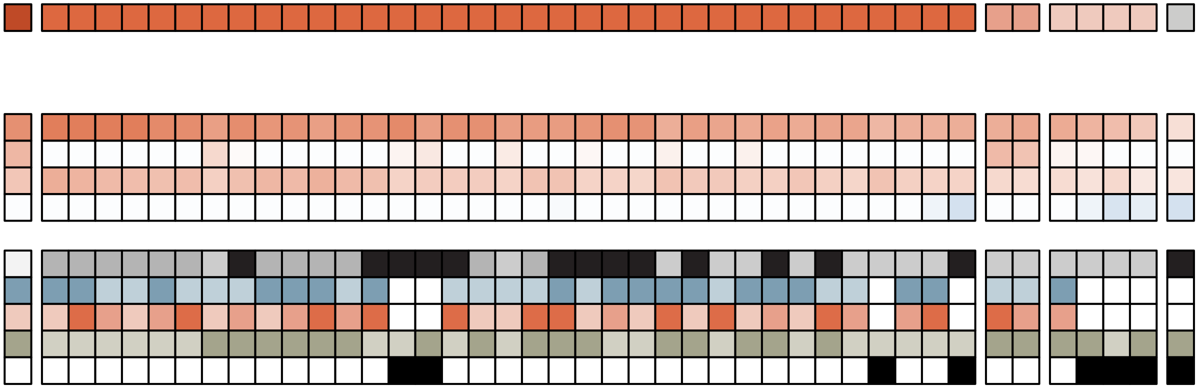

**TF** = SRF

**Perturbation condition** = SRF-KD (HEPG2)

**Off-target perturbation condition** = NA

**Cell type** = mESC

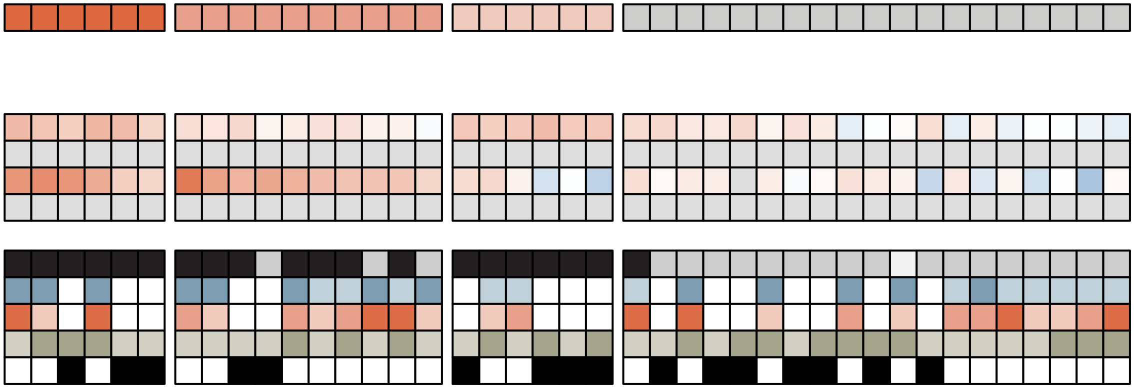

**TF** = STAT1::2

**Perturbation condition** = STAT1-KD (HEPG2)

**Off-target perturbation condition** = Calcimycin (HCT116) **Cell type** = U2OS

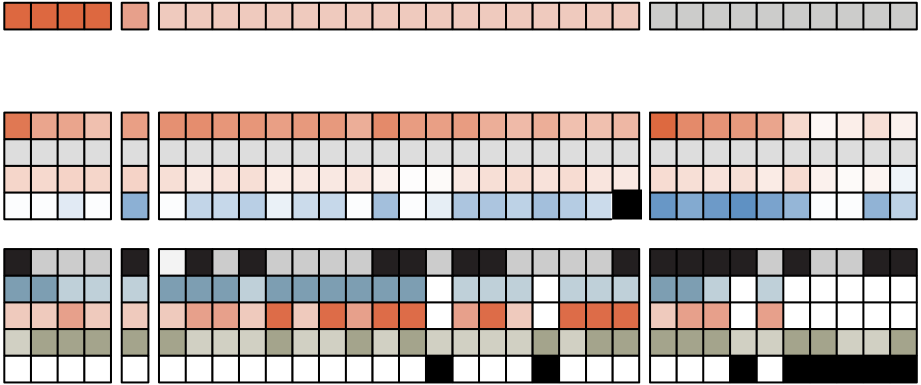

**TF** = STAT3

**Perturbation condition** = LIF removal (mESC)

**Off-target perturbation condition** = STAT1-KD (HEPG2)

**Cell type** = mESC

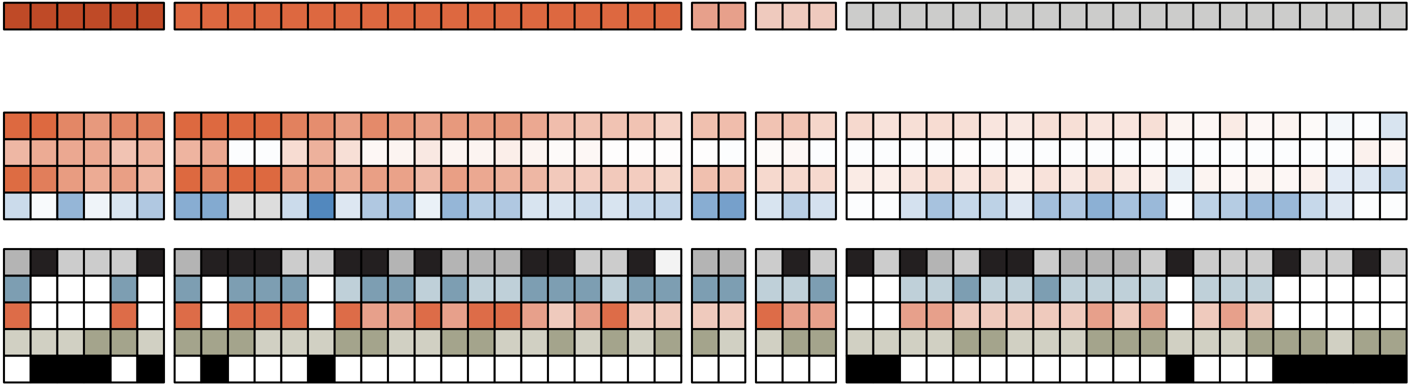

**TF** = TCF7

**Perturbation condition** = CH removal (mESC)

**Off-target perturbation condition** = TCF7L2-KD (mESC)

**Cell type** = mESC

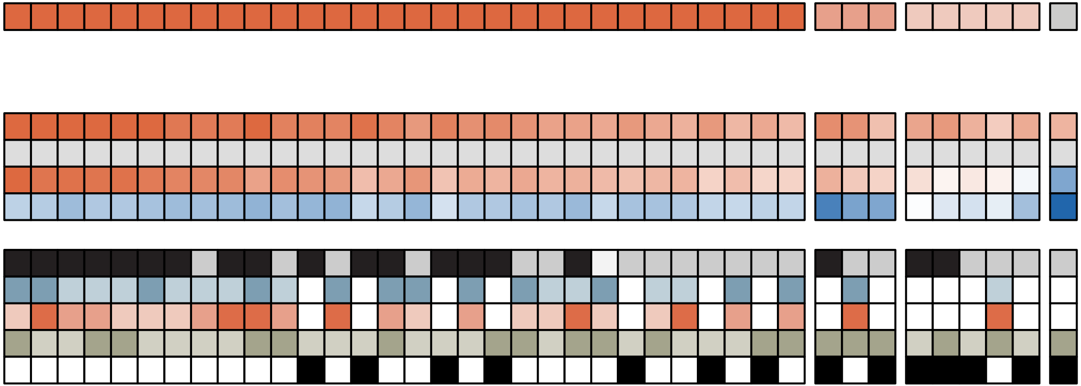

**TF** = TCF7L2

**Perturbation condition** = CH removal (mESC)

**Off-target perturbation condition** = TCF7-KD (mESC)

**Cell type** = HCT116

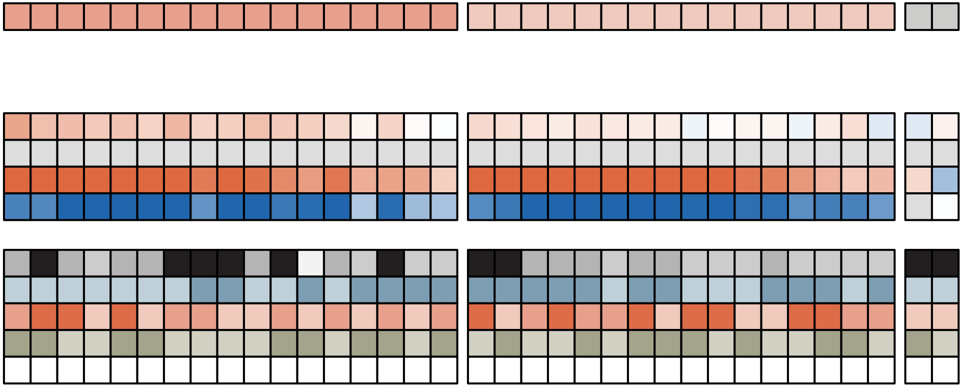

**TF** = TEAD1

**Perturbation condition** = TEAD1-KD (mESC)

**Off-target perturbation condition** = TEAD2-KD (mESC)

**Cell type** = mESC

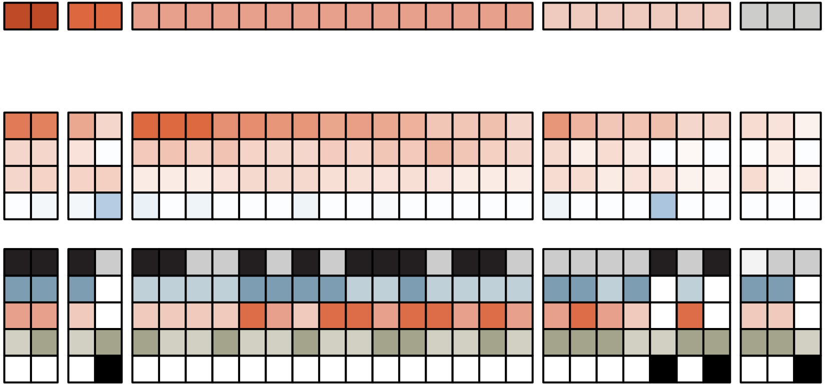

**TF** = TFAP2A

**Perturbation condition** = NA

**Off-target perturbation condition** = NA

**Cell type** = MCF7

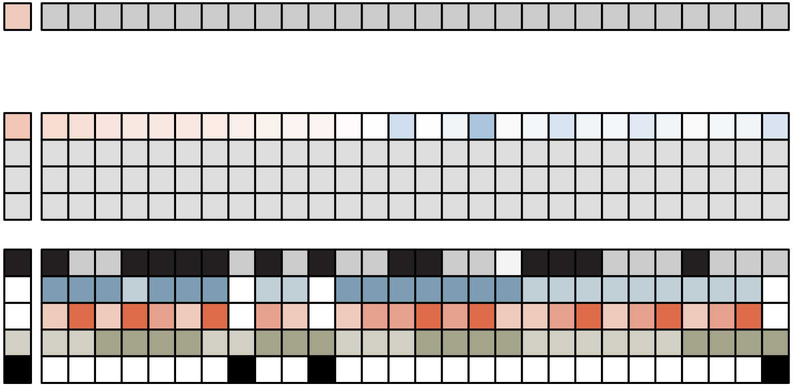

**TF** = TFCP2L1

**Perturbation condition** = TFCP2L1-KD (mESC)

**Off-target perturbation condition** = GRHL1-KD (mESC)

**Cell type** = mESC

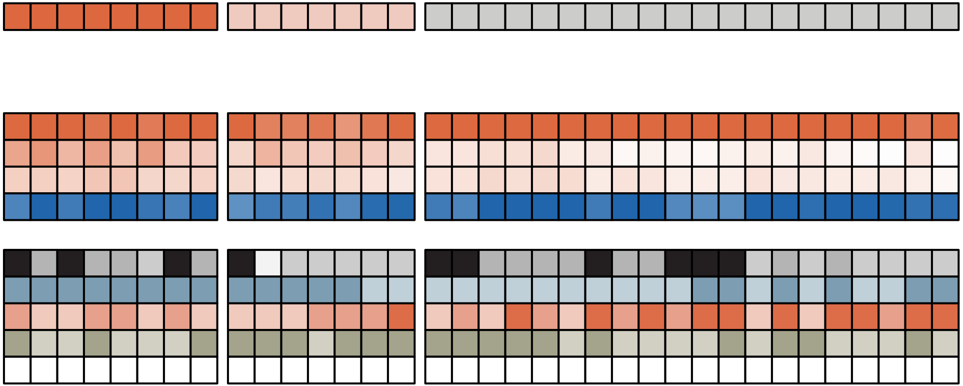

**TF** = THRA

**Perturbation condition** = THRA-KD (HEPG2)

**Off-target perturbation condition** = Hexestrol (MCF7)

**Cell type** = HEK293

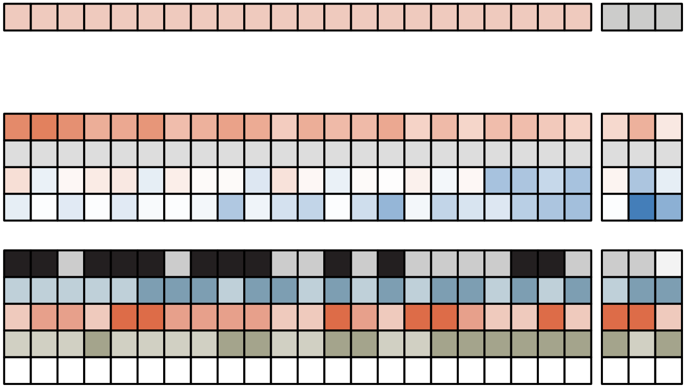

**TF** = THRB

**Perturbation condition** = THRB-KD (HEPG2)

**Off-target perturbation condition** = Hexestrol (MCF7)

**Cell type** = HEK293

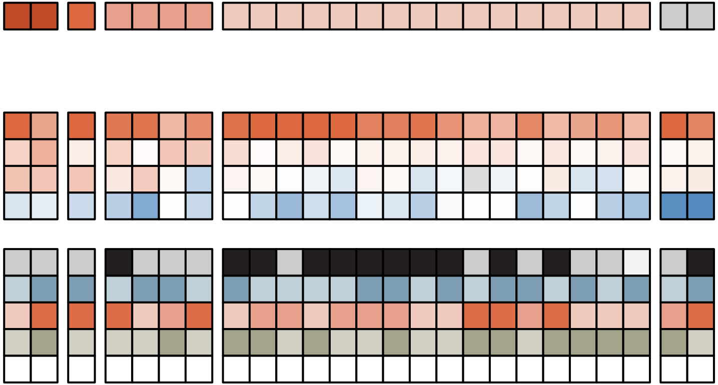

**TF** = TP53

**Perturbation condition** = Nutlin-3a (A549)

**Off-target perturbation condition** = IRX3-KD (HEPG2)

**Cell type** = A549

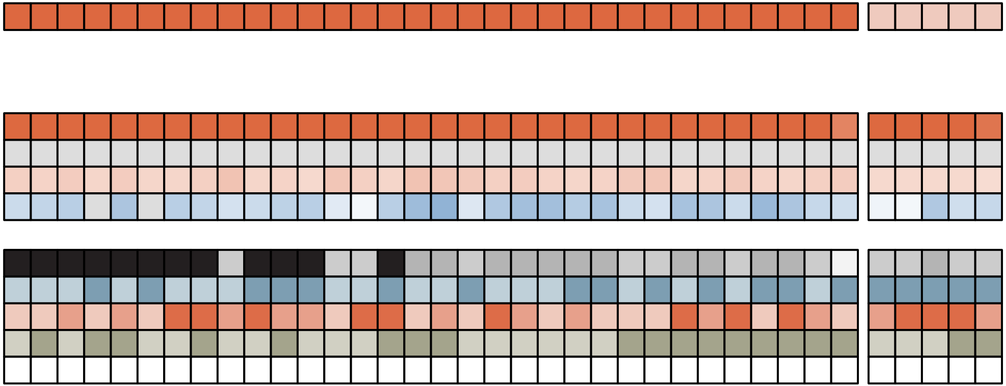

**TF** = VDR

**Perturbation condition** = Calcitriol (U2OS)

**Off-target perturbation condition** = Rifampicin (HEPG2)

**Cell type** = U2OS

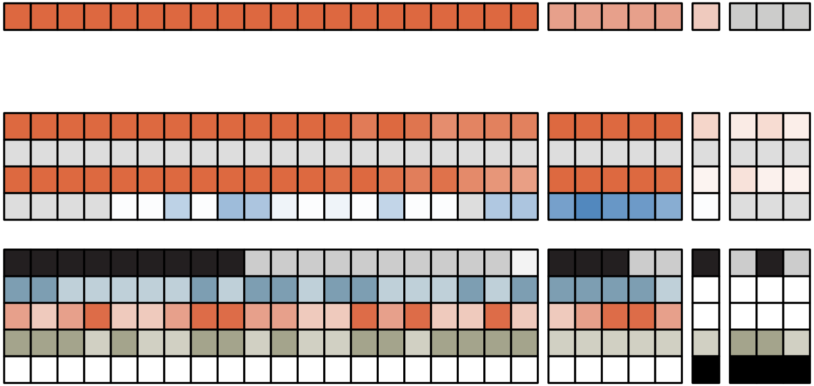

**TF** = WT1

**Perturbation condition** = NA

**Off-target perturbation condition** = PD removal (mESC)

**Cell type** = U2OS

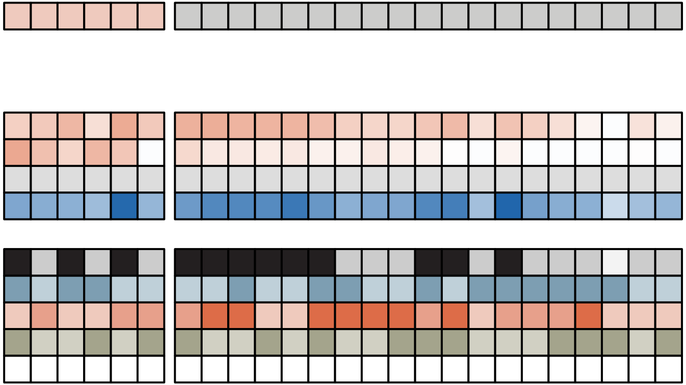

TF = XBP1

Perturbation condition = XBP1-KD (HEPG2)

Off-target perturbation condition = ATF6-KD (HEPG2)

Cell type = MCF7

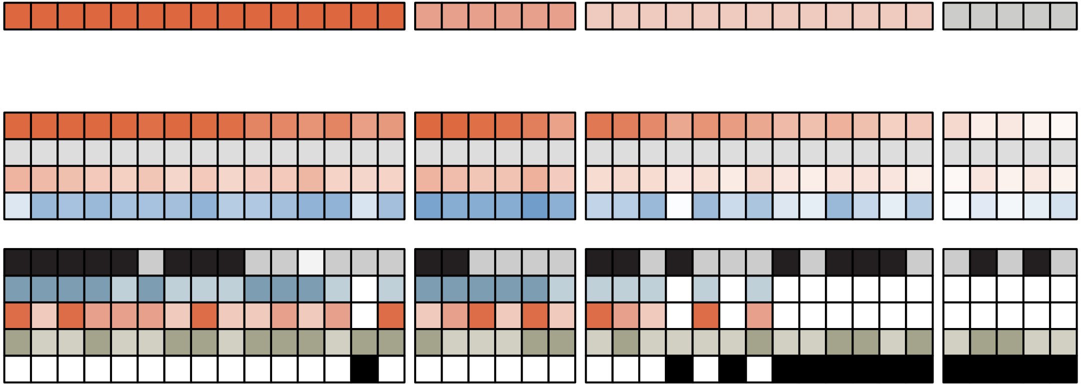

**TF** = ZFP42

**Perturbation condition** = NA

**Off-target perturbation condition** = NA

**Cell type** = mESC

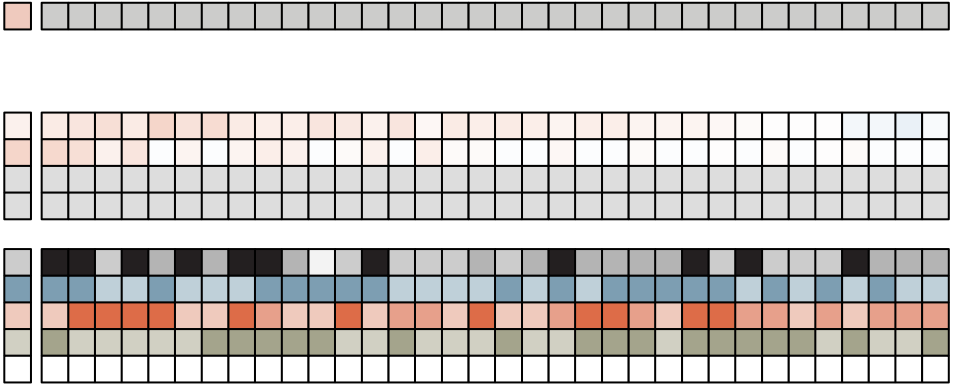

**TF** = ZFX

**Perturbation condition** = NA

**Off-target perturbation condition** = NA

**Cell type** = mESC

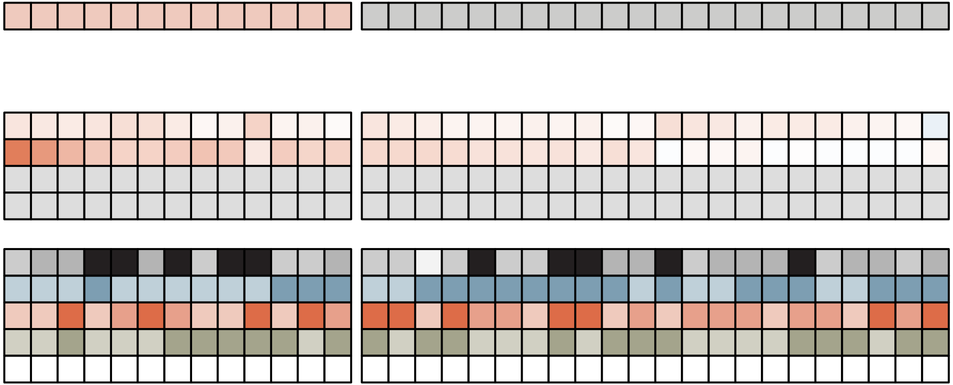
